# Weighting by Gene Tree Uncertainty Improves Accuracy of Quartet-based Species Trees

**DOI:** 10.1101/2022.02.19.481132

**Authors:** Chao Zhang, Siavash Mirarab

**Affiliations:** Bioinformatics and Systems Biology, UC San Diego, CA, USA; Department of Electrical and Computer Engineering, UC San Diego, CA, USA

**Keywords:** Phylogenomics, ILS, Summary methods, ASTRAL, gene tree estimation error

## Abstract

Phylogenomic analyses routinely estimate species trees using methods that account for gene tree discordance. However, the most scalable species tree inference methods, which summarize independently inferred gene trees to obtain a species tree, are sensitive to hard-to-avoid errors introduced in the gene tree estimation step. This dilemma has created much debate on the merits of concatenation versus summary methods and practical obstacles to using summary methods more widely and to the exclusion of concatenation. The most successful attempt at making summary methods resilient to noisy gene trees has been contracting low support branches from the gene trees. Unfortunately, this approach requires arbitrary thresholds and poses new challenges. Here, we introduce threshold-free weighting schemes for the quartet-based species tree inference, the metric used in the popular method ASTRAL. By reducing the impact of quartets with low support or long terminal branches (or both), weighting provides stronger theoretical guarantees and better empirical performance than the original ASTRAL. More consequentially, weighting dramatically improves accuracy in a wide range of simulations and reduces the gap with concatenation in conditions with low gene tree discordance and high noise. On empirical data, weighting improves congruence with concatenation and increases support. Together, our results show that weighting, enabled by a new optimization algorithm we introduce, dramatically improves the utility of summary methods and can reduce the incongruence often observed across analytical pipelines.

## Introduction

Genome-wide data are increasingly available across the tree of life, giving researchers a chance to systematically resolve the evolutionary relationships among species (i.e., species trees) using phylogenomic data. A central promise of phylogenomics is that processes such as incomplete lineage sorting (ILS) that can cause discordance (Degnan and Rosenberg, 2009; Maddison, 1997) among evolutionary histories of different parts of the genome (i.e., gene trees) can be modeled (Edwards, 2009). There has been much progress in developing the theory and methods for species tree inference in the presence of ILS (Mirarab et al., 2021) and other sources of discordance (Elworth et al., 2019; Smith and Hahn, 2021). These phylogenomics approaches have also been widely and increasingly adopted in practice. Yet, substantial challenges remain. Analyses of real data using different methods often reveal incongruent results (Gatesy et al., 2019; Reddy et al., 2017; Shen et al., 2017; Smith et al., 2015; Walker et al., 2018), sparking debate about the cause. Meanwhile, simulation studies have revealed that the best choice of the method is data-dependent (e.g., Bayzid and Warnow, 2013; Mirarab and Warnow, 2015).

A major challenge in phylogenomics is that when we infer gene trees, often from relatively short sequences, the results tend to be highly error-prone (Mirarab et al., 2014a; Patel, 2013; Springer and Gatesy, 2016). Co-estimation of gene trees and species trees (Szöllõsi et al., 2014) is perhaps the most accurate approach to dealing with such noise (Knowles et al., 2012; Leaché and Rannala, 2011). However, despite some progress (Ogilvie et al., 2017), these methods have remained limited in their scalability to even moderately large numbers of species. The approach that is far more scalable and is used often is the “summary” approach: first estimate gene trees from sequence data independently and then summarize them into a species tree by solving optimization problems that provide guarantees of statistical consistency if we allow ourselves to ignore the error in the input tree.

Many summary methods (e.g., Liu and Yu, 2011; Liu et al., 2010, 2009; Mossel and Roch, 2010; Vachaspati and Warnow, 2015) were developed and proved statistically consistent under the multi-species coalescent (MSC) model (Takahata, 1989) of the discordance caused by ILS. Species trees inferred by these tools can be highly accurate even under high levels of ILS. Among the summary tools, ASTRAL (Mirarab et al., 2014b) is among the most widely used and is integrated into other packages (Alanjary et al., 2019; Wang et al., 2020). ASTRAL simply seeks the species tree that maximizes the number of shared quartets (unrooted four-taxon subtrees) between gene trees and the species tree, an optimization problem that guarantees a statistically consistent estimator under the MSC model. The empirical accuracy and scalability of ASTRAL have compared favorably to other methods (e.g., Mirarab, 2019). Moreover, it has now been shown that ASTRAL is also consistent and/or accurate under the gene duplication and loss (GDL) model (Legried et al., 2021; Yan et al., 2021), some horizontal gene transfer models (Davidson et al., 2015), and combined models of ILS and GDL (Markin and Eulenstein, 2021), but not gene flow (Solís-Lemus et al., 2016). Zhang et al. 2020 have further adopted the quartet-based approach to multi-copy inputs.

Nevertheless, all summary methods, ASTRAL included, have a shortcoming: inaccuracies in input gene trees can translate to errors in the output species tree (DeGiorgio and Degnan, 2014; Huang and Knowles, 2016; Lanier and Knowles, 2015; Molloy and Warnow, 2018; Patel, 2013). In fact, Roch et al. 2019 proved that summary methods (and concatenation) are positively misleading under pathological examples even in the absence of much true gene tree discordance. These concerns are not just theoretical and can impact biological analyses. For example, on an order-level avian phylogenomic dataset (Jarvis et al., 2014), summary methods, including ASTRAL, produce species trees contradicting the well-established relationships when given input gene trees that have extremely low support (Bayzid et al., 2015), a condition that motivated Mirarab et al. 2014a to bin multiple genes together. As an alternative, Zhang et al. 2018 showed that contracting very low-support branches before running ASTRAL can improve accuracy in simulations and on biological datasets such as the avian dataset. However, this form of reduction in species tree estimation error comes with caveats. Contracted branches may still include signals that will be lost. In particular, when contraction is overly aggressive (e.g., with moderately high thresholds such as 50% or 75%), filtering is often harmful. More pragmatically, the best choice of threshold is dataset dependent, and making a principled choice is challenging if not impossible.

Threshold-free approaches for incorporating gene tree branch support into summary methods have also been proposed. Multi-locus bootstrapping (MLBS) runs the summary method on the bootstrap replicates of gene trees, repeating the process many times to obtain several species trees, which are then combined using a consensus method (Seo, 2008). MLBS can be understood as weighting inferences made from each gene by their uncertainty, and thus, a way to deal with noise. However, previous studies show that MLBS, in fact, reduces the accuracy compared to using Maximum Likelihood (ML) trees (Mirarab et al., 2016). The related method of simply combining all bootstrap replicates into a single run of the summary method has also not been accurate (Mirarab et al., 2014b). A plausible explanation is that bootstrap replicates have much higher rates of discordance and error than ML trees (Sayyari and Mirarab, 2016), and thus, using them directly as input adds noise, even if it reveals uncertainty.

An alternative to using bootstrap trees is to use ML trees as input but explicitly weight gene tree branches (or their quartets) by their statistical support. We can generalize the moderately successful gene contraction approach, which effectively assigns weights zero or one to quartets, to weight each quartet shared between an estimated gene tree and the proposed species tree according to the statistical support of the quartet resolution. Such an approach will free us from picking arbitrary contraction thresholds and may lead to better accuracy. However, weighting by branch support has not yet been incorporated into existing summary methods such as ASTRAL for several reasons. *i)* Quartet weights must be implicitly calculated, as explicitly examining all quartets of *n* species alone will take Θ(*n*^4^) time. The existing general (e.g., Avni et al., 2015) and MSC-based weighted quartet methods (Richards and Kubatko, 2021; Yourdkhani and Rhodes, 2020) require weights *explicitly* calculated for every quartet, making them less scalable with *n*. The reason ASTRAL can scale to a large number of species is that it optimizes a score defined over all quartets without explicitly examining them. Designing a scalable weighting method will require weights that can be implicitly computed based on examining *O*(*n*) gene tree branches. ii) It is difficult to design efficient algorithms to optimize a weighted score. Unless weights satisfy certain properties, it may not be possible to find an algorithm better than *O*(*n*^4^) even for the much simpler problem of computing the total quartet weights of a gene tree. However, with favorable definitions of weights, these difficulties are not insurmountable.

Here, we introduce implicit weighting schemes that avail themselves to efficient optimization with weights conveniently obtained from tree branch lengths (wASTRAL-bl), branch support values (wASTRAL-s), or both (wASTRAL-h). We introduce the weighted ASTRAL algorithm, an efficient method that is similar to ASTRAL in optimizing a quartet score but is different in several ways: *i*) Its optimization criteria weights each gene tree quartet. *ii*) Its optimization algorithm is entirely different from ASTRAL. While the algorithm is more complex and slower in some cases, it scales much better (linearly instead of quadratically) as the number of genes (*k*) increases. *iii*) Its software package is implemented from scratch and is in C++ instead of Java. Our results show that weighted ASTRAL is superior to ASTRAL in terms of theoretical guarantees that it provides, accuracy on simulated data, and the accuracy of its branch support values. Weighted ASTRAL is more accurate than CA-ML in our simulations except when there is a large number of inaccurate gene trees or low levels of discordance, where concatenation is slightly more accurate. Most interestingly, weighted ASTRAL is more congruent than the original ASTRAL with concatenation on real datasets.

## Result

### Weighted ASTRAL algorithm

Unlike ASTRAL-III, where each (resolved) quartet in each gene tree contributes equally to the objective function, weighted ASTRAL assigns each quartet with a weight based on the support or lengths of branches corresponding to it. More specifically, we define three weighting schemes (Fig. 1a).

**FIG. 1.**
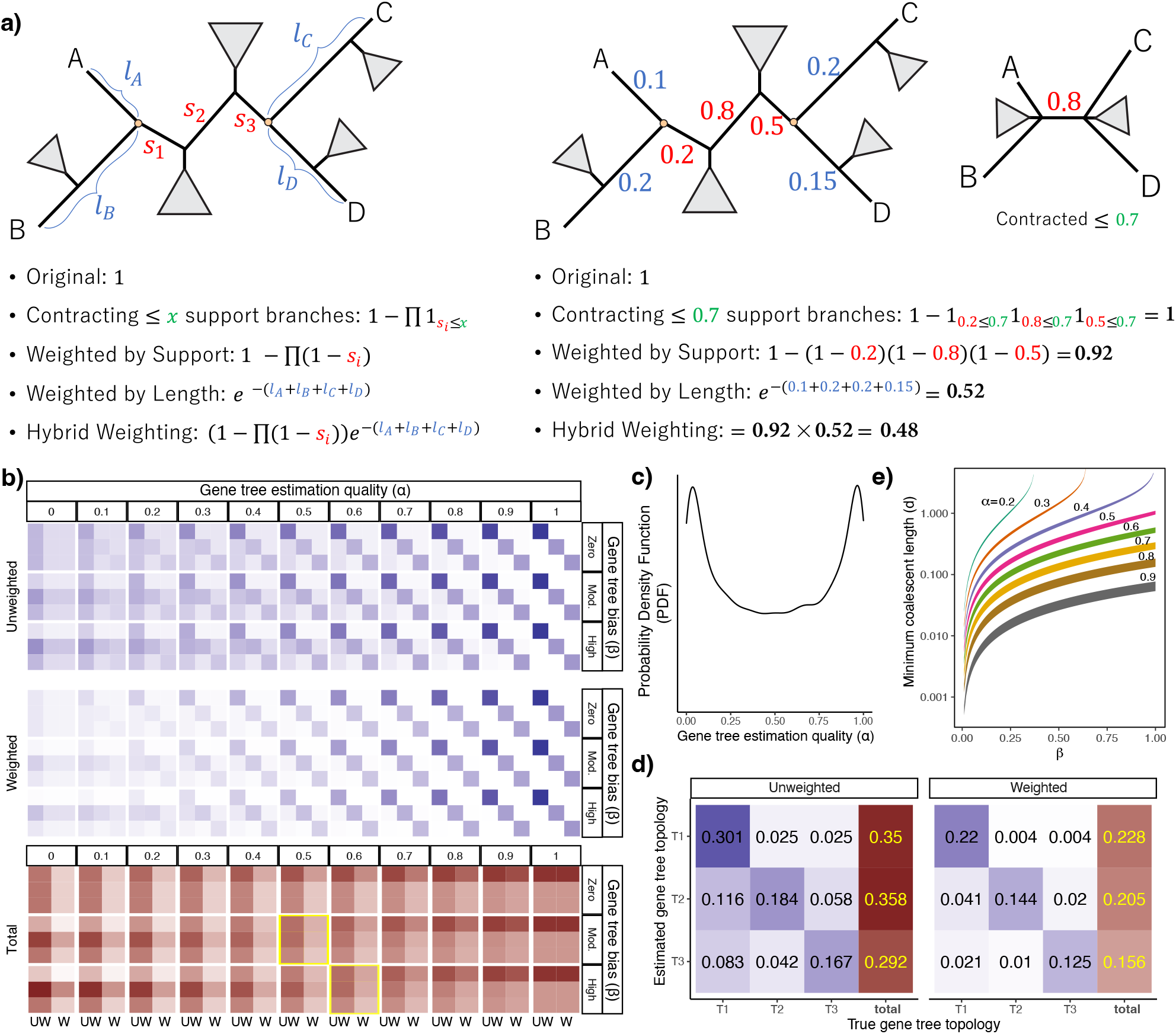
(a) Illustration of weighting methods. The generic formula and an example of weighting gene tree quartet *ab*|*cd*. Trees are annotated with the support (red) of all branches between anchors (orange dots) and the substitution per site unit length of each leaf-to-anchor path (blue). (b-e) Illustration of the impact of weighting using our MSC+Error+Support model for a quartet species tree with internal branch length set to — ln0.75 CU. (b) Top: each 3 × 3 square shows the joint probabilities of true (by column) and estimated (by row) gene trees for each of the three possible quartet topologies. The first row/column represents the topology matching species tree, and the second row/column corresponds to the topology towards which gene tree estimation is biased. The gene tree estimation quality *α* ranges in [0,1], and the bias in gene tree estimation *β* is set to zero, moderate (0.4), or high (0.6). These probabilities correspond to expected weights in normal ASTRAL. Middle: The expected weights in wASTRAL-s for each scenario. Bottom: The 3 × 2 grids show the marginal expected score of each topology (rows) for unweighted ASTRAL (UW; first column) and weighted ASTRAL (W; second column). Note the reduced darkness of W columns as *α* decreases. The two girds highlighted in yellow: the score is highest for the wrong (second row) topology without weights but is higher for the correct topology (first row) with weights. (c) Distribution *α* drawn from *Beta*(0.5,0.5) across genes in a toy example. (d) Joint (blue) and marginal (red) probabilities of topologies with and without weighting with moderate bias (*β* = 0.4) and *α* drawn from the distribution shown on top. (e) Each band shows the range of coalescent unit (CU) quartet internal branch length where ASTRAL is not consistent but support weighted ASTRAL is, for different *α* and *β* valeus.

*Weighting by support* extends the definition of branch support to a quartet. Let 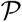 be the set of branches on the path between internal nodes of a quartet tree (also called anchors; orange dots in Fig. 1a) and let *s*(*e*) denote the support of a branch *e*. We define the support of the quartet as

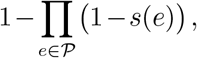

which essentially assumes support values are probabilities of correctness and that branches are independent (both assumptions can be disputed). Given a set of gene trees where each internal branch has a support value, using this definition, we define the weight of each quartet of each gene tree to be its support. The goal is to improve the accuracy by down-weighting quartets with low support. While we study this goal in our simulation and empirical analyses, we also provide some theoretical results.

Making theoretical statements about estimated gene trees is difficult because we lack an accepted way of modeling gene tree estimation errors. To be able to interrogate theoretical properties of weighted ASTRAL, we propose a simple model of gene tree estimation error called MSC+Error (Material and Methods). In this model, for any true gene tree topology on a quartet *Q*, the estimated topology is drawn from a distribution that has two features: first, each gene *G* has a gene-specific level of signal, controlled by a parameter *α_G,Q_*, and second, all genes can be adversarially biased towards any topology by an amount bounded by a parameter called *β_Q_*. The joint distribution of true and estimated quartet gene trees in the most difficult case can be expressed as a function of *α_G,Q_* and *β_Q_* as well as *θ_Q_* = 1 – *e^-d^* where *d* is the coalescent unit (CU) length of the internal branch of the quartet (Table 1 and Fig. 1b). Under the MSC+Error model, the distribution of quartet gene tree topologies, written as a vector with the first element corresponding to the species tree, changes (in the worst case) from 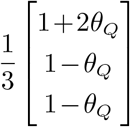 for true gene trees to 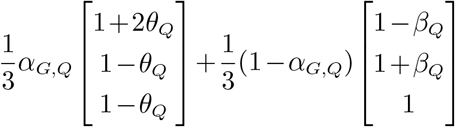 for estimated gene trees.

**Table 1.** Joint probabilities (top) and weights (bottom) of estimated and true gene tree topologies under the MSC+Error+Support with the worst-case scenario when 3*p*_1_ = 1 – β, 3*p*_2_ = 1 + *β*, and 3*p*_3_ = 1 for all genes; note that parameters are per quartet and per gene but we omit *Q* and *G* superscript for brevity.

| $\mathbb{E}[(\cdot)(\cdot)\vert \alpha]$ | $\delta_{G^*}(ab\vert cd)$ | $\delta_{G^*}(ac\vert bd)$ | $\delta_{G^*}(ad\vert bc)$ |
| --- | --- | --- | --- |
| $\delta_G(ab\vert cd)$ | $\frac{1}{3}(1+2\theta)(\alpha + \frac{1}{3}(1-\alpha)(1-\beta))$ | $\frac{1}{3}(1-\theta)(\frac{1}{3}(1-\alpha)(1-\beta))$ | $\frac{1}{3}(1-\theta)(\frac{1}{3}(1-\alpha)(1-\beta))$ |
| $\delta_G(ac\vert bd)$ | $\frac{1}{3}(1+2\theta)(\frac{1}{3}(1-\alpha)(1+\beta))$ | $\frac{1}{3}(1-\theta)(\alpha + \frac{1}{3}(1-\alpha)(1+\beta))$ | $\frac{1}{3}(1-\theta)(\frac{1}{3}(1-\alpha)(1+\beta))$ |
| $\delta_G(ad\vert bc)$ | $\frac{1}{3}(1+2\theta)(\frac{1}{3}(1-\alpha))$ | $\frac{1}{3}(1-\theta)(\frac{1}{3}(1-\alpha))$ | $\frac{1}{3}(1-\theta)(\alpha + \frac{1}{3}(1-\alpha))$ |
| $w_G(ab\vert cd)$ | $\frac{1}{3}(1+2\theta)(\alpha + \frac{1}{3}(1-\alpha)(1-\beta))^2$ | $\frac{1}{3}(1-\theta)(\frac{1}{3}(1-\alpha)(1-\beta))^2$ | $\frac{1}{3}(1-\theta)(\frac{1}{3}(1-\alpha)(1-\beta))^2$ |
| $w_G(ac\vert bd)$ | $\frac{1}{3}(1+2\theta)(\frac{1}{3}(1-\alpha)(1+\beta))^2$ | $\frac{1}{3}(1-\theta)(\alpha + \frac{1}{3}(1-\alpha)(1+\beta))^2$ | $\frac{1}{3}(1-\theta)(\frac{1}{3}(1-\alpha)(1+\beta))^2$ |
| $w_G(ad\vert bc)$ | $\frac{1}{3}(1+2\theta)(\frac{1}{3}(1-\alpha))^2$ | $\frac{1}{3}(1-\theta)(\frac{1}{3}(1-\alpha))^2$ | $\frac{1}{3}(1-\theta)(\alpha + \frac{1}{3}(1-\alpha))^2$ |

The estimated gene tree distribution matches the MSC model when *α_G,Q_* = 1 and is uniformly random when *α_G,Q_* = *β_Q_* =0. A choice of *α_G,Q_* < 1 adds noise to the MSC probabilities, and any *β_Q_* > 0 creates an adversarial bias towards the second topology (Fig. 1b). Because noise and bias parameters can change across genes and quartets, the MSC+Error model is very general and makes minimal assumptions.

Under the MSC+Error model, the original ASTRAL is statistically consistent with estimated gene trees under limited choices of *α_G,Q_* and *β_Q_*. Assuming that the support of a quartet matches the estimated gene tree distribution, we can get our main result. Theorem 1 in Material and Methods proves that support-weighted ASTRAL (wASTRAL-s) is statistically consistent under a strictly larger super-set of *α_G,Q_* and *β_Q_* parameters than those of unweighted ASTRAL. Thus, there are levels of bias in gene tree estimation (e.g., due to long branch attraction) that, combined with low signal, render unweighted ASTRAL inconsistent (as shown by Roch et al. 2019) but keep wASTRAL-s consistent.

Examining the marginal probabilities and expected weights can illuminate the reason behind the advantage of wASTRAL-s (Fig. 1b). First, gene trees with higher levels of noise (i.e., lower *α_G,Q_*) are down-weighted relative to gene trees with less noise (Fig. 1b: note lighted colors as *α* decreases). Thus, the correct topology benefits from summing weights over gene trees with different *α_G,Q_*. For example, assume some genes have high noise, and others have low noise following the *α_G,Q_* distribution shown in Figure 1c. The less noisy genes will be up-weighted such that wASTRAL-s becomes consistent even when unweighted ASTRAL is not (Fig. 1d). Second, unless gene trees are extremely noisy (i.e., very low *α_G,Q_*), wASTRAL-s down-weights the species tree topology less than the other two topologies; in extreme cases, we have scenarios (Fig. 1b, bottom, highlighted boxes) where the species tree is dominant with weighted scores but not with unweighted scores. In fact, for fixed *α* and *β*, there exists a range of CU quartet internal branch lengths for which ASTRAL is not consistent but wASTRAL-s is (Fig. 1e).

*Weighting by length* down-weights quartets with long terminal branches. Let *L* be the sum of terminal branch lengths in the gene tree induced to a quartet provided in substitution-per-site units (SU). We assign *e*^-*L*^ as the weight of the quartet and offer two justifications. First, deeper coalescence events tend to generate longer terminal branch lengths; thus, gene trees that match the species tree are expected, on average, to have shorter branch lengths (see proof of Theorem 2). Thus, down-weighting gene tree quartets with long terminal branches is expected to down-weight genes that do not match the species tree. Doing so can provably provide a bigger gap between the score of the true species tree and alternatives, as shown in Theorem 2. Besides the connection to the MSC model, it has also been long appreciated that the so-called long quartets are harder to estimate correctly due to long branch attraction (Erdos et al., 1999; Snir et al., 2008). Many quartet-based methods focus their attention on the so-called short quartets (Nelesen et al., 2012; Warnow et al., 2001). Our weighting scheme naturally achieves the same impact by down-weighting long quartets versus short quartets around difficult species tree branches (Fig. 1a).

*Hybrid weighting* combines both weighting schemes where each quartet is assigned with weight

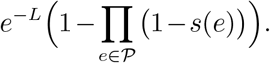

This weighting scheme aims to combine the strengths of both weighting by support and weighting by length and to improve over both; we will empirically show that such improvements are obtained.

While defining weighting schemes is easy, designing scalable algorithms to optimize the weighted quartet score is not. Adopting the existing ASTRAL algorithm to incorporate per-quartet weights is challenging for reasons elaborated in Material and Methods. A major contribution of this paper is designing a set of algorithms (Algorithm S1–S3) to optimize the weighted quartet using a set of new techniques paired with a dynamic programming (DP) step similar to ASTRAL. We leave the detailed description of the algorithm to the Optimization algorithm section; esp., see Theorems 3, 4, and 6 for correctness and Theorem 5 for the asymptotic running time being *O*(*kn*^1.5+*ϵ*^*H*) where *H* is the average gene tree height.

### Simulation results

#### Comparison of weighting schemes

We start by comparing the accuracy of weighting schemes and branch support types on two simulated datasets (S100 and S200). Our default method for computing branch support, used unless otherwise specified, is approximate Bayesian supports from IQ-TREE (aBayes) normalized to range from 0 to 1.

*S100*. This dataset adopted from Zhang et al. 2018 has gene trees inferred from sequences with varying lengths resulting in various levels of gene tree error (see Datasets). In most cases, weighting by support (wASTRAL-s) produces species trees with higher accuracy than weighting by length (wASTRAL-bl), and the improvements are statistically significant (Fig. S1); p-value < 10^-15^ according to a repeated-measure ANOVA test (see Statistical tests). The improvement in accuracy varies with *k* (*p* < 10^-15^) and perhaps sequence length (*p* ≈0.04). The accuracy of hybrid weighting (wASTRAL-h) on average is better than the accuracy of wASTRAL-s on all model conditions (*p* < 10^-10^) and the improvement in accuracy may depend on *k* (*p* ≈ 0.06) and sequence length (*p* ≈ 0.03). With ≥ 500 genes, wASTRAL-h is better than *both* support and length, showing that combining the two weightings makes wASTRAL-h more powerful.

On this dataset, bootstrap support computed using FastTree-2 is provided by Zhang et al. 2018. Thus, we also compute weighted ASTRAL trees using bootstrap supports (wASTRAL-s* and wASTRAL-h*). For weighting by support, aBayes weighting is much better than bootstrap weighting (*p* < 10^-15^), but the gap in error significantly (*p* < 10^-9^ for both) shrinks as *k* and sequence length increase (Fig. S1). For hybrid weighting, aBayes weighting is, on average, only slightly better than bootstrap weighting (the mean error increases across all conditions by only 0.2%).

*S200.* This 200-taxon dataset has species trees sampled under two birth rates (10^-6^,10^-7^), which control whether speciations are dispersed at random or closer to the tips (Fig. S2), and tree heights, which control levels of ILS (see Datasets). On this dataset, bootstrapped gene trees are not available; instead, local SH-like support from FastTree-2 is available, which we use (wASTRAL-s* and wASTRAL-h*). Patterns of accuracy across wASTRAL versions are similar to S100 (Fig. S3) as wASTRAL-h is more accurate than wASTRAL-s on all model conditions (*p* < 10^-6^), and the improvements depend on *k* (*p*≈ 10^-4^), ILS level (*p*< 10^-7^), and birth rate (*p* < 10^-10^). Using SH-like support with wASTRAL-h is, on average worse than aBayes support, increasing the error by 9%.

#### Comparison of topological accuracy to other methods

We next compare wASTRAL-h, the most accurate version of wASTRAL, to other methods.

*Impact of gene tree estimation error (S100 dataset).* On the S100 dataset (Fig. 2a and S4), wASTRAL-h is more robust to gene tree estimation error than ASTRAL-III, regardless of whether low bootstrap support (BS) branches (≤ 5%) are contracted. While contracting low support branches improves the accuracy of ASTRAL-III, weighting improves accuracy even more. For example, the average error with 1000 200bp genes goes down from 9% with ASTRAL-III to 7% after contracting ≤ 5% BS branches and 6% with wASTRAL-h. While wASTRAL-h dominates ASTRAL-III in all conditions with or without contraction (*p*< 10^-15^), the difference in accuracy varies across sequence lengths (*p*< 10^-6^ without contraction and *p*≈0.003 with contraction). Similar to wASTRAL-h, wASTRAL-h* has mean error lower than that of ASTRAL-III-5% in every condition (*p*< 10^-11^).

**FIG. 2.**
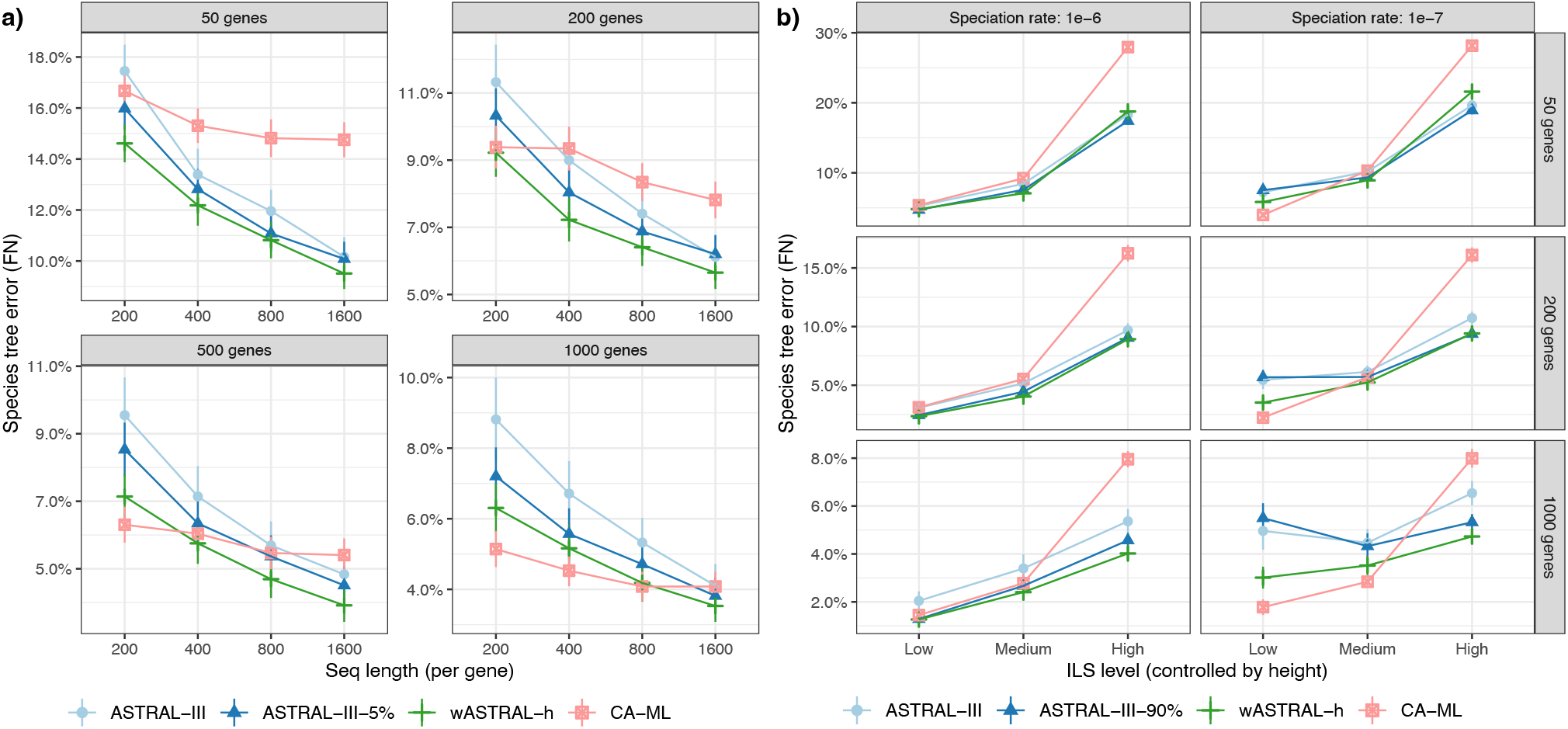
Species tree topological error on simulated datasets, comparing weighted ASTRAL hybrid (wASTRAL-h) against ASTRAL-III using fully resolved and contracted gene trees and concatenation using ML (CA-ML). (a) Results on the S100 dataset with *k* = {50,200,500,1000} gene trees (boxes) and gene sequence length {200,400,800,1600} (x-axis). Gene trees and CA-ML both inferred using FastTree-2. ASTRAL-III-5% contracts branches with < 5% BS. (b) Results on the S200 dataset with *k* = {50,200,1000}, rates of speciation 1E-6 and 1E-7, and three ILS levels. Gene trees and CA-ML both inferred using FastTfee-2. ASTRAL-III-90% contracts branches with aBayes support < 90%. See Fig. S4 and S5 for box plots.

The clearest patterns are observed when comparing wASTRAL-h and concatenation using ML performed using ML (CA-ML). While increasing the sequence length (and hence reducing the gene tree error) dramatically reduces the error of all ASTRAL variants, it has a much more subdued impact on CA-ML. As a result, the relative accuracy significantly depends on *k* (*p*< 10^-15^) and gene sequence length (*p*< 10^-9^) and the choice of the best method varies across conditions. Generally, wASTRAL-h tends to be more accurate than CA-ML under smaller *k* and greater sequence lengths. With *k* ≤ 200, wASTRAL-h dominates CA-ML for all sequence lengths. With *k*> 200, CA-ML is better for smaller gene alignments, and wASTRAL-h is better for longer alignments, with the only conditions when CA-ML has noticeable improvements over wASTRAL-h corresponding to 200bp genes.

*Impact of ILS level (S200 dataset).* On the S200 dataset that controls levels of ILS (see Datasets), overall, error rates of wASTRAL-h are lower than that of ASTRAL-III (Fig. 2b and S5) and the improvements are significant (*p*< 10^-15^). The improvements of wASTRAL-h compared to ASTRAL-III increase with more gene trees (*p* ≈ 7 × 10^-4^) but appear to decrease with more ILS (*p* ≈ 0.08). While Mirarab and Warnow 2015 reported no improvement in accuracy when contracting branches with low SH-like support, contracting branches with aBayes support < 90% (ASTRAL-III-90%) does improve accuracy. Nevertheless, wASTRAL-h has yet lower error (*p* < 10^-5^). Also, improvements of wASTRAL-h are significantly larger for the 10^-7^ birth rates, which tend to have earlier speciations (Fig. S2), than the 10^-6^ rate (*p*≈ 1.5 × 10^-5^).

The comparison between wASTRAL-h and CA-ML significantly depends on several factors (birth rate: *p*< 10^-7^; ILS: *p*< 10^-15^; *k: p*< 10^-11^). Overall, CA-ML is less robust to ILS levels and is always worse than wASTRAL-h when ILS is high and in most cases when ILS is at the medium level. However, with low ILS and birth rate =10^-6^ (more recent speciation), wASTRAL-h is better than CA-ML (*p*≈ 1.7 × 10^-5^) while with low ILS and birth rate =10^-7^ (earlier speciation), CA-ML is better (*p*< 10^-11^). Thus, in some conditions with low enough ILS, wASTRAL-h has reduced but not eliminated the gap between ASTRAL and CA-ML. For example, given 1000 gene trees and low ILS with 10^-6^ birth rate, ASTRAL-III has 5% error, which is not helped by branch contraction, whereas wASTRAL-h has 3%, which is much closer to the 2% achieved by CA-ML. To summarize, wASTRAL-h retains and magnifies the advantages of ASTRAL-III over CA-ML for high ILS conditions and eliminates or reduces the advantages of CA-ML under medium and low ILS conditions.

#### Support accuracy

We next test whether, by accounting for gene tree uncertainty, wASTRAL improves support values computed using the local Posterior Probability (PP) measure (see Branch support). We examine the calibration of support (i.e., whether the support matches the probability of correctness of a branch), its ability to distinguish correct and incorrect branches examined through Receiver operating characteristic (ROC) curves, and distributions of support (see Evaluation criteria).

*S100.* While wASTRAL-h generally gives higher support values than ASTRAL-III (Fig. S7), it has fewer cases of highly supported incorrect branches, especially with higher *k* and shorter sequences (Fig. 3a). For both ASTRAL-III and wASTRAL-h, while increased support often leads to increased frequency of correctness (Fig. 3b), support under-estimation or over-estimation can also be observed for certain sequence length and *k* combinations. For example, wASTRAL-h has a tendency to overestimate for large *k* values and short sequences. In terms of predictive power, for any desired false positive rate (FPR), the recall of wASTRAL-h is as good as or better than ASTRAL-III in all conditions (Fig. 3c), though the improvements in ROC can be small. Moreover, in most conditions, the minimum FPR obtained by wASTRAL-h (e.g., at 1.0 support) is lower than the minimum FPR obtained by ASTRAL-III.

**FIG. 3.**
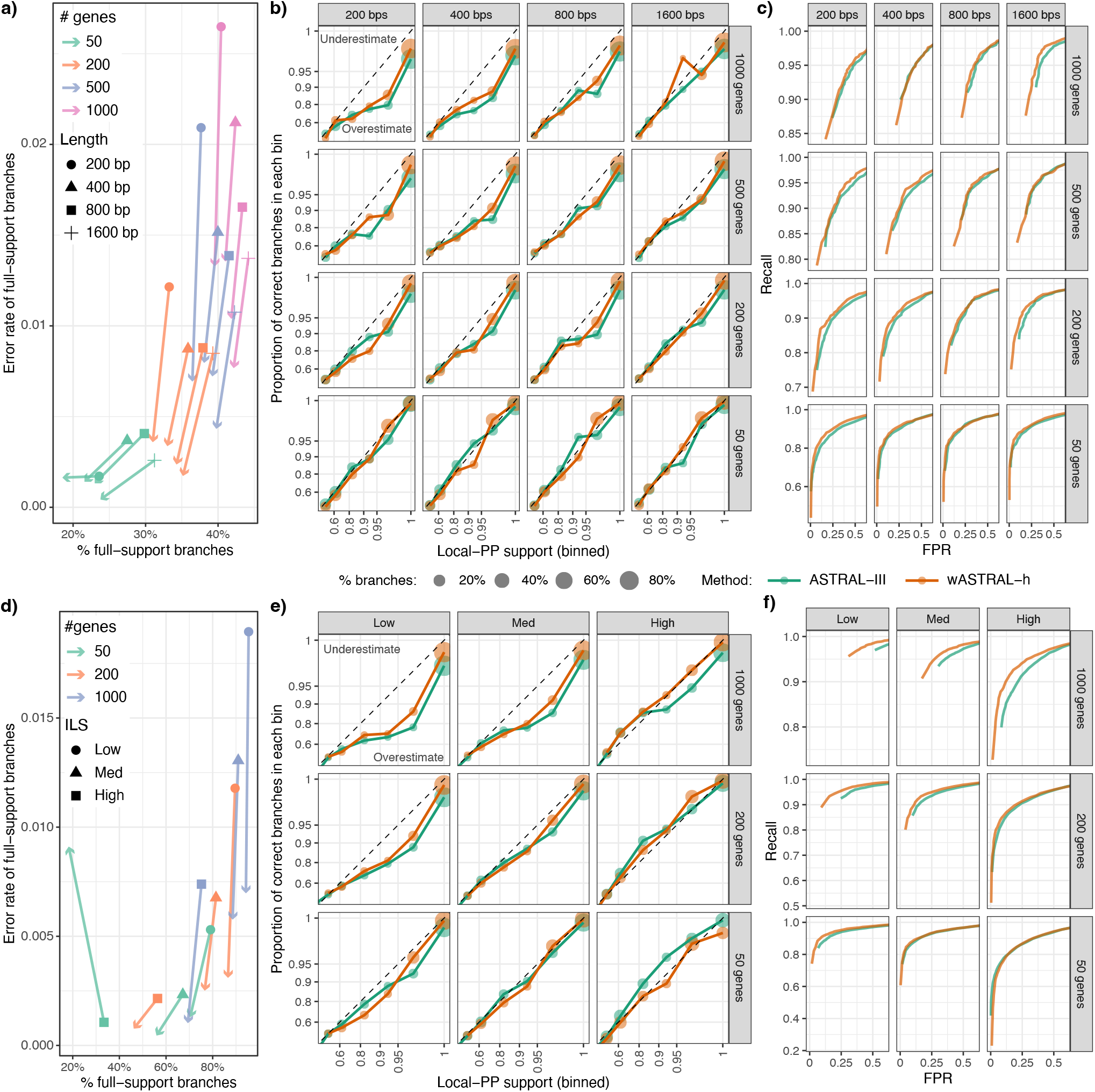
Support accuracy across (a-c) S100 dataset with *k* = {50,200,500’1000} and sequence length {200,400’800,1600} and (d-f) S200 dataset with *k* = {50,200,1000} and levels of ILS from low to high. (a,d) Change in 100% support branches. Each line shows the portion of full-support branches that are wrong (y-axis) and the percentage of all branches that have full support (x-axis) for wASTRAL-h (the arrowhead) and ASTRAL-III (other shapes). Arrows pointing downwards indicate less frequent errors in wASTRAL-h. (b,e) Support calibration. Branches are binned by their support, and for each bin, the percentage of branches that are correct are depicted versus the center of the bin. The dotted lines indicate ideal (calibrated) support. Top (bottom) triangle corresponds to the under-estimation (over-estimation) of support. (c,f) Receiver operating characteristic (ROC) curves where each dot corresponds to a contraction threshold, (Evaluation criteria). See Figs. S6-S11.

*S200.* Support values on the S200 dataset exhibit similar patterns to S100 (Fig. 3d-f). The most notable difference is that when *k* = 1000, wASTRAL-h has a clear advantage over ASTRAL-III in trading off precision and recall according to ROC curves (Fig. 3f and S9). This advantage shrinks as *k* decreases. Here, wASTRAL-h has a slight tendency to under-estimate support values < 1 (Fig. S10 and S11), and this tendency is most pronounced with 50 genes, high ILS level, and birth rate 10^-6^ (Fig. 3e and S11).

#### Comparison of the optimization algorithms

Assigning weights to quartets forced us to develop a new optimization algorithm, which can also be used for unweighted optimization. We next study whether the new optimization algorithm (denoted as DAC) is as effective as that of ASTRAL-III (denoted as A3) when no weights are used.

Testing on the S200 dataset, without missing data, DAC is in most cases slower than the A3 (Figs. 4a and 4c), a pattern that is pronounced with lower ILS levels. The change in relative running time with ILS levels is due to the dependence of the search space of A3 but not DAC on gene tree discordance levels (Zhang et al., 2018). In terms of accuracy, DAC and A3 are comparable for low and medium ILS levels (Fig. 4c). However, in the high ILS case, A3 is clearly better with only 50 genes, slightly better with 200 genes, and perhaps slightly worse with 1000 genes. Cases with reduced accuracy also have reduced quartet scores for the 50 genes scenario and high ILS (Fig. 4a), showing that A3 is preferable, especially with few gene trees. Thus, the improved accuracy of wASTRAL over ASTRAL-III is *despite* the fact that its DAC optimization algorithm is not always as effective as A3.

**FIG. 4.**
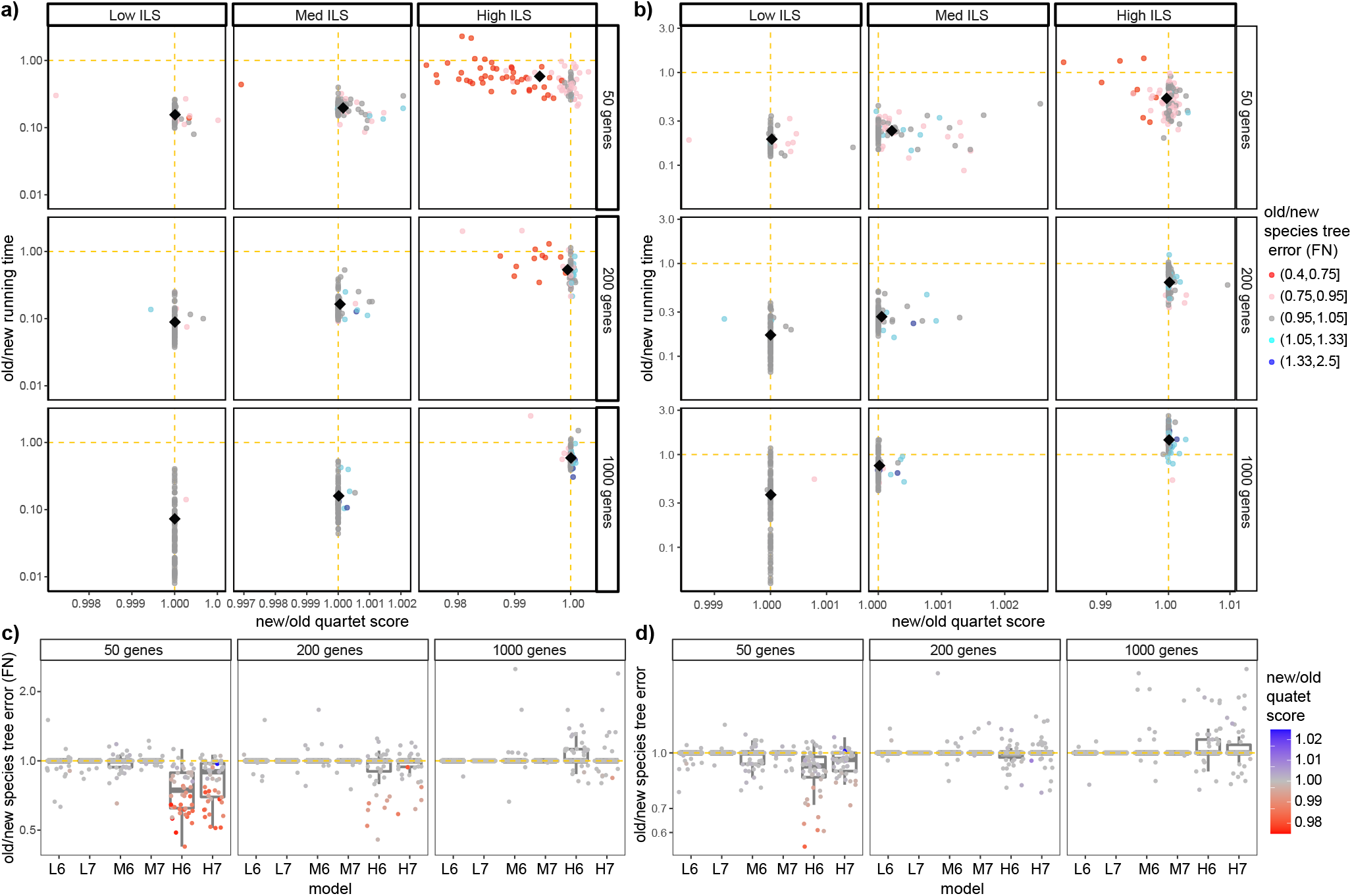
Comparison of the running time, quartet score, and accuracy between the old DP-based (A3) and the new optimization algorithms (DAC), both run without weighting on the S200 dataset. a,b) The ratio between the running time and quartet scores before (a) and after (b) randomly removing 5% of taxa from each gene tree; colors denote the ratio of species tree estimation error between the two methods. Note that the upper-right corner and blue color favor DAC. Results are separated by ILS levels from low to high and by *k* = {50,200,1000}. c,d) The species tree topological error using the A3 algorithm divided by the DAC algorithm before (c) and after (d) randomly removing of 5% taxa from each gene tree with colors denoting the ratio of quartet scores. L6 to H7 indicate model conditions with low, medium, and high ILS with 1E-6 and 1E-7 rates.

These patterns change when we add low levels of missing data by randomly removing 5% of leaves in each gene tree (Figs. 4b and 4d). DAC becomes closer to A3 in terms of running time in most cases and is even faster with high ILS and *k* = 1000 (Fig. 4b). Regarding accuracy, A3 and DAC are comparable in low and medium ILS levels (Fig. 4d). However, in the high ILS case, the error of A3 is slightly less, comparable, and slightly higher with 50, 200, and 1000 genes, respectively. Substantial changes in accuracy are caused by changes in quartet scores (Fig. 4b). Thus, DAC is competitive or better than the A3 in the presence of even low levels of missing data found to varying degrees in biological datasets.

### Biological data

We next study seven biological datasets (Datasets). On the canis dataset, which was the only input with at least 5 hours of running time for wASTRAL-h (Table S2), we also examine the running time.

#### OneKp

Overall, 47 out of 1175 (4%) branches change between the published ASTRAL-III tree and our wASTRAL-h tree. Most of these branches had low support in the ASTRAL-III tree (mean: 62%, max: 99%) but not in the wASTRAL-h tree (Fig. S12). OneKP Initiative 2019 focused most of their attention on 20 branches, corresponding to nine major evolutionary events that have been historically hard to resolve (e.g., early Eudicot diversification). Among 47 branches that change in wASTRAL-h, four of them are among the 20 focal branches. Beyond topological changes, the support values tend to increase in wASTRAL-h (Fig. 5a). In particular, all of the 20 focal branches that had less than full support in the original ASTRAL-III tree have increased support in the wASTRAL-h tree, leaving only four with support below 0.95 (as opposed to 12 branches with ASTRAL-III).

**FIG. 5.**
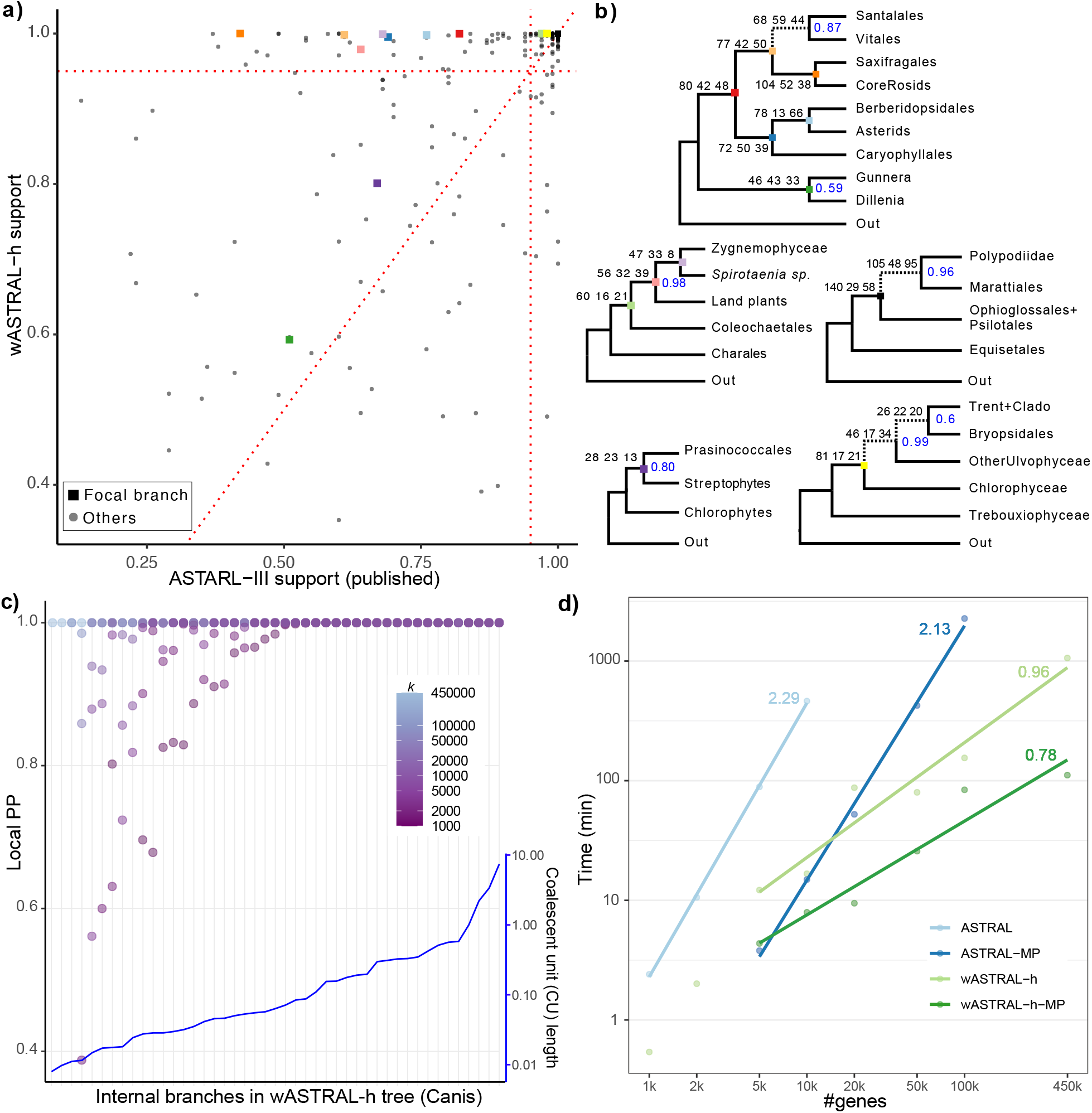
Results on OneKp (a,b) and canis (c,d) datasets. (a) Local posterior probabilities (PP) support of all species tree branches shared between wASTRAL-h and the published ASTRAL-III. Focal branches (squares) with support less than 100% in one of the two trees are colored and labeled in panel b. (b) wASTRAL-h resolutions of focal branches that differ from ASTRAL-III in topology or support. Branch labels: total weights of all quartets around each branch for the three possible topologies computed using (7) with weights coming from (5); the species tree topology is shown first. Node labels: localPP support when not equal to 100%. Dashed: focal branches that differ from ASTRAL-III. (c) Local PP of wASTRAL-h internal branches versus the number of genes *k* for each branch found in the wASTRAL-h output tree with all gene trees as input (x-axis). The inset with right y-axis scale shows the internal branch lengths in coalescent units on ASTRAL-III tree, sorted from low to high. The leftmost three branches are found only with *k* ≥ 100000. d) Log-log plot of total running time of ASTRAL-III and wASTRAL-h using both a single core (light colors) and 16 cores (dark colors) vs *k* on the canis dataset for *k* ranging from 1000 to 450000; Slopes of fitted lines, which estimate asymptotic growth exponent, are labeled. All test cases are performed on a server with AMD EPYC 7742 CPUs.

Significantly, all four focal branches that change from ASTRAL-III to wASTRAL-h become consistent with CA-ML, whereas the original ASTRAL-III tree was inconsistent with CA-ML. At the base of eudicots, Vitales (grapes) becomes sister to Santalales in wASTRAL-h tree with moderate support (0.87), which is consistent with CA-ML (Fig. 5b). Two branches in the so-called TUC clade also change: ASTRAL-III breaks down the class Ulvophyceae by uniting Bryopsidales with Chlorophyceae while wASTRAL-h recovers Ulvophyceae as sister to Chlorophyceae, which is the traditional resolution and is in agreement with CA-ML. Finally, the early diversification of ferns differs between CA-ML and ASTRAL-III but is identical between CA-ML and wASTRAL-h. Thus, wASTRAL-h makes coalescent analyses more congruent with CA-ML for the focal branches.

#### Canis

On the canis dataset of Gopalakrishnan et al. 2018 that spans a relatively shallow time scale (many branches are among populations of the same species), the majority of branches of the ASTRAL-III tree are shorter than 0.1 CU (Fig. 5c). Despite that, due to the large numbers of genes used, both wASTRAL-h and ASTRAL-III produce species trees with at least 99% support on all branches (Fig. S13). The ASTRAL-MP tree (on 100k gene trees) is identical to the published consensus tree, while the wASTRAL-h tree (on 450k gene tees) differs from it in only one branch (i.e., placement of the Egyptian dogs).

The linear running time scaling of wASTRAL-h with respect to *k* enables us to analyze randomly sampled subsets of 1000-450000 genes (Fig. 5c). The shortest branches need very many genes to achieve universal full support. Using fewer genes (even as many as 100,000) always leaves at least one branch with less than 99% support. Since many of the shortest branches are within species, a tree-like model of evolution is likely insufficient for such branches (Gopalakrishnan et al., 2018). Longer branches, which are mostly across species, do not require large numbers of genes to reach high support; the 21 longest branches have at least 99% support with as few as 1000 gene trees. Furthermore, wASTRAL-h is more scalable compared to ASTRAL-III with respect to the number of genes *k* (Fig. 5d). As Theorem 3 predicts, the running time of wASTRAL-h scales almost linearly with k, while ASTRAL-III scales close to quadratically (Fig. 5d and Fig. S14). ASTRAL-III fails to finish for *k* ≥ 2 × 10^3^ within 24 hours, and ASTRAL-MP with 16 cores takes more than 36 hours for *k* = 10^5^. By contrast, wASTRAL-h finishes on *k* = 4.5 × 10^5^ within 18 hours and 2 hours with one and 16 cores, respectively. Even when *k* = 10^3^, ASTRAL-III takes 4× more than wASTRAL-h due to the high levels of gene tree discordance and abundance of missing data, both of which increase the running time of ASTRAL-III but not wASTRAL-h.

#### Avian

On the avian dataset, the wASTRAL-h tree fully agrees with the ASTRAL-III trees after contracting low support branches and is very similar to original trees published by Jarvis et al. 2014 based on CA-ML (only five branches differ) and statistical binning (only two branches differ). This is in contrast to the ASTRAL-III tree without contraction from Zhang et al. 2018, which is in conflict with strong results from the literature and other methods. Moreover, all but one branch in the wASTRAL-h tree has higher or equal support compared to ASTRAL-III with any thresholds of contraction (Fig. S15). Interestingly, the only branch that experiences a reduction in support, the placement of Caprimulgimorphae as sister to Telluraves (core land-birds), is a branch that disagrees with both the published CA-ML and statistical binning trees. Finally, four branches with 99-100% support in wASTRAL-h are found by all coalescent-based methods (wASTRAL-h, ASTRAL-III and binned MP-EST) but not CA-ML, possibly pointing to a consistent signal that can be recovered only using coalescent-based analyses.

#### Cetaceans

The wASTRAL-h tree (Fig. S16) is similar to ASTRAL-multi and CA-ML trees reported by McGowen et al. 2020 with only a few differences (three branches to ASTRAL-multi and four to CA-ML). Interestingly, wASTRAL-h agrees with CA-ML and earlier studies (McGowen et al., 2009) and disagrees with ASTRAL-multi tree on the position of the Lissodelphis with high support (though the placement has low support in the ASTRAL-multi). On the other hand, both wASTRAL-h and ASTRAL-III break the monophyly of the genus Tursiops as *Tursiops truncatus* moves away from *Tursiops aduncus* and Stenella with high support. The question of the monophyly of Tursiops, supported by morphology, has been answered differently in two recent analyses and remains likely (Moura et al., 2020) but uncertain due to evidence for gene flow (Guo et al., 2022). Close to Tursiops is also the placement of the two *Stenella clymene* individuals, which is a known hybrid species evolved from *Stenella longirostris* and *Stenella coeruleoalba*. Interestingly, the two *Stenella clymene* individuals are placed apart, one as sister to *Stenella longirostris* and the other at the most recent common ancestor of *Stenella longirostris* and *Stenella coeruleoalba*. This placement is in contrast to CA-ML, which puts both individuals as sister to *Stenella longirostris*. Beyond Delphininae, two branches, the placements of *Orcinus orca* and *Neophocaena phocaenoides*, disagree with both ASTRAL-multi and CA-ML, but both branches have very low support in wASTRAL-h and cannot be trusted. These two are among 11 species where McGowen et al. 2009 used data from existing genomes and transcriptomes instead of their own targeted capture, and it is possible that differences in the analytical pipeline may have caused the low support in wASTRAL.

#### Insect datasets

On all three insect datasets, the differences between wASTRAL-h and ASTRAL-III are minimal and strictly limited to branches with low support. On the Nomiinae dataset, there is no conflict among highly supported branches. wASTRAL-h and ASTRAL-III differ in only one low support branch, and both trees differ from CA-ML in two low support branches (Fig. S17). On the Lepidoptera dataset, only seven out of 200 branches differ between wASTRAL-h and ASTRAL-III, and all of these branches have support below 75% (Fig. S18). Across the tree, wASTRAL-h has slightly more branches with support above 95% than ASTRAL-III (173 versus 169). On the Papilionidae datasets, wASTRAL-h tree and ASTRAL-III tree share the same topology, and all branches in both trees have high (≥ 99%) support (Fig. S19).

## Discussion

We introduced a family of new weighting schemes for quartet-based species tree estimation, including weighting quartets by terminal branch length (wASTRAL-bl), internal branch support (wASTRAL-s), or both (wASTRAL-h). We saw that the combined method (wASTRAL-h) has the best accuracy among the three and dominates unweighted ASTRAL in terms of accuracy. We next further comment on more subtle patterns observed in the data and end by pointing out directions for future research.

### Further observations based on the results

The choice between CA-ML and summary methods has been a long-standing debate (Edwards et al., 2016; Giarla and Esselstyn, 2015; Leaché et al., 2015; Meiklejohn et al., 2016; Simmons and Gatesy, 2015). While CA-ML is inconsistent under MSC (Roch and Steel, 2015), the most careful simulation studies have found that the best method depends on the dataset: CA-ML has been more accurate when gene discordance is low *and* gene signal is limited, and summary methods have been more accurate when discordance is high. Other factors such as deep versus shallow radiations, changes in evolutionary rates across genes, heterotachy, and the number of genes may also matter. Since we cannot reliably predict the superior method in practice, studies often report both types of analyses. We saw that weighting dramatically reduced (but did not fully eliminate) the gap between CA-ML and ASTRAL in conditions with lower ILS or heightened gene tree error (Fig. 2). Overall, our results point to wASTRAL-h being a reasonable, if not always optimal, choice *regardless* of the condition. Consistent with simulations, on real datasets, we observed that wASTRAL-h eliminates many of the differences between ASTRAL and CA-ML. Thus, using wASTRAL-h can help reduce the long-standing challenge of getting incongruent results from different analyses.

In our simulations, wASTRAL-h dominates ASTRAL in all model conditions in terms of accuracy, leaving no incentive to prefer ASTRAL in this regard. Contracting low support branches improved ASTRAL trees, but the weighting is more accurate than contracting and does not require hard-to-tune (Bossert et al., 2021) thresholds. Interestingly, the improvements, which were modest in many conditions but substantial in others, appeared more pronounced as the number of genes increased. We speculate the reason is that with more genes, not only the noise in the frequency of observed quartet *topologies* reduces, but also, the quartet weights become less noisy. Thus, having more genes benefits wASTRAL in two ways (less topological noise and better weights), only one of which is enjoyed by ASTRAL.

While topological improvements of wASTRAL-h over ASTRAL were marginal in many cases, the improvements in support were dramatic. The percentage of full support branches that were wrong was reduced in wASTRAL by half or more in most conditions (Fig. 3ad), rendering the full support branches more reliable. This increase in precision did not come at the cost of lowering support. Both real and simulated datasets (e.g., Figs. S7 and S10) saw *increased* support with wASTRAL. Two aspects of how we compute support have changed (Branch support). One is the handling of missing data (see (8)); it can be easily shown that, all else being equal, this change will decrease the localPP. Thus, the increase has to be due to the second change, which is the incorporation of weights. Since localPP support is a function of discordance, the increased support is empirical evidence that down-weighted gene tree quartets tend to be those that are more incongruent with others and the output species tree.

Branch support used as input by weighted ASTRAL can be computed in numerous ways with vastly different computational requirements. One practical question is whether one method should be preferred and, if so, which? We tested three ways of computing support on simulated data and noticed that IQ-TREE’s aBayes has the best accuracy, closely followed by bootstrapping (Fig. S1). In contrast, SH-like support was noticeably less effective. IQ-TREE’s aBayes is a local measure of support (i.e., computed for the nearest neighbor interchanges around a branch), and a local notion of support is consistent with how we interpret branch support (i.e., as independent, leading to a product). Moreover, computing local support is much faster than bootstrapping. Thus, while bootstrapping is a good option in terms of accuracy, IQ-TREE’s aBayes support can be used to build an accurate *and* efficient pipeline. Nevertheless, note that in the presence of rouge taxa that move widely across a gene tree, local measures of support may provide high support for most branches, whereas global support can result in low support for many branches, effectively down-weighting that gene. In such situations, global support may be more robust.

### Limits and future work

The wASTRAL-s optimization, when solved exactly, gives a statistically constant species tree estimator given *estimated* gene trees under our MSC+Error+Support model. While this model is general, our assumptions about support values are strong, and support estimation methods do not necessarily fulfill them (e.g., see debates in Felsenstein and Kishino, 1993; Hillis and Bull, 1993; Susko, 2009). Thus, the proofs of consistency should be taken more as a theoretical justification of the weighting approach used rather than a prediction of behavior on real data. Support values that over or under-estimate branch supports (compared to our assumptions) may or may not lead to inconsistency of the method, as our assumptions are sufficient but not necessary. Future work can seek more forgiving conditions for support that retain consistency, or conversely, conditions where the method is misleading.

We only proved the statistical consistency of wASTRAL-s and wASTRAL-bl under the MSCand MSC+Error+Support models, respectively, and hope that future works can prove wASTRAL-h is also consistent. Even more intriguing is whether wASTRAL (which can take multi-individual/multi-copy trees as input) is statistically consistent under combined models of GDL and ILS, as ASTRAL-multi is (Hill et al., 2020; Markin and Eulenstein, 2021). This question is particularly important for datasets where assumptions of MSC are violated. For example, on the OneKP dataset, examining the relative support for the three topologies around each branch (Fig. 5b) reveals that the quartet frequencies do not always follow the MSC expectations (one high frequency and two equal low frequencies). We believe weighting will continue to be beneficial for models of GDL. However, it is unclear whether weighting by branch length is profitable when gene tree discordance is due to GDL and especially horizontal transfer; thus, we caution the use of branch length when these processes are suspected. Finally, future work can incorporate weighting in the ASTRAL-Pro (Zhang et al., 2020) algorithm that natively supports paralogy.

While wASTRAL-h was more accurate than ASTRAL-III, if we turned off the weights, the new optimization algorithm (DAC) was slower (in many conditions) and less accurate (in some conditions) than the old algorithm (A3). While DAC tended to be as accurate or more accurate in the presence of missing data (Fig. 4b), our simulation results had no missing data, showing that the improved accuracy of wASTRAL-h was due to a better optimization objective, not a better optimization algorithm. Similar to A3, DAC is also a heuristic method addressing an NP-hard problem. Just as the speed and accuracy of ASTRAL changed substantially through tweaks to the heuristics from ASTRAL-I to ASTRAL-III, we anticipate that future work can further increase our accuracy, speed, or both. ASTRAL is also finely optimized for CPU, GPU, and vectorization (Yin et al., 2019). Currently, wASTRAL is only trivially parallelized for CPU, and future work can further optimize the code and implement GPU parallelization.

Our simulations, like any other, lacked some of the complexities of real biological data (Philippe et al., 2017; Springer and Gatesy, 2018). We did not include recombination, horizontal transfer, gene flow, hidden paralogy, alignment error, mistaken homology, violations of the model of sequence evolution, or missing data. It can be hoped that weighting helps alleviate the effects of some of these other sources of error as well. However, since many of these can lead to high support for the wrong trees, there is no guarantee that weighting would not leave these misleading signals intact or even amplified. Methods for simulating many of these effects are available and can be used in future studies to compare wASTRAL with both CA-ML and unweighted ASTRAL. A related promising avenue for future research is exploring other ways of weighting quartets. For example, future work can incorporate homology and alignment quality metrics into the weighting schemes. The weights could also reflect other factors, such as evidence of heterotachy impacting gene trees (Braun et al., 2019) and deviations from stationarity (Jeffroy et al., 2006). Even more ambitious approaches could be imagined where biases in support estimation could be predicted using machine learning (Suvorov et al., 2020). In designing and testing such weighting schemes, one must remember that not every weighting method will allow fast optimization using DP.

Finally, several features of ASTRAL are missing from wASTRAL, but future work can address this limitation. Currently, wASTRAL does not output branch lengths since the natural branch lengths that it could compute would be in a hard to interpret unit (e.g., *CU* + 2 × *SU*). Future work can examine ways to compute branch lengths in substitution or coalescent units. Other missing features left to future work are the test of polytomy (Sayyari and Mirarab, 2018), integration with visualization tools such as DiscoVista (Sayyari et al., 2018), and completion of gene trees with respect to each other. Nevertheless, the most valuable features of ASTRAL-III, including handling multi-individual datasets, handling polytomies, and outputting branch support, are all supported.

## Material and Methods

### Common notations and background

Let 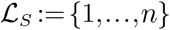 be a set of *n* species. Let us suppose that we are given a set of input binary gene trees 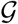 with 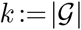. For each tree 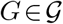, let its leaf set be 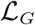 and its edge set be *E_G_*. For each branch *e*∈*E_G_*, we let *l_G_*(*e*) note its length. For a species set *A*, let *G* | *A* denotes *G* restricted to *A*. We refer to a set of four species as a quartet and define 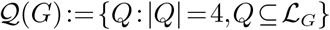 as the set of all quartets in *G*. We define *δ_G_*(*ab*|*cd*):=1 when 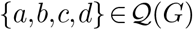 and *G*|{*a,b,c,d*} has topology *ab*|*cd*; otherwise we define *δ_G_*(*ab*|*cd*):=0. For nodes *u* and *v* of a gene tree *G*, we let 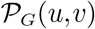 denote the set of branches on the path between *u* and *v* and let 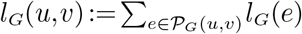. For a quartet *Q* = {*a,b,c,d*}, we denote 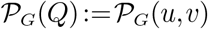, for *u* and *v* being nodes of *g* corresponding to the internal nodes (called the *anchors*) of *G*|*Q*; i.e., in case that *G*|*Q* has topology *ab*|*cd*, anchors are the only node on 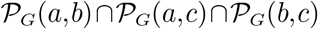 and on 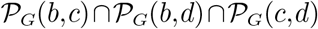.

We assume each true gene tree *G*\* is generated from the true species tree *S*\* under the MSC model. Branch lengths of *G*\* are in coalescent units (CUs). For each quartet 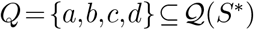 with topology *ab*|*cd* in the species tree, let *θ_Q_* = 1 — *e*^-*d*^ where *d* is the CU length of the internal branch of the quartet. Under MSC, for each true gene tree *G*\*, the following holds (Degnan, 2013): 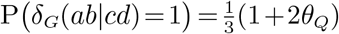 and 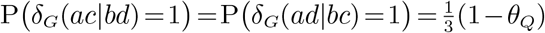. The input set 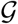 is a set of estimated gene trees, not true gene trees. In practice, these gene trees are estimated from sequence data using methods such as ML with branch lengths *l_G_*(*e*) given in the substitution-per-site units (SU). Moreover, input gene trees are furnished with support values: *s_G_*(*e*) maps each edge *e* of *G* to a support value in [0,1].

Theoretical results: improved consistency and sample complexity

For a given species tree topology *S*, we define its score against gene tree set 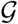 as

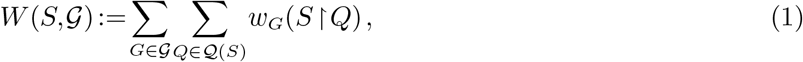

where *w_G_* is a function mapping a quartet of *G* to a number. In unweighted ASTRAL, for any {*a,b,c,d*},

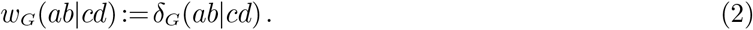

In this paper, we introduce three new ways of defining *w_G_*. Weighting by support sets:

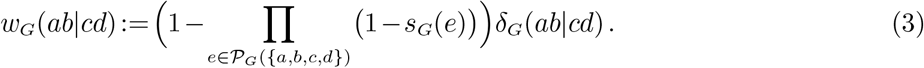

Weighting by branch length uses

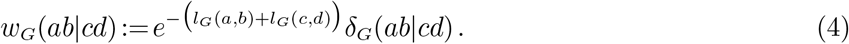

Finally, the hybrid weighting scheme combines weighting by support and weighting by length and uses:

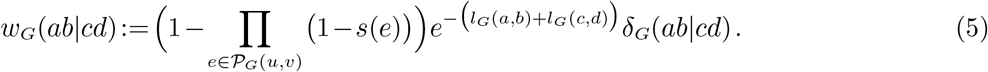

We study hybrid weighting only empirically but provide theoretical justifications for weighting by support (for estimated gene tree topologies) and weighting by length (for true gene tree topologies).

#### Weighting by support

Genes have varying levels of signal, and hence gene tree estimation error,and estimated gene trees can also be biased towards a specific topology due to factors such as long branch attraction. When bias goes against the species tree topology, unweighted ASTRAL can be positively misleading (Roch et al., 2019). It is reasonable to assume that gene trees with lower signals have lower support regardless of bias. By down-weighting those genes, wASTRAL-s can rescue consistency. To formalize this intuition, we introduce a model of gene tree error that allows us to make a more formal statement, showing that support weighted ASTRAL is consistent under some conditions where unweighted ASTRAL is not. **MSC+Error+Support model.** We assume each input estimated gene tree *G* is a draw from a distribution that depends on the true gene tree *G*\*. For each quartet 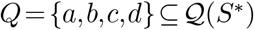 and each gene *G*, let *α_G,Q_* ∈ [0,1] denote a parameter controlling the quality of the estimated quartet gene tree *G*|*Q*. We assume *α_G,Q_* is independently drawn from the topology of *G*\* and we let the expected value and variance of *α_G,Q_* across genes be denoted by 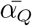 and 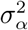. For each true gene tree topology, with probability *α_G,Q_*, we simply set the estimated gene tree to the true topology. With probability 1 – *α_G,Q_*, we choose among the three topologies with probabilities 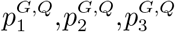. When these numbers are equal, there is no bias in gene tree estimation, and ASTRAL remains consistent (easy to prove). However, in our model, we allow systematic bias towards any topology. Let 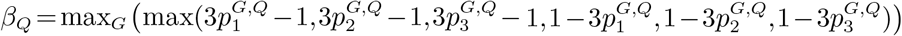 be the maximum bias towards or away any topology across genes. Under this model, the joint probability of true and estimated gene trees would follow the inequalities laid out in Table 2. For example, in the worst case, where 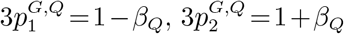, and 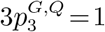, the joint distribution of true and estimated gene trees is given in Table 1 and depicted in Figure 1b.

**Table 2.** Joint probabilities (top) and weights (bottom) of estimated and true gene tree topologies under the MSC+Error+Support will follow the inequalities shown here. We omit *Q* and *G* superscript for brevity.

| $\mathbb{E}[(\cdot)(\cdot)\vert \alpha_{G,Q}]$ | $\delta_G(ab\vert cd)$ | $\delta_G(ac\vert bd)$ |
| --- | --- | --- |
| $\delta_{G^*}(ab\vert cd)$ | $\geq \frac{1}{3}(1+2\theta_Q)(\alpha_{G,Q} + \frac{1}{3}(1-\alpha_{G,Q})(1-\beta_Q))$ | $\leq \frac{1}{3}(1+2\theta_Q)(\frac{1}{3}(1-\alpha_{G,Q})(1+\beta_Q))$ |
| $\delta_{G^*}(ac\vert bd)$ | $\geq \frac{1}{3}(1-\theta_Q)(\frac{1}{3}(1-\alpha_{G,Q})(1-\beta_Q))$ | $\leq \frac{1}{3}(1-\theta_Q)(\alpha_{G,Q} + \frac{1}{3}(1-\alpha_{G,Q})(1+\beta_Q))$ |
| $\delta_{G^*}(ad\vert bc)$ | $\geq \frac{1}{3}(1-\theta_Q)(\frac{1}{3}(1-\alpha_{G,Q})(1-\beta_Q))$ | $\leq \frac{1}{3}(1-\theta_Q)(\frac{1}{3}(1-\alpha_{G,Q})(1+\beta_Q))$ |
| $\mathbb{E}[(\cdot)(\cdot)\vert \alpha_{G,Q}]$ | $w_G(ab\vert cd)$ | $w_G(ac\vert bd)$ |
| $\delta_{G^*}(ab\vert cd)$ | $\geq \frac{1}{3}(1+2\theta_Q)(\alpha_{G,Q} + \frac{1}{3}(1-\alpha_{G,Q})(1-\beta_Q))^2$ | $\leq \frac{1}{3}(1+2\theta_Q)(\frac{1}{3}(1-\alpha_{G,Q})(1+\beta_Q))^2$ |
| $\delta_{G^*}(ac\vert bd)$ | $\geq \frac{1}{3}(1-\theta_Q)(\frac{1}{3}(1-\alpha_{G,Q})(1-\beta_Q))^2$ | $\leq \frac{1}{3}(1-\theta_Q)(\alpha_{G,Q} + \frac{1}{3}(1-\alpha_{G,Q})(1+\beta_Q))^2$ |
| $\delta_{G^*}(ad\vert bc)$ | $\geq \frac{1}{3}(1-\theta_Q)(\frac{1}{3}(1-\alpha_{G,Q})(1-\beta_Q))^2$ | $\leq \frac{1}{3}(1-\theta_Q)(\frac{1}{3}(1-\alpha_{G,Q})(1+\beta_Q))^2$ |

We assume that for each quartet, the quartet support defined using (3) matches the probability of that topology being observed given the true gene tree. Thus, with our model for estimated gene tree distributions, the support of the quartet topology *i* is 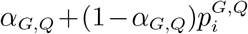 if it matches the true tree and 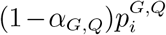 if it does not, leading to expected topology weights *w_G_*(.) given in Tables 1-2.

We now state our main results. Proofs of all results are given in Appendix Proofs.

##### Proposition 1.

*For each estimated gene tree G*, 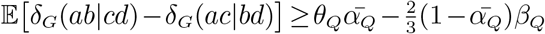 *and* 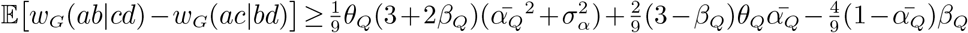.

For consistency of ASTRAL and wASTRAL-s, we need 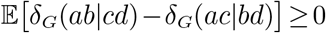 and 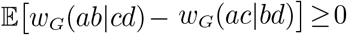, respectively. Figure 1e depicts the RHS of equations of Proposition 1, solving for *θ_Q_* and setting 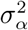 to zero (which is the worst-case for wASTRAL-s). It shows that wASTRAL-h is consistent for a larger set of species tree CU branch lengths, even in absence of any variation in gene tree quality. We next state this observation formally.

##### Theorem 1.

*Given estimated gene trees furnished with support generated under MSC+Error+Support model, there exist conditions where* (3) *guarantee a statistically consistent estimator of S*\* *but* (2) *does not, and the reverse is not true.*

#### Weighting by length

Our next result shows that using the length-based weighting function (4) leads to a larger gap than unweighted ASTRAL between the expected score of the true species tree and the alternative trees and thus has better sample complexity. Shekhar et al. 2017 has established that the number of gene trees required by ASTRAL to recover the species tree scales with *f*^-2^ as *f* → 0 where f is the CU length of the shortest species tree branch. Following that paper, we focus on the regime with *k* = Θ(*f*^-2^) gene trees and show a constant factor improvement in sample complexity. All theoretical results in this section assume that an input gene tree G matches the true gene tree *G*\* in *topology*.

The improved sample complexity essentially follows from the fact that under the MSC model, gene trees that match the species tree have shorter CU terminal branch lengths on average because discordance is caused by deep coalescence. However, a theoretical difficulty is that input gene trees have SU branch lengths instead of CU length. Thus, we need a model to translate CU lengths in *G*\* to SU lengths in *G*, capturing the effects of change in mutation rates and population sizes. We examine two such models.

##### Naive model

We start with a simple choice akin to a strict clock. Under this naive model, all branches of G are scaled from branches of *G*\* using a fixed multiplier λ.

##### Variable rate model

Let branches of the species tree *S*\* be broken into segments of arbitrary length (Fig. S20). For each gene tree *G*\*, a species tree in SU units *S*^†^ is drawn from a fixed distribution **D** (which does not change with *G*\*). *S*^†^ matches *S*\* in topology. The length of each segment *I* in *S*^†^ is scaled from the length of its corresponding segment in *S*\* using a multiplier 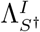. The set of all multipliers can be jointly drawn from any distribution as long as for each segment 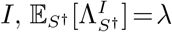. Segments in *S*\* can be used to divide *G*^*^ into segments defined at the same points along each branch (Fig. S20). The gene tree *G* is obtained from *G*\* by multiplying the CU length of each of its segments by the multiplier assigned to that segment in *S*^†^. Because segments have different multipliers (even though they have the same expectation), gene tree *G*^†^ deviates from ultrametricity. Because multipliers are drawn separately for each gene, deviations from ultrametricity happen in different ways across different genes.

We now state the results. Let *X_G_*: = *w_G_*(*ab*|*cd*) — *w_G_*(*ac*|*bd*) and *Y_G_*:= *δ_G_*(*ab*|*cd*) – *δ_G_*(*ac*|*bd*). Then,

###### Proposition 2.

*For a true quartet species tree S*\* *with topology ab*|*cd and input gene trees* 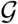 *generated under the naive model with any multiplier λ, let f be the distance between anchors of S*\*. *As f* → 0, *given k* = Θ(*f*^-2^) *gene trees, we have Var*[*X_G_*]=Θ_*f*_(1) *and*

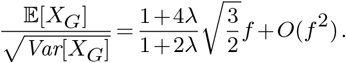

Similarly, under the variable rates model and assuming limited variance of rates across genes, we prove:

###### Proposition 3.

*For a true quartet species tree S*\* *with topology ab*|*cd and input gene trees* 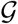 *generated under the variable rate model, let f be the distance between anchors of S*\* *and L be the total length of all other branches. Assume that for every branch segment I, the variance of its multiplier is bounded above*: 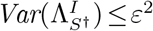 *where* 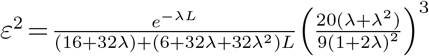 *As f* → 0, *given k* = Θ(*f*^-2^) *gene trees, we have Var*[*X_G_*]=Θ_*f*_(1) *and*

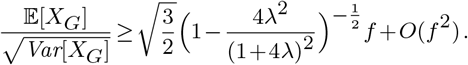

These propositions lead us to the main result.

###### Theorem 2.

*Under the conditions of Proposition 2 or Proposition 3*,

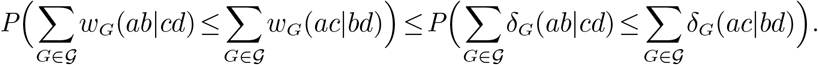

### Optimization algorithm

The objective of our optimization task is to find *S* maximizing 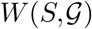 given in (1) for one of the *w_G_* functions (2)-(5). For a species tree *S*, let 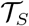 denote the set of tripartitions corresponding to the internal nodes of *S*. For a tripartition 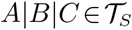 corresponding to an internal node *v* in *S* and a gene tree *G*, let *W*(*A*|*B*|*C,G*) be the total score of all shared quartets of *S* and *G* that anchor at *v*. Then,

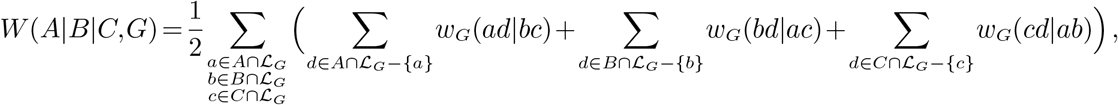

and 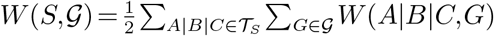.

ASTRAL-III uses a traversal of gene trees to compute *W*(*A*|*B*|*C,G*) with weight function (2) without enumerating all 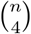 quartets. At each gene tree node, the total number of shared quartets between that node and *v* is computed using simple combinatorics. When quartets are weighted differently using weight functions (3)-(5), computing the aggregated weights of quartets around a node becomes more difficult as simple combinatorial equations become unavailable in the general case. Thus, we cannot simply use the same algorithm as ASTRAL and instead propose a new algorithm. In its simplest form (called the *base* version), the algorithm works as follows.

1. Starting from an empty tree, add each species to the tree one-by-one with a random order to obtain a full tree (see the Placement algorithm section and Algorithm S1). The algorithm also computes and stores tripartition scores *W*(*A*|*B*|*C,G*) for all tripartitions of the output tree.
2. Repeat the previous step for *r* rounds; by default *r* ∈ [16,32] (see details under Placement algorithm).
3. Combine results of the r rounds using a final dynamic programming (DP) step, which reuses the tripartition weights computed in step 1; each internal node of the output is constrained to be in at least one of the r greedy trees (see Dynamic programming section and Algorithm S2).

What makes this approach possible is step 1: a new algorithm that allows each addition to an existing tree to be performed optimally and efficiently. Importantly, while the base algorithm is a greedy heuristic, as Theorem 4 and the remark afterward will show, it retains the statistical consistency properties proved in Theorems 1–2. The running time of the base algorithm scales with *O*(*kn*^3^log(*n*)) in the worst case (Proposition 4) and is better with respect to *k* but worse with respect to n compared to ASTRAL-III, which is *O*((*kn*)^2.73^) in the worst case and roughly *O*(*k*^2^*n*^2^) in practice. Thus, we also propose a divide-and-conquer (DAC) algorithm for n ≥ 200 that uses the base algorithm on subsets of size 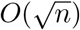 (see the DAC algorithm section and Algorithm S3). This strategy improves the running time to *O*(*n*^2.5+*ϵ*^*k*) under some assumptions, as detailed below under Theorem 5. The DAC algorithm also retains the statistical consistency guarantees (Theorem 6). We next detail each algorithmic component mentioned above.

#### Placement algorithm

Mai and Mirarab 2022 use the idea pioneered by Brodal et al. 2013 to design a quasi-linear algorithm to find the optimal placement of a species on a backbone tree that minimizes its quartet distance to a set of reference trees (e.g., gene trees). This algorithm traverses a binary (or multifurcating) species tree in a top-down manner and colors species using three (or more) colors, *A, B*, and *C*. When entering any node *u* of the species tree, all species under *u* are already colored A and all other species are colored *C*. At this point, the smaller child of *u* is recolored with *B*. The recoloring is done one species at a time; for each species, the path from the associated leaf in each gene tree to the root is visited, and several counters assigned to each gene tree node are updated. These counters enable calculating the score for placing the query on each species tree branch. After this recoloring is done and before moving from *u* to any of its children *v*, the sister of *v* is colored *C*, and if *v* is the smaller child of *u*, then *v* is changed back to *A*. This algorithm performs only *O*(*n*log*n*) species recoloring steps due to the smaller-child trick, which recolors the larger child of each node less often than the smaller child. Moreover, by representing each gene tree using an *O*(log*n*)-height tree called HDT adopted from Brodal et al. 2013, it ensures each recoloring takes *O*(*k*log*n*) time.

We build on the idea by Mai and Mirarab 2022 and adapt it to optimally solve the weighted quartet score placement problem (Algorithm S1), changing it in three substantial ways. *i*) We have created a new set of counters that enable us to compute the total *weighted* quartet score of all tripartitions resulting from all possible placements of the query. These counters essentially count the total weight of all the quartets with the same MRCA using recursive equations shown in Figure 6 and Table S1. The derivation of these counters is the heart of the algorithm but is too complex to detail here. We leave a full description to Proof of Theorem 3. *ii*) At each node *u*, we also recolor the query species as *A, B*, and *C* and recompute the counters; this allows us to compute the quartet score for all tripartitions resulting from all placements of the query. *iii*) Since our counters are more complex than Mai and Mirarab 2022, we use input gene trees instead of HDTs, which would be hard to implement. As a result, the cost of a leaf recoloring in our algorithm is *O*(*k*H) where *H* is the average height of gene trees instead of *O*(*k*log*n*) had we used HDTs. Note that for sufficiently balanced gene trees, *O*(*k*H) and *O*(*k*log*n*) are similar. We next prove that this algorithm finds the optimal solution.

**FIG. 6.**
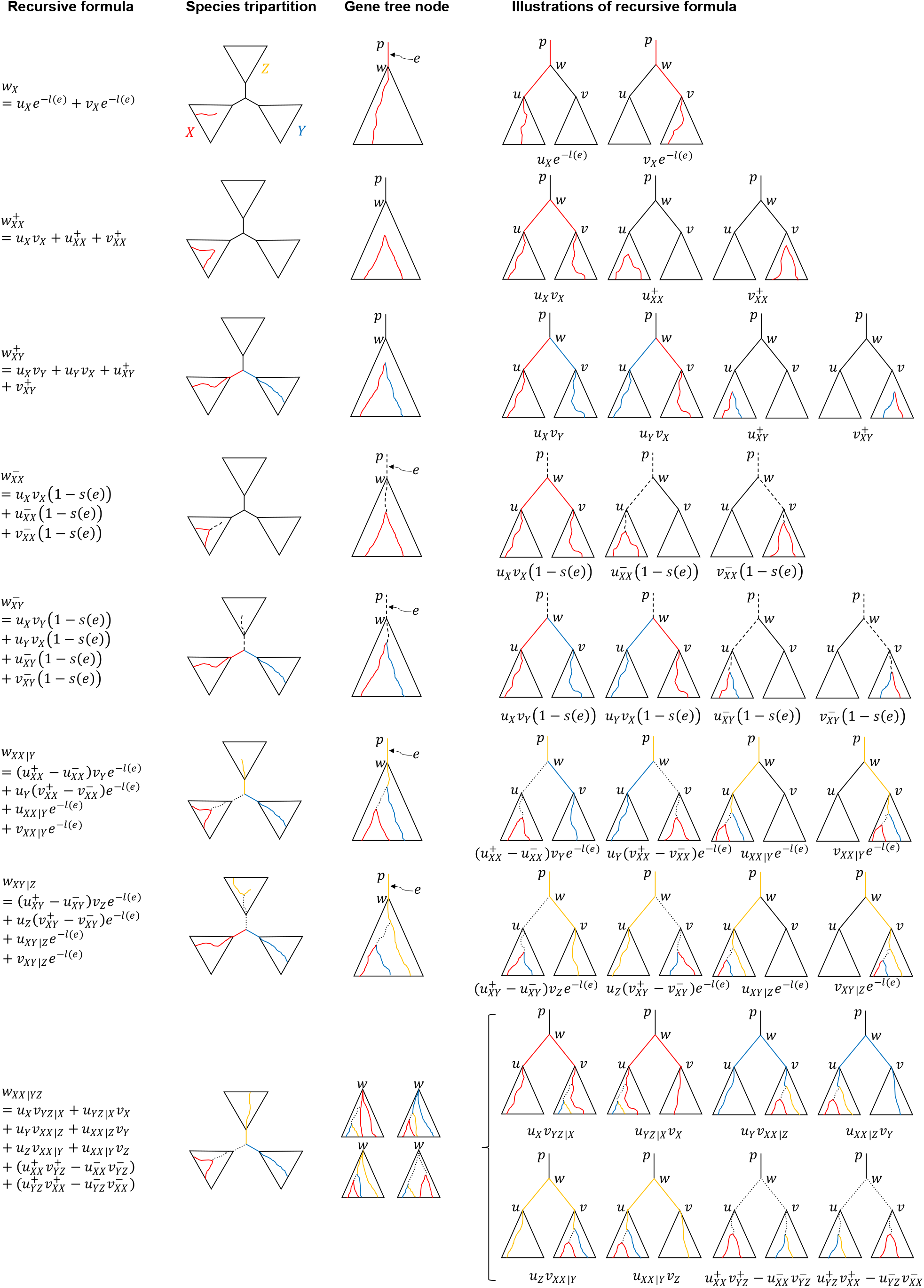
Recursive definitions of Counters. For a species tree tripartition 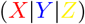, and a gene tree node *w,* we compute the total hybrid weight of all quartets anchored at the species tripartition and with w as the MRCA on the gene tree. Each solid colored path is weighed by the negative exponent of its length; each dashed path is weighted by one minus its support; each dotted path is weighted by its support. See also Table S1.

##### Theorem 3.

*Let S be a species tree, i be a species not in* 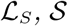 *be the set of possible species tree topologies by placing i onto S, and S*’ *be the output of Algorithm S1. Then, W* 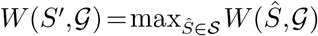.

While each individual placement is optimal, the greedy search is not guaranteed to find the optimal tree. We run the greedy search r times each of which produces a full tree *S_i_*. Empirically, we found *r* ≥ 4 to have minimal impact on the accuracy, but small improvements in the quartet score are observed for up to *r* = 32 rounds in outlier cases (Fig. S21). Based on these results, we set *r* (which the user can adjust) using a dynamic heuristic: *i*) start with 12 rounds and perform the DP algorithm to get an optimal score; *ii*) run another 4 rounds and perform DP using bipartitions from all previous rounds; iii) repeat step (*ii*) until no improvement to the optimal score is obtained or step (*ii*) has been repeated five times.

#### Dynamic programming

In each greedy search, we add the tripartitions of each *S_i_* and their weights to a lookup table *W*\*. The DP step computes an optimal species tree restricted to the tripartitions of *W*\* (Algorithm S2). The DP algorithm proceeds almost identically to ASTRAL, with one difference: While the search space in ASTRAL is the set of bipartitions found in all of the *S_i_* trees, here, the search space is the set of all tripartitions. With this change, we do not need to compute weight scores for any tripartition as those are precomputed and stored in *W*\* in the placement step.

##### Proposition 4.

*The time complexity of Algorithm S2 is O*(*k*H*n*^2^log*n*).

Since *H* = *O*(log*n*) for balanced trees and *H* = *O*(*n*) for caterpillar trees, the time complexity of Algorithm S2 is *O*(*kn*^2^ log^2^*n*) when gene trees are roughly balanced and *O*(*kn*^3^log*n*) when they are not. Note that because the counters are linearly related to counters of children of a node, in theory, the HDT structure can be adopted in our algorithm leading to a *O*(*kn*^2^ log^2^*n*) worst-case complexity. Since adopting HDT would add much more complexity for (potentially) little gain, we do not pursue it further.

Algorithm S2 is not guaranteed to find the optimal solution. However, a positive theoretical result ensures that this lack of optimality does not impede the statistical consistency of the solution:

##### Theorem 4.

*If there exists a species tree topology S* satisfying that for each quartet subtree ab*|*cd*,

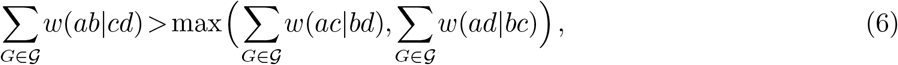

*then the output of Algorithm S2 will be S*\*.

##### Remark

*For a binary true species tree S*, as k*→∞, *S*\* *satisfies the condition of Theorem 4 with an arbitrarily high probability for wASTRAL-s under the assumptions of Theorem 1 and for wASTRAL-bl under the assumptions of Theorem 2. To see this point, note that due to the consistency of the estimator, for a quartet Q to achieve a high probability* 1 – *e*’ *a certain k_ϵ’,Q_ must be sufficient. Setting* 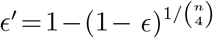 *and using a union bound, it is easy to see that k* = max_*Q*_ *k*_*ϵ*’,*Q*_ *is enoughto achieve the probability* 1 – *ϵ of correctness for all quartets. Thus, by Theorem 4, Algorithm S2 is a statistically consistent estimator of the species tree under the assumptions of Theorems 1 and 2. We conjecture that wASTRAL-h can also be proved statistically consistent under assumptions similar to Theorem 1 for topology and support and Theorem 2 for branch length*.

#### DAC algorithm

The DAC procedure (Algorithm S3) first computes a backbone tree on fewer species, adds all the remaining species onto the backbone tree, and then locally refines the topology around the backbone branch.

1. To compute a backbone tree *S_i_*, we randomly select 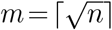 leaves and apply the Algorithm S2 with 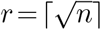 rounds of placements to get a backbone tree with *m* species.
2. For the remaining *n* – *m* species, we independently find their optimal placement on *S_i_* using the Algorithm S1. We group species placed on the same branch together to obtain 2*m* – 3 clusters.
3. For each cluster *C_e_* corresponding to a branch *e*, we sequentially place species in *C_e_* onto *S_i_* using the Algorithm S1 and remove any “orphan” species that are not placed on *e* or its derived branches; the result is called *S_e_*.
4. All trees in {*S_e_*: *e* ∈ *E_S_i__*} induce the same scaffold tree *S_i_* on their shared taxa. Thus, they can be easily merged into a uniquely defined tree 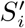.
5. If 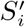 orphan species exist, at the end, we place them onto 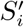 using the Algorithm S2.

The potential for orphan taxa makes it harder to establish the time complexity of the DAC algorithm theoretically, but a result can be proved:

##### Theorem 5.

*When the inequality condition in Theorem 4 is satisfied, then the time complexity of the DAC algorithm is O*(*n*^1.5+*ϵ*^*kH*) *with arbitrarily high probability.*

Similar to the base algorithm, the DAC algorithm retains statistical consistency.

##### Theorem 6.

*Under the conditions of Theorem 4, the DAC Algorithm S3 will output S*\*.

##### Remark.

*Under assumptions of Theorem 1 for wASTRAL-s and Theorem 2 for wASTRAL-bl, Algorithm S3 gives a statistically consistent estimator of the species trees.*

### Branch support

We adopt the quartet-based metric introduced by Sayyari and Mirarab 2016 used for measuring branch support. This metric essentially quantifies the probability of the true quartet score around a species tree branch being more than 3 given the observed quartet topologies assuming that gene trees are fully independent, but the quartets around the branch are fully dependent. The original metric gives all gene trees with at least one quartet around a branch of interest an equal weight of one. In wASTRAL-h, we instead weight each gene tree by the total support of all three topologies and normalize the counts. Removing an internal branch e of the species tree and its four adjacent branches defines a quadripartition of species *A|B|C|D*, and we assume (*A*∪*B*)|(*C*∪*D*) is the bipartition defined by e. Note that any quartet (*a,b,c,d*) ∈ *A* × *B* × *C* × *D* has the same internal branch as *e*. Let 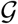 denote the subset of gene trees with at least one element from each of *A, B, C*, and *D*. We define *x*_1_, the normalized quartet count for branch *e*, as

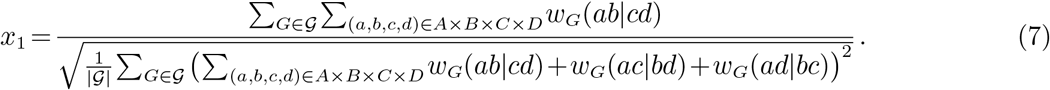

The quartet counts for (*A*∪*C*)|(*B*∪*D*) and (*A*∪*D*)|(*B*∪*C*) are similarly defined and are denoted by *x*_2_ and *x*_3_. This form of normalization models the observation that gene trees with higher weights also have higher variance in their weights. Using the localPP method of Sayyari and Mirarab 2016, we set the localPP support to: 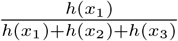, where 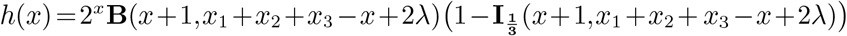, **B** is the beta function, **I**_x_ is the regularized incomplete beta function, and λ is birth rate in the Yule prior distribution (default: 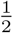).

When all weights are set to 1, as in ASTRAL-III, the new definition is identical to the original one in the absence of missing data but can be different with missing data. Let 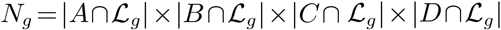 be the number of quartets around the branch of interest present in a gene tree *g*; let *n_g_* be the number of those quartets that are compatible with (*A*∪*B*)|(*C*∪*D*). Then, the old definition sets 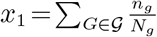 while the new definitions uses

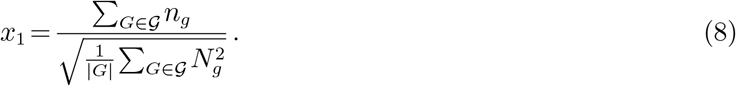

The two definitions are identical only when all *N_g_* values are the same, which is the case when there is no missing data but can also happen in other scenarios. When patterns of missing data are different, the old calculations made sure all genes had equal weights (each gene has *x*_1_ + *x*_2_ + *x*_3_ = 1). In the new definition, since each gene is weighted differently in wASTRAL, to begin with, we also allow genes to have a different total vote depending on their patterns of missing data. In the new formula, each gene votes (contributes to *x*_1_ + *x*_2_ + *x*_3_) proportionally to the number of quartets they have around a branch.

### Datasets

#### Simulated data

S100 Simulated dataset by Zhang et al. 2018, includes 100 ingroups and one outgroup and is simulated using SimPhy (Mallo et al., 2016) with 50 replicates. The species trees are simulated under the birth-only process with birth rate 10^-7^, a fixed haploid population size of 4 × 10^5^, and the number of generations sampled from a log-normal distribution with mean 2.5 × 10^6^. 1000 true gene trees are simulated under the MSC model. The ILS level substantially varies across replicates, with a mean of 0.46 when measured by the average normalized (Robinson and Foulds, 1981) (RF) distance between the true species trees and true gene trees. Gene alignments of length {200,400,800,1600} bps are simulated using Indelible (Fletcher and Yang, 2009) under the GTR model after assigning SU gene tree branch lengths that deviate from the clock using rate multipliers. Gene trees are reconstructed under the GTR+Γ model using FastTree-2 (Price et al., 2010). The gene tree estimation error, measured by the False Negative (FN) rate between the true gene trees and the estimated gene trees is {0.55,0.42,0.31,0.23} for lengths {200,400,800,1600}. The original publication has made bootstrap support obtained from 100 replicates run using FastTree-2 available for each gene tree.

S200 Simulated dataset by Mirarab and Warnow 2015 includes 200 ingroup species and an outgroup. Its species trees are generated under two different birth rates 10^-6^,10^-7^ each with 50 replicates and three different ILS levels, low (≈ 10%), medium (≈ 35%), and high (≈ 70%), controlled by max tree heights 10^7^,2 × 10^6^,5 × 10^5^ generations, respectively. The sequence length of each gene is uniformly drawn between 300 and 1500 bps, resulting in a wide range of gene tree estimation errors across replicates (mean: 25%, 31%, and 47%, for low, medium, and high ILS). Gene trees are estimated using FastTree-2, but because bootstrap replicates are not available, we compute aBayes support using IQ-Tree with fixed topologies.

By default, we compute branch length and support using IQ-TREE (v 1.6.12) aBayes option (`--abayes`) under GTR+Γ model. As each support value s is between 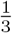 and 1, we normalize support value to 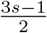 so that the minimum is 0. To run wASTRAL (which currently takes only binary trees as input), we randomly resolve polytomies in input trees with length and support set to 0, which is equivalent to a polytomy for wASTRAL-s and -h.

ASTRAL-III version 5.7.4 is used throughout. ASTRAL-III-5% (S100 dataset) denotes running ASTRAL-III on gene trees with low bootstrap support branches (< 5%) contracted. The 5% threshold is used because Zhang et al. 2018 found it to have the best accuracy overall. On the S200 dataset, because bootstrap support is not available, we instead rely on IQ-TREE aBayes support, which tends to be much higher than bootstrap support. Thus, we contract branches with support below a 0.90 threshold with aBayes, denoted as ASTRAL-III-90%.

CA-ML is performed using unpartitioned ML. On the S200 dataset, CA-ML was available from the original study (where they used FastTree-2 as the ML method) and is used here. On S100, we ran CA-ML using FastTree-2. Thus, on both datasets, the same tool is used for gene tree estimation and CA-ML, ensuring the comparisons are fair.

#### Biological datasets

Seven biological datasets were used.

*OneKP (OneKP Initiative, 2019)* dataset includes 1178 species spanning the plant tree of life obtained using transcriptomics. The original study has inferred 410 AA-based gene trees from putative single-copy genes using RAxML with bootstrap support (which we use), an ASTRAL-III species tree, and CA-ML using RAxML.

*Canis (Gopalakrishnan et al., 2018)* dataset includes 48 genomes across genus Canis with taxon sampling that allows reconstruction at both species and population levels. Loci with roughly 10kbp lengths were selected across the genome at random, leading to 449,450 gene trees. Since ASTRAL-II could not handle this size, the original study partitioned the gene tree into 100 subsets and inferred one ASTRAL-II species tree per subset and published a consensus of those trees. We used wASTRAL-h to analyze all the available gene trees, which the original paper estimated using FastTree-2; we were also able to analyze up to 100,000 gene trees using ASTRAL-MP (Yin et al., 2019) (within 48 hours). Due to the large number of genes, we simply use the provided SH-like FastTree-2 support instead of re-estimating support.

*Avian (Jarvis et al., 2014)* dataset includes 48 species designed to resolve the order-level avian relationships, which experience extremely high levels of gene tree discordance potentially due to a rapid radiation. Authors studied three data types: concatenation of exons per gene (exons), concatenation of introns per gene (introns), and Ultra Conserved Elements (UCE). Here, we analyze all 14,446 input gene trees (8251 exons, 2516 introns, and 3679 UCEs) with bootstrap-annotated branches available from the original study. The main challenge on this dataset is the low gene tree resolution, which led to the development of the statistical binning method (Mirarab et al., 2014a). Without binning, the analyses of all 14,446 loci resulted in species trees that were clearly wrong. More recently, species tree inferred from ASTRAL-III without dealing with gene tree error also resulted in incorrect species trees (Zhang et al., 2018); however, contracting low support branches (e.g., **≤** 3, 5, and 10%) appeared to solve the problem.

*Cetaceans (McGowen et al., 2020)* dataset includes targeted-captured exonic data for 100 individuals from 77 cetacean species and 12 outgroups. The original study estimated gene trees using RAxML under the GTRCAT model but without support for 3191 protein-coding genes. We computed Bayesian local supports and branch lengths for fixed gene tree topologies using IQ-Tree, and reanalyzed the dataset using wASTRAL-h. We compare the results to two trees produced by the original study: a CA-ML tree and an ASTRAL-multi tree that forces individuals of the same species to be grouped together.

*Insect datasets.* We also tested three insect datasets, in each case, using available gene trees. i) a 32-taxon collection of 853 RAxML gene trees with bootstrap supports obtained from alignments of ultraconserved elements focused on the bee subfamily Nomiinae and particularly genus Pseudapis (Bossert et al., 2021), ii) a 203-taxon set of 1930 RAxML gene trees with bootstrap support obtained from transcriptomic alignments focused on Lepidoptera (butterflies and moths) (Kawahara et al., 2019), and iii) a 61-taxon dataset of the Papilionidae (swallowtail butterflies) with 6407 IQ-TREE gene trees with supports that we computed using aBayes (Allio et al., 2020) and obtained from amino-acid alignments of orthologous protein-coding genes.

### Evaluation criteria

To compare topological accuracy, we use the false-negative rate (FN) in recovering bipartitions of the true species tree. Since the true species tree and the reconstructed species tree are both binary, falsenegative rate, false-positive rate, and normalized RF are all the same. For measuring the accuracy of support, we use three methods with different goals.

Calibration plots ask if support values perfectly indicate correctness (i.e., are *calibrated*). We break support values into these bins: [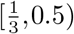, [0.5,0.75), [0.75,0.9), [0.9,0.95), [0.95,1), and {1} (note that 1 means anything rounded to 1 by the tool). For each bin, we compute the average accuracy of branches with support in that range and plot it versus the midpoint of the boundaries of that bin. On such plots, points above (below) diagonal indicate under-estimation (over-estimation) of branch support. Even when support is not calibrated, it can be useful if higher support *correlates* with correctness; e.g., if all support values are uniformly increased or decreased (say, divided by two), it can still be perfectly correlated with support. To measure this aspect, we use ROC curves. For a large number of thresholds between 0 and 1, we contract all branches with support below that threshold. We call contracted correct branches FN, contracted incorrect branches TN, kept correct branches TP, kept incorrect branches FP; these allow us to define true positive rate and recall, and thus ROC. Note that the ROC curve remains the same with any monotonic transformation of support values assuming an infinite number of thresholds. We also plot the empirical cumulative density function (ECDF) of correct and incorrect branches. We expect higher support for correct branches than for incorrect branches; thus, the accuracy can be judged by the gap between ECDF curves of correct and incorrect branches.

#### Statistical tests

We perform repeated measures ANOVA tests between two species tree reconstruction methods to test the significance of topological accuracy differences and whether the gap in accuracy depends on simulation model parameters. We limit the data to only the two methods being compared, and for each experimental condition, we use replicates as the repeated measures (i.e., the error term). We perform double-sided ANOVA tests on reconstruction methods vs. experimental conditions and report p-values for the difference between methods and the impact of other variables on that difference.

## Availability

The wASTRAL software is available publicly at https://github.com/chaoszhang/ASTER. Data used here are available at https://github.com/chaoszhang/Weighted-ASTRAL_data.

