## Supplement for "Weighting by Gene Tree Uncertainty Improves Accuracy of Quartet-based Species Trees"

### Supplementary material

**Commands**

#### Approximate Bayesian Branch Support Annotation

```
iqtree2 -s SEQ_ALIGNMENT -te GENE_TREE -m TVM+I+G4 -abayes -pre ANNOTATED_GENE_TREE
```

Note: When inferring support as a post-processing step, the same model used for inferring the tree should be used, a task that requires care when the original trees are inferred using a different tool (e.g., RAxML). TVM+I+G4 is simply an example.

#### Running wASTRAL

Exact commands when running on gene trees with approximate Bayesian/Bootstrap/SH-like supports.

```
astral-hybrid -x 1 -n 0.333 APPROXIMATE_BAYESIAN_ANNOTATED_GENE_TREE
astral-hybrid -x 100 -n 0 BOOTSTRAP_ANNOTATED_GENE_TREE
astral-hybrid -x 1 -n 0 SH_LIKE_ANNOTATED_GENE_TREE
```

**Table S1.** Counters $w_*^*$ are defined for each node $w$ in each gene tree, and $Q$ is defined globally. Here, $X$,$Y$,$Z$ are distinct colors of $A$, $B$, and $C$. Let $u$,$v$ be the children of $w$; $e$ be the parental edge of $w$; $p$ be the parent of $w$; $\mathcal{P}_{x,w}$ be the path between $x$ and $w$; $s(\mathcal{P})=1-\prod_{\hat{e}\in\mathcal{P}}(1-s(\hat{e}))$; $m(i,j)=$ MRCA of $i$ and $j$. Counters for leaves are set to zero unless explicitly noted. For each counter, we show a recursive equation on top and the equivalent non-recursive definition on the bottom.

| | |
|---|---|
| $w_X$ | $(u_X+v_X)e^{-l(e)}$ for internal node $w$; $e^{-l(e)}$ for leaf node $w$ colored $X$ |
| | $\sum_i e^{-l(\mathcal{P}_{i,p})}$ for all leaf nodes $i$ colored $X$ under $w$ |
| $(w^+_{XX},w^+_{XY})$ | $\left(u^+_{XX}+v^+_{XX}+u_Xv_X, u^+_{XY}+v^+_{XY}+u_Xv_Y+u_Yv_X\right)$ |
| | $\sum_{i,j} e^{-l(\mathcal{P}_{i,j})}$ for all leaf nodes $i$ colored X and $j$ colored X/Y under $w$ |
| $(w^-_{XX},w^-_{XY})$ | $\left((u^-_{XX}+v^-_{XX}+u_Xv_X)(1-s(e)),(u^-_{XY}+v^-_{XY}+u_Xv_Y+u_Yv_X)(1-s(e))\right)$ |
| | $\sum_{i,j} e^{-l(\mathcal{P}_{i,j})}\left(1-s(\mathcal{P}_{m(i,j),p})\right)$ for all leaf nodes $i$ colored X and $j$ colored X/Y under $w$ |
| $(w_{XX\|Y}, w_{XY\|Z})$ | $\Big((u_{XX\|Y}+v_{XX\|Y}+(u^+_{XX}-u^-_{XX})v_Y+u_Y(v^+_{XX}-v^-_{XX}))e^{-l(e)},$ <br> $(u_{XY\|Z}+v_{XY\|Z}+(u^+_{XY}-u^-_{XY})v_Z+u_Z(v^+_{XY}-v^-_{XY}))e^{-l(e)}\Big)$ |
| | $\sum_{i,j,k} e^{-l(\mathcal{P}_{i,j})-l(\mathcal{P}_{k,p})}s(\mathcal{P}_{m(i,j),m(i,k)})$ for leaf nodes $i$ colored X, $j$ colored X/Y, $k$ colored Z under $w$, and $m(i,j)$ under $m(i,k)$ |
| $w_{XX\|YZ}$ | $v_Xu_{YZ\|X}+u_Xv_{YZ\|X}+u_{XX\|Z}v_Y+v_{XX\|Z}u_Y+u_{XX\|Y}v_Z+v_{XX\|Y}u_Z$ <br> $+(u^+_{YZ}v^+_{XX}-u^-_{YZ}v^-_{XX})+(u^+_{XX}v^+_{YZ}-u^-_{XX}v^-_{YZ})$ |
| | $\sum_{h,i,j,k} w_G(hi\|jk)$ for all leaf nodes $h$,$i$ colored X, $j$ colored Y, $k$ colored Z, and $w=$ MRCA $h,i,j,k$ |
| $Q$ | $\sum_{G\in\mathcal{G}}\sum_w(w_{AA\|BC}+w_{BB\|AC}+w_{CC\|AB})$ for internal nodes $w$ in $G$ |
| | $\sum_{G\in\mathcal{G}}\sum_{h,i,j,k} w_G(hi\|jk)$ for leaf nodes $h,i,j,k$ in $G$ where $h$,$i$ have the same color and $i,j,k$ have different colors; when species coloring matches all gene trees, $Q=W[A\|B\|C]=\sum_{G\in\mathcal{G}}W(A\|B\|C,G)$ (Proposition 5). |

**Table S2.** Running time of species tree inference methods on biological datasets. We use 5.17.3 version of ASTRAL-III if not otherwise clarified.

| Dataset | $n$ | $k$ | Method | #Cores | Wall-clock time | CPU time |
|---|---|---|---|---|---|---|
| OneKP | 1178 | 410 | wASTRAL-h | 16 | 17.1 min | 4.57 hr |
| | | | ASTRAL-III (5.0.3) | 1 | 17.2 hr | 17.2 hr |
| Canis | 48 | 449450 | wASTRAL-h | 1 | 17.7 hr | 17.7 hr |
| Avian | 48 | 14446 | wASTRAL-h | 16 | 1.76 min | 28.1 min |
| | | | ASTRAL-III | 16 | 20.9 min | 5.57 hr |
| Cetacean | 98 | 3191 | wASTRAL-h | 16 | 35.2 sec | 9.39 min |
| | | | ASTRAL-III | 16 | 1.97 min | 31.5 min |
| Nomiinae | 32 | 853 | wASTRAL-h | 1 | 5.93 sec | 5.93 sec |
| | | | ASTRAL-III | 1 | 8.64 sec | 8.64 sec |
| Lepidoptera | 203 | 1930 | wASTRAL-h | 16 | 2.02 min | 32.3 min |
| | | | ASTRAL-III | 16 | 9.14 min | 2.44 hr |
| Papilionidae | 61 | 6405 | wASTRAL-h | 16 | 24.8 sec | 6.61 min |
| | | | ASTRAL-III | 16 | 1.11 min | 17.8 min |

#### Supplementary Figures and Tables

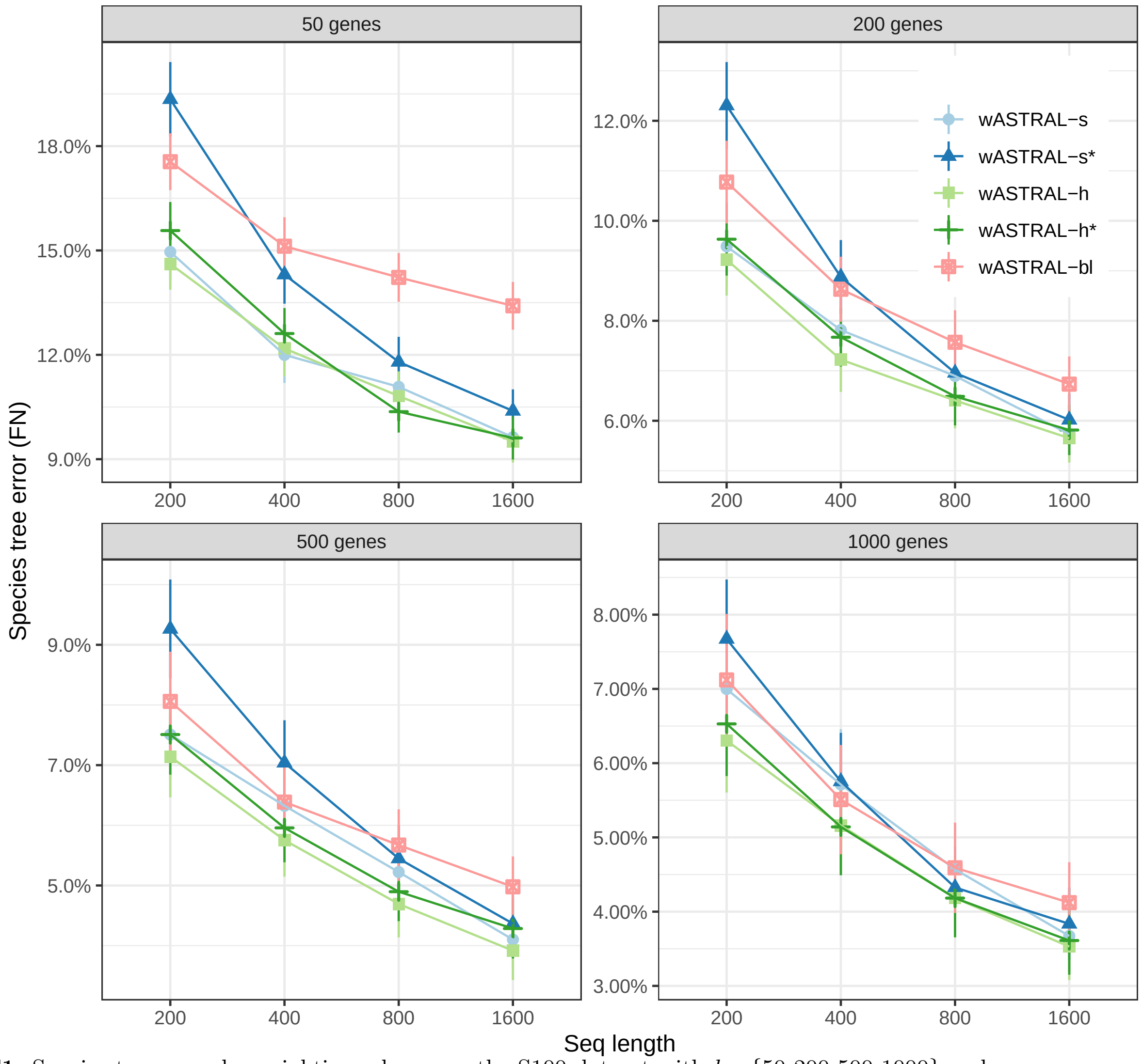


**FIG. S1.** Species tree error by weighting scheme on the S100 dataset with $k=\{50,200,500,1000\}$ and gene sequence length $\{200,400,800,1600\}$. Results with aBayes supports are labeled wASTRAL-s and wASTRAL-h; results with bootstrap support are labelled wASTRAL-s* and wASTRAL-h*.

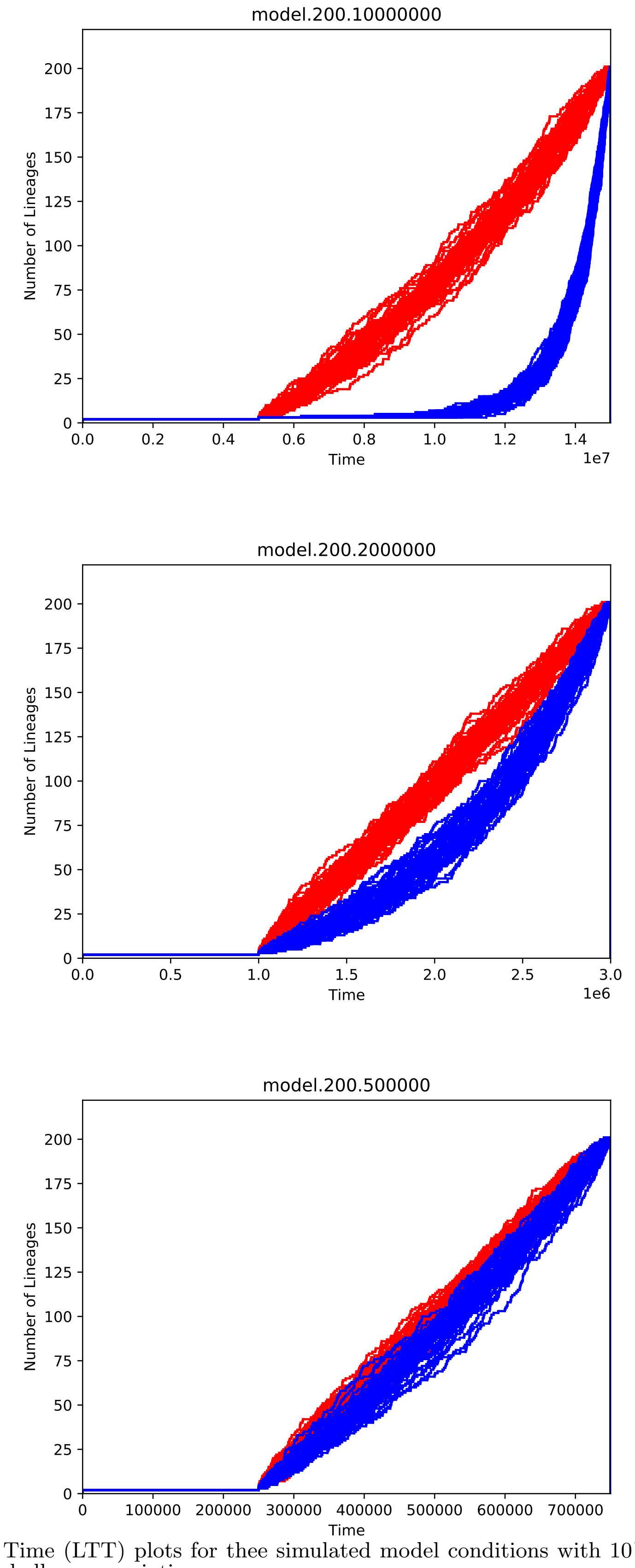


**FIG. S2.** Lineage Through Time (LTT) plots for thee simulated model conditions with $10^{-7}$ (red) and $10^{-6}$ (blue) rates tend to lead to deeper and shallower speciation.

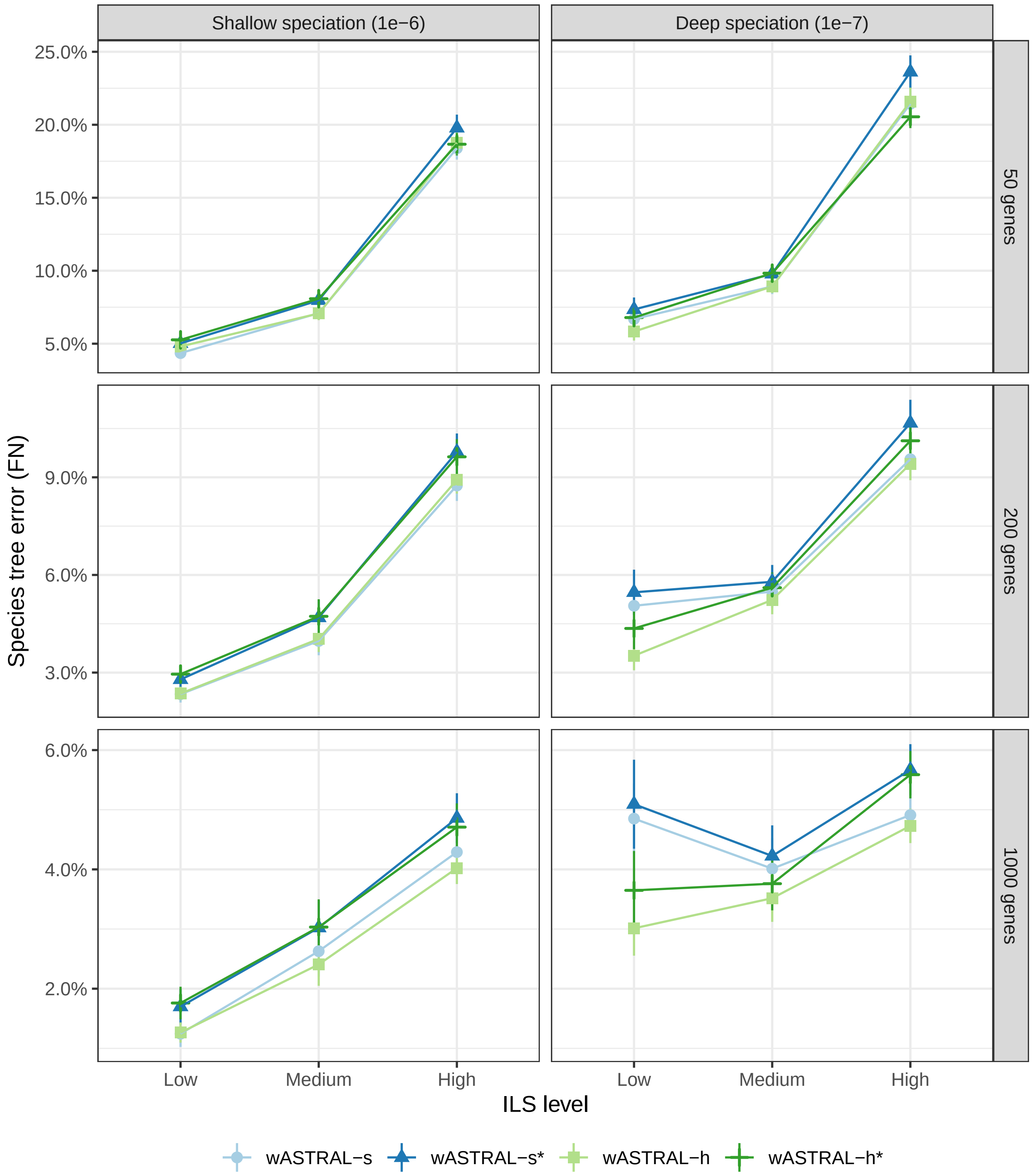


**FIG. S3.** Species tree error by weighting scheme on the S200 dataset with $k=\{50,200,1000\}$ and population size (ILS levels). Species tree shape with parameters E1-6 and E1-7 are used. Results with aBayes supports are labeled wASTRAL-s and wASTRAL-h; results with SH-like support are labeled wASTRAL-s* and wASTRAL-h*.

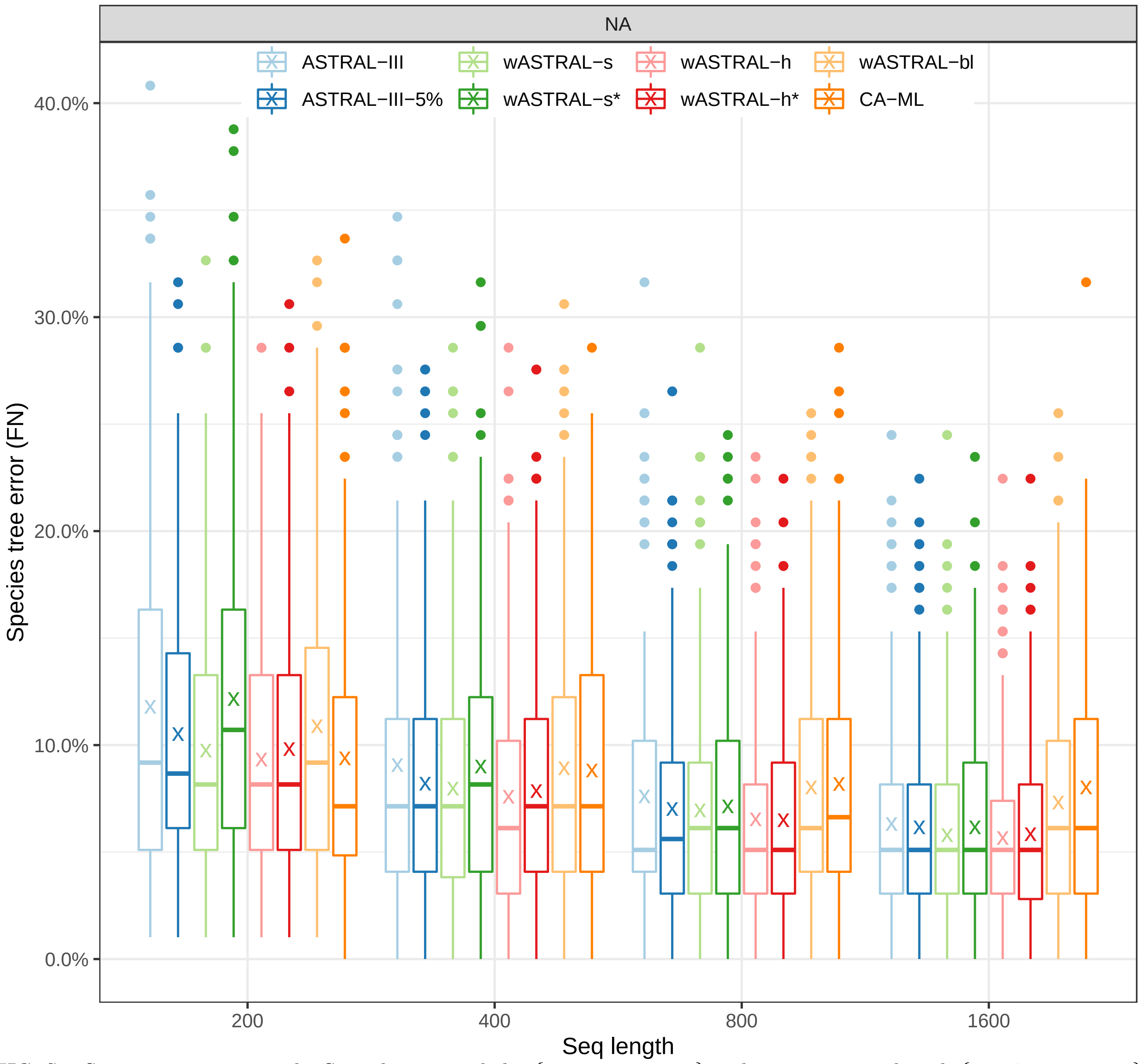


**FIG. S4.** Species tree error on the S100 dataset with $k=\{50,200,500,1000\}$ and gene sequence length $\{200,400,800,1600\}$. Results with aBayes supports are labelled wASTRAL-s and wASTRAL-h; results with bootstrap support are labelled wASTRAL-s* and wASTRAL-h*.

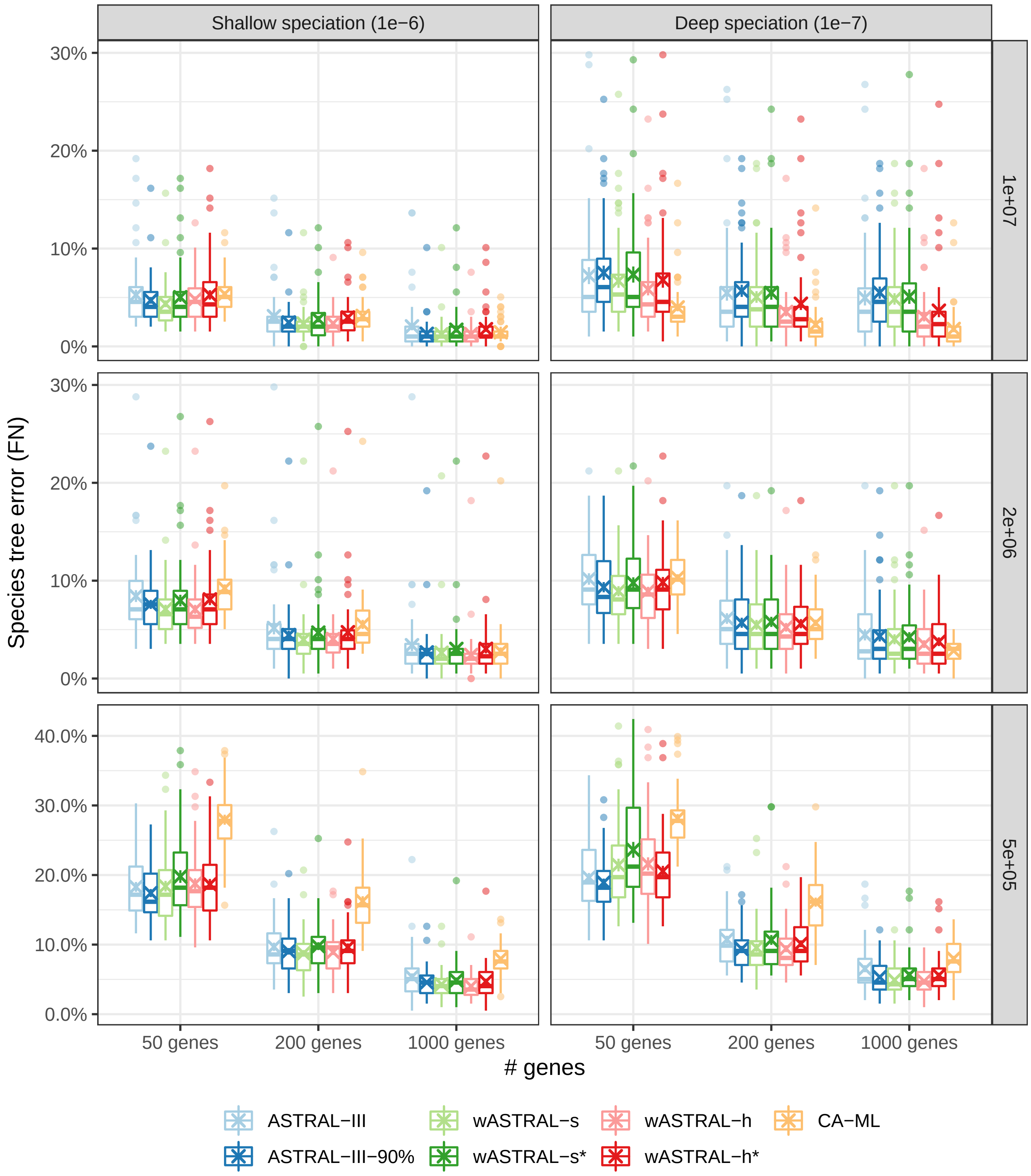


**FIG. S5.** Species tree error on the S200 dataset with $k = \{50, 200, 1000\}$ and population size (ILS levels). Species tree shape with parameter E1-6 and E1-7 (box columns) and ILS levels (box rows) low (1e+07), medium (2e+06), and high (5e+05) are used. Results with Bayesian supports are labeled wASTRAL-s and wASTRAL-h; results with SH-like support are labeled wASTRAL-s* and wASTRAL-h*.

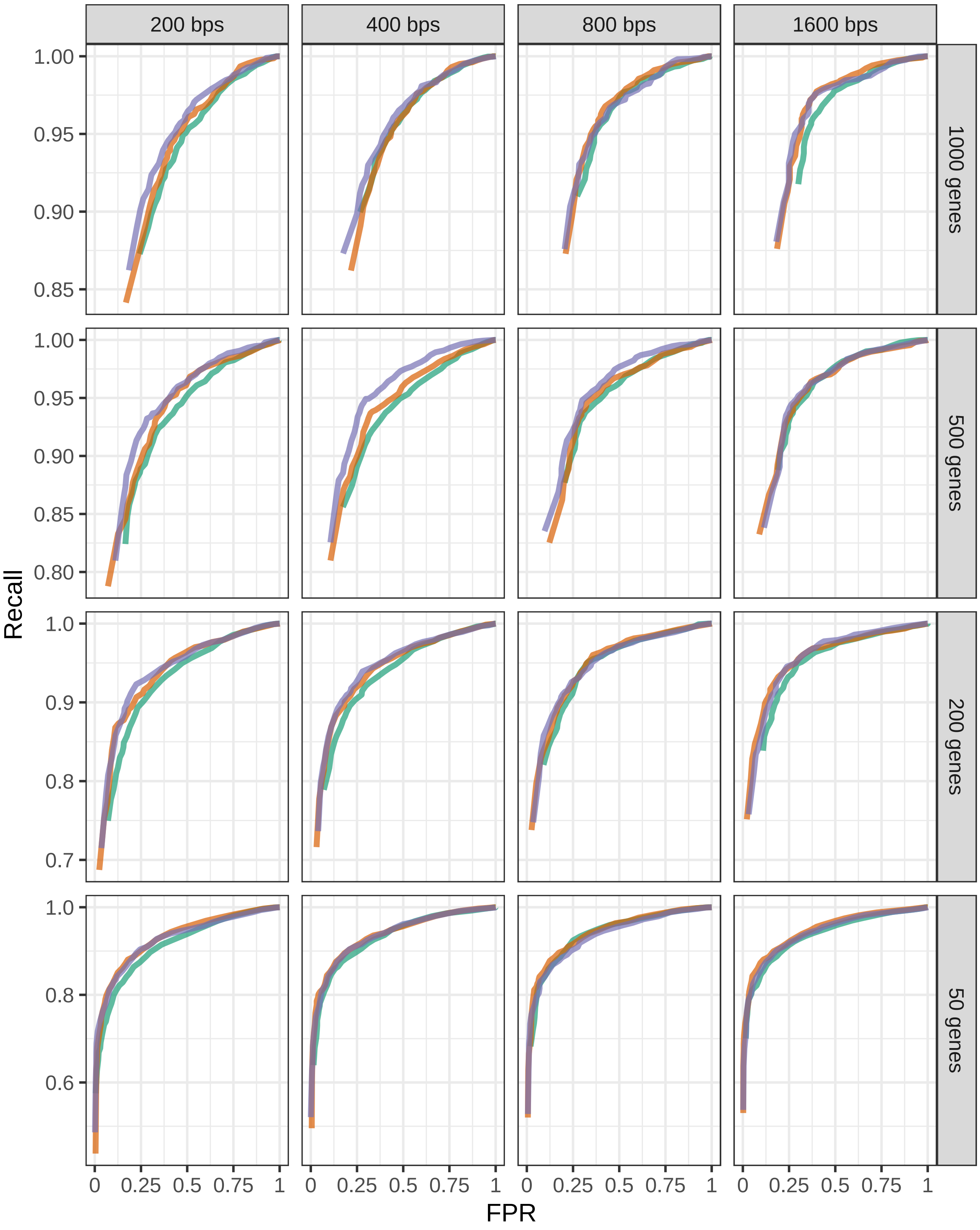


**FIG. S6.** ROC of S100 dataset with $k = \{50, 200, 500, 1000\}$ and gene sequence length $\{200, 400, 800, 1600\}$ as we change the threshold of support considered. Results with aBayes supports are labelled wASTRAL-s and wASTRAL-h; results with FastTree-2 bootstrap support are labelled wASTRAL-s* and wASTRAL-h*.

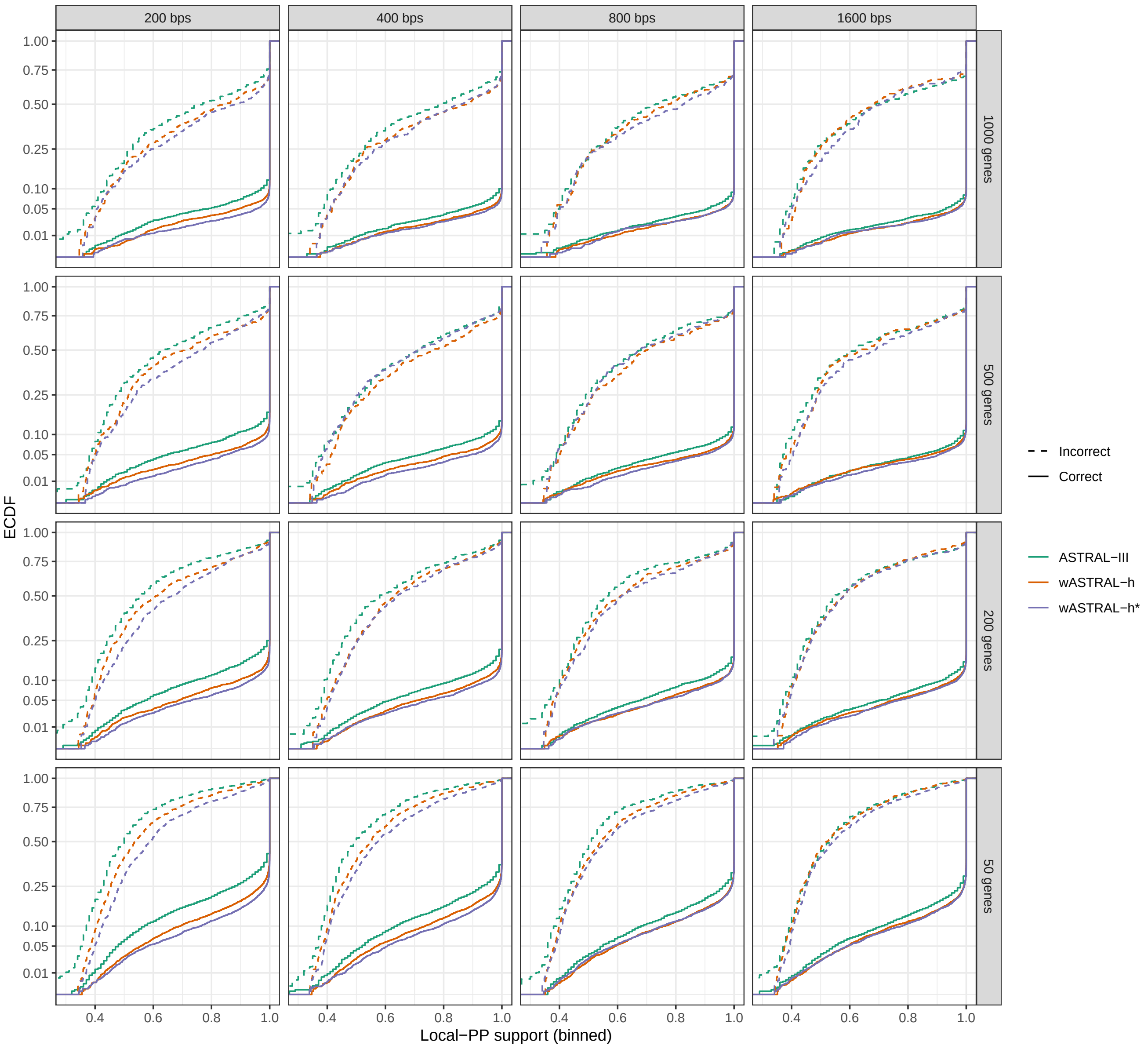


**FIG. S7.** ECDF of S100 dataset with $k = \{50,200,500,1000\}$ and gene sequence length $\{200,400,800,1600\}$. Results with aBayes supports are labelled wASTRAL-s and wASTRAL-h; results with FastTree-2 bootstrap support are labelled wASTRAL-s* and wASTRAL-h*.

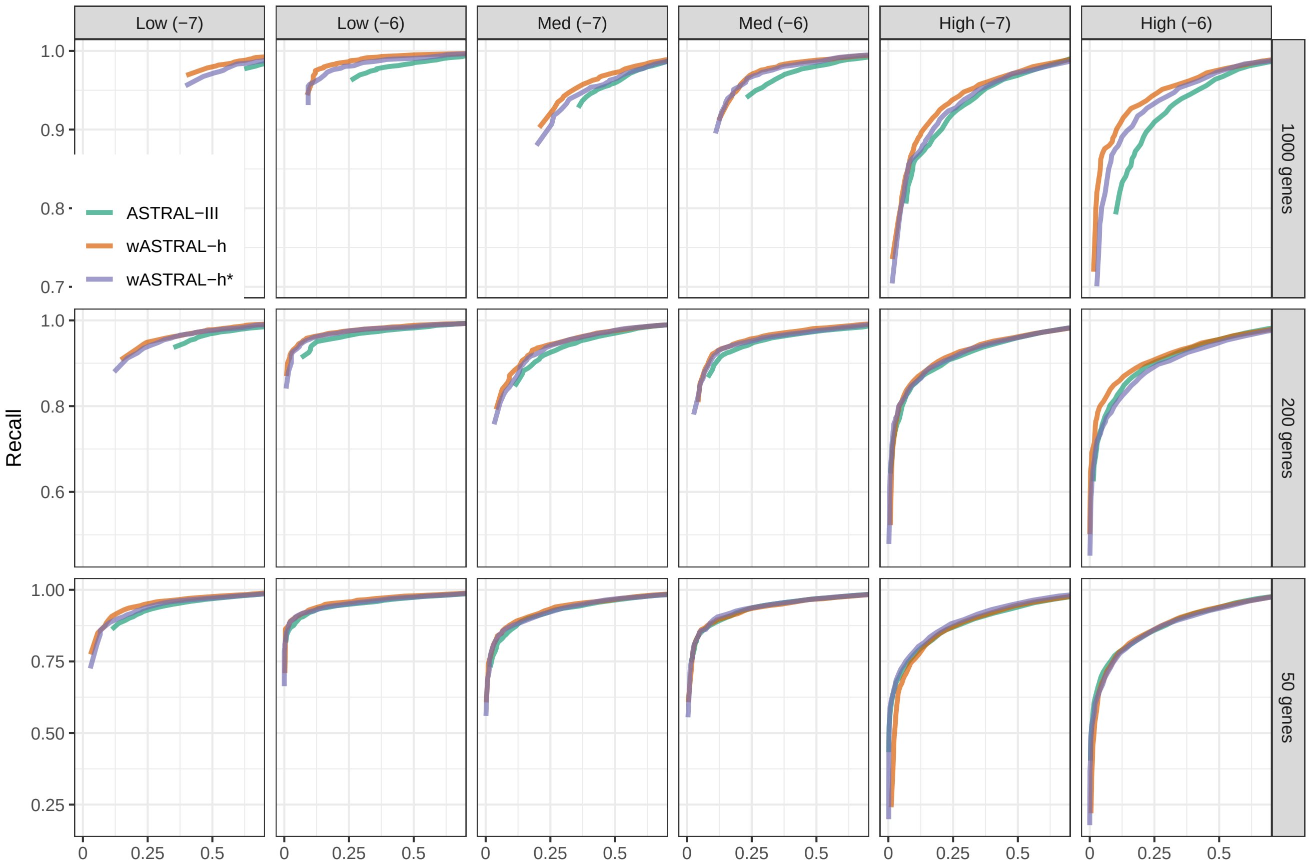

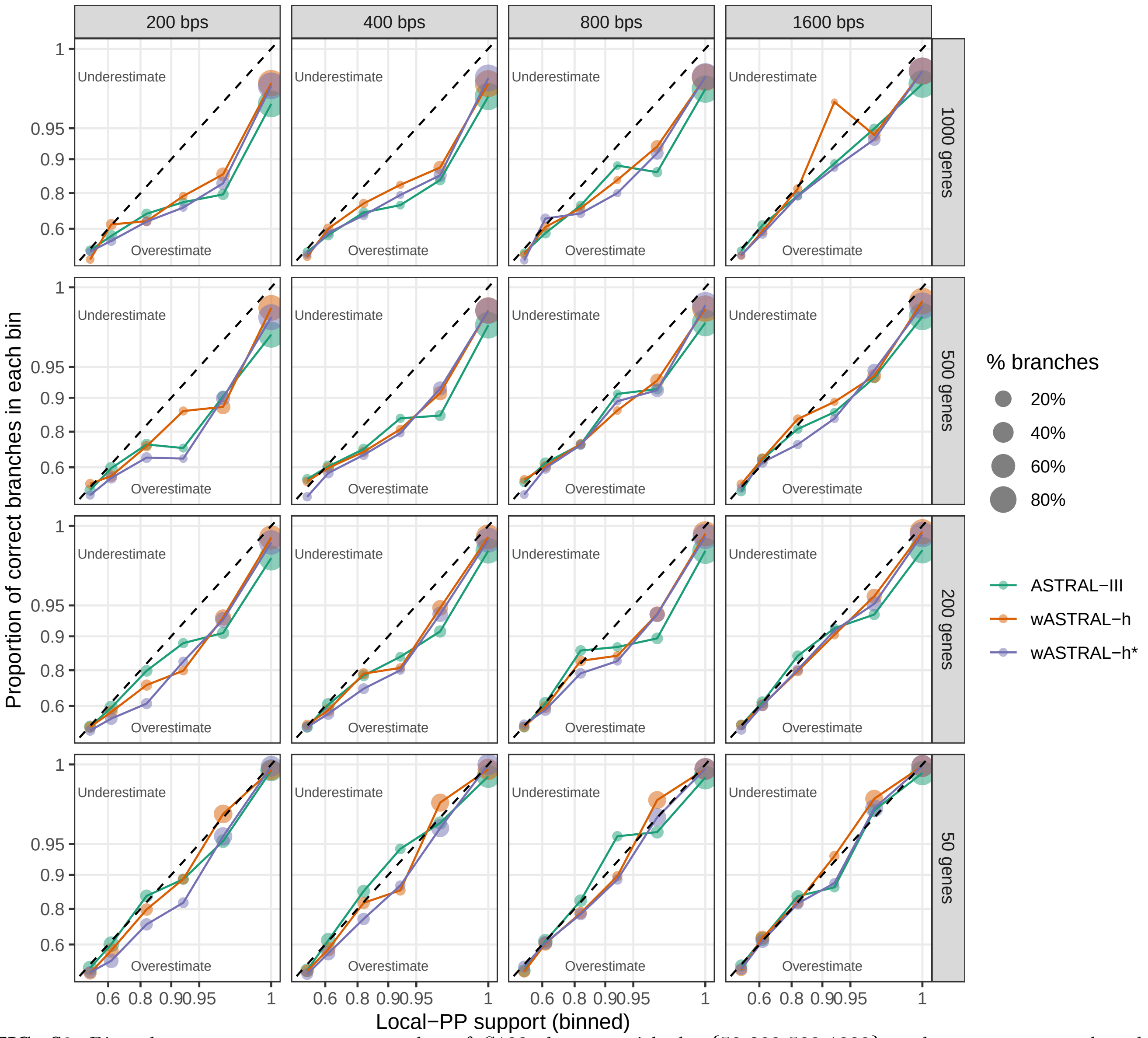


**FIG. S8.** Binned accuracy-verses-support plot of S100 dataset with $k$={50,200,500,1000} and gene sequence length {200,400,800,1600}. Results with aBayes supports are labelled wASTRAL-s and wASTRAL-h; results with FastTree-2 bootstrap support are labelled wASTRAL-s* and wASTRAL-h*.

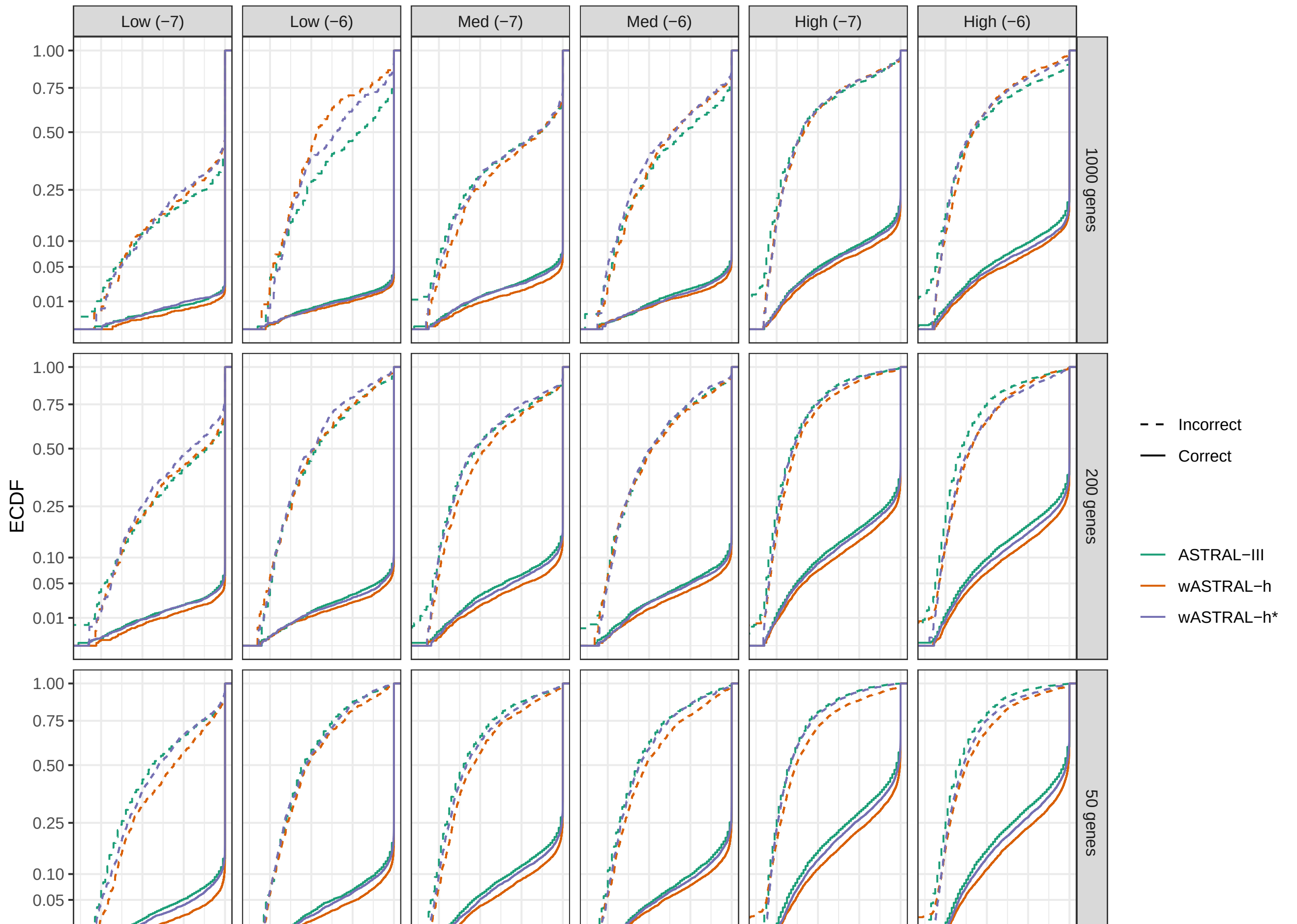

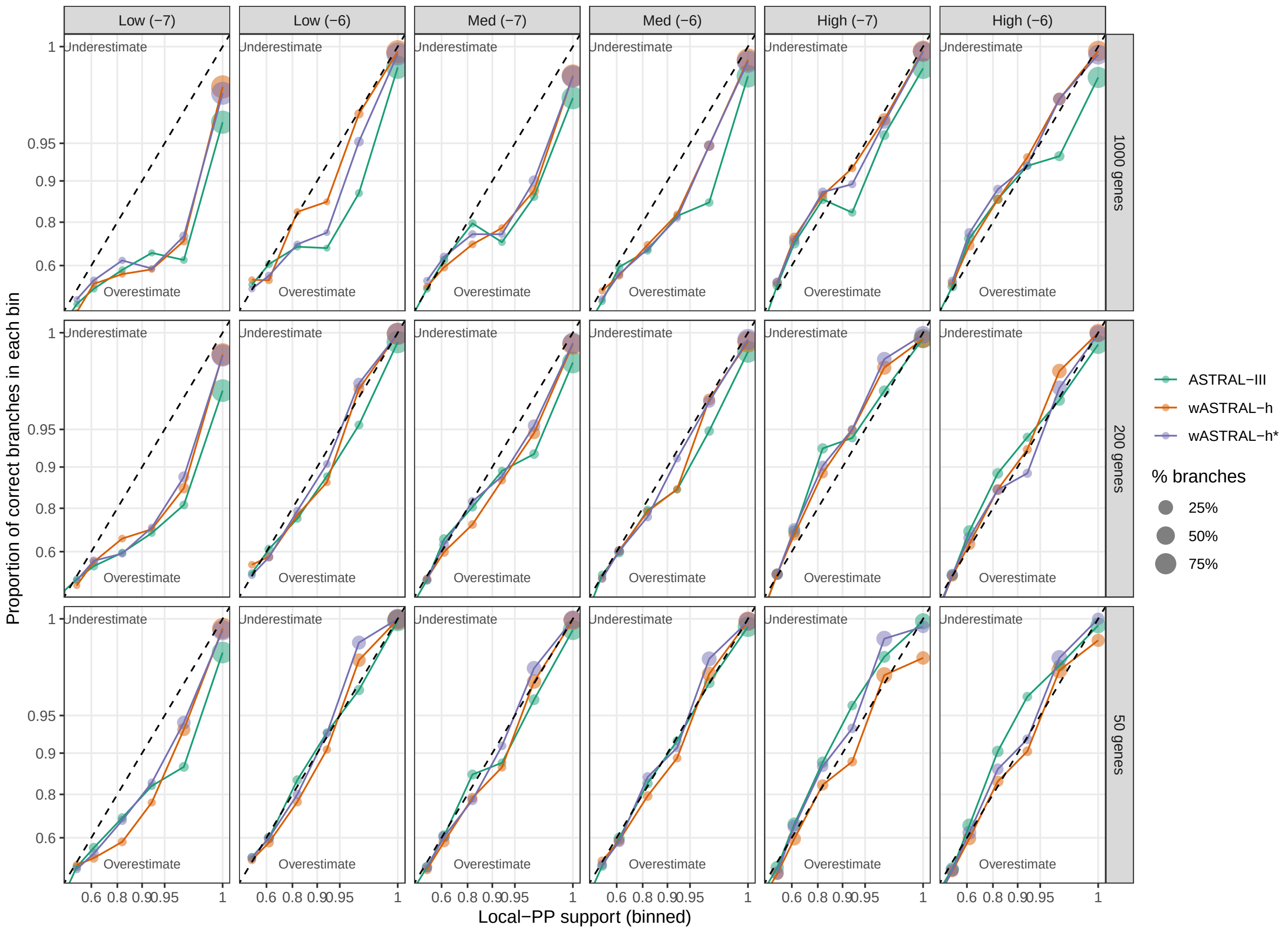


**FIG. S11.** Binned accuracy-verses-support plot of S200 dataset with $k = \{50, 200, 1000\}$ and population size (ILS levels). Species tree shape with parameter E1-6 and E1-7 are used. Results with aBayes supports are labeled wASTRAL-h; results with SH-like support are labeled wASTRAL-h*.

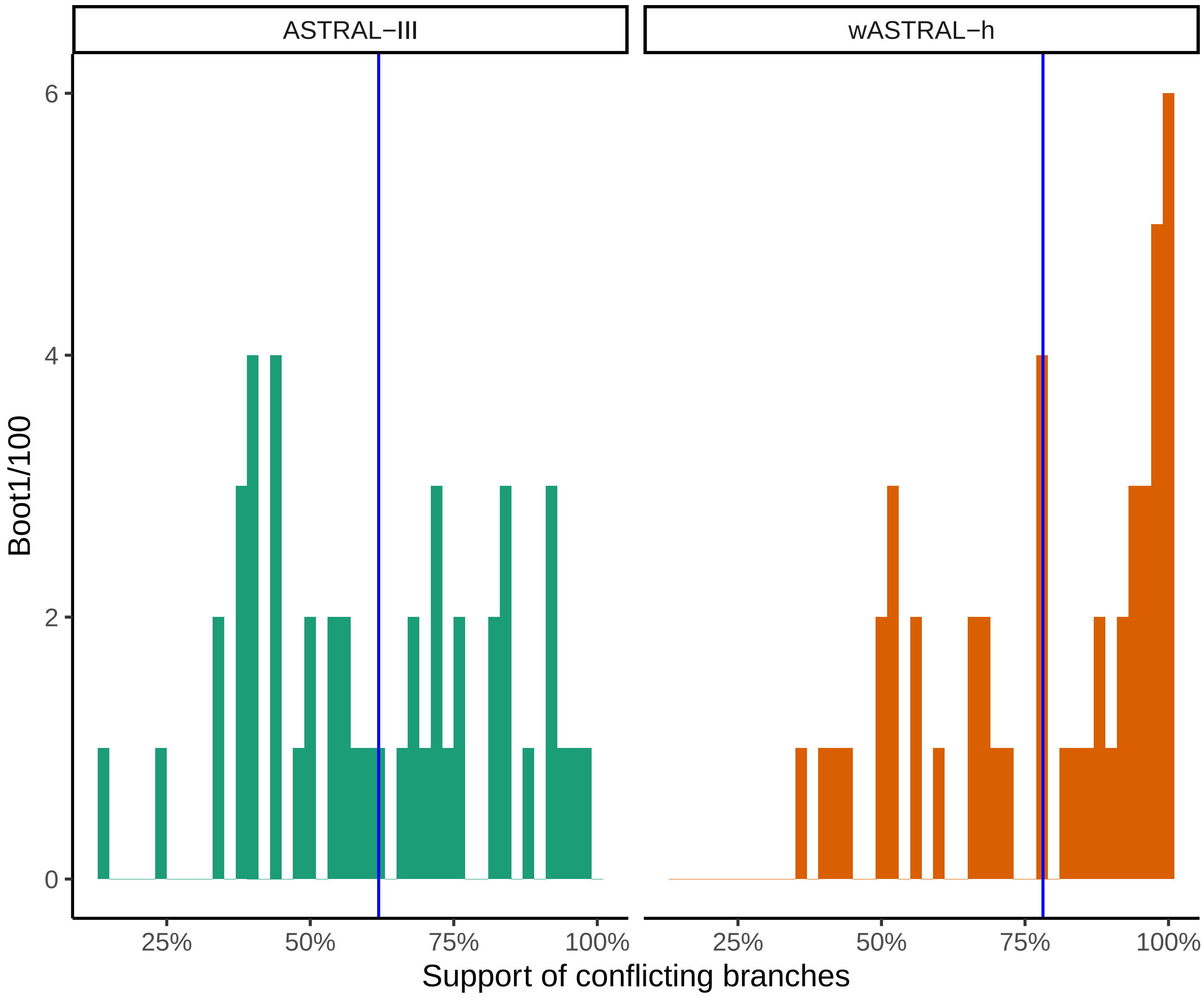


**FIG. S12.** The distribution of support values of conflicting branches between wASTRAL-h and ASTRAL-III on the 1kp dataset. The ASTRAL-III conflicting branches range between 14% and 99.00% with a mean of 62%. The wASTRAL-h conflicting branches range between 37% and 99.98% with a mean of 78%.

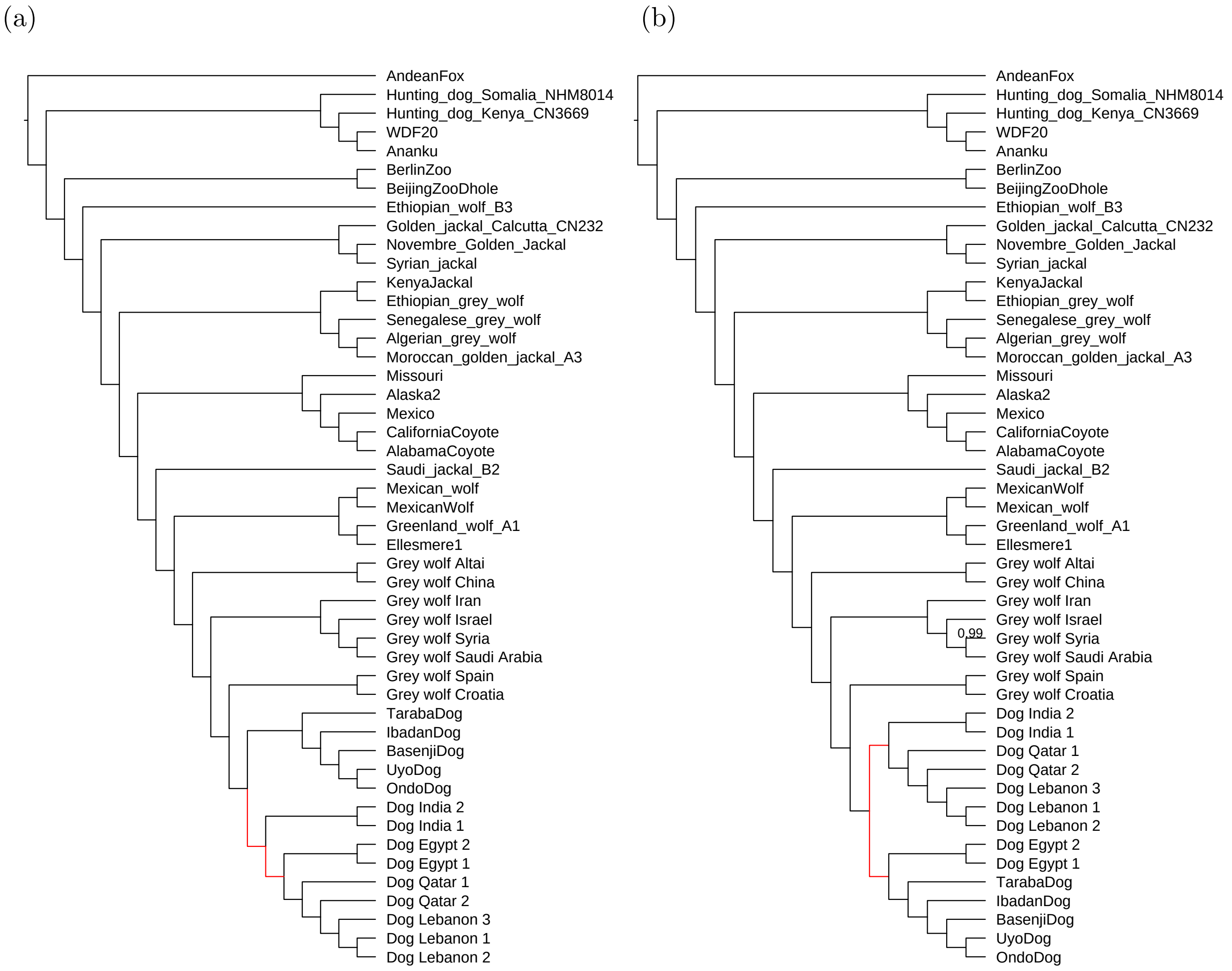


**FIG. S13.** Inferred species trees (a) from wASTRAL-hybrid with FastTree-2 branch support values as weights using all 459,450 gene trees and (b) from ASTRAL-III using a subset of 100,000 gene trees on canis dataset. Branches support of 100% are omitted.

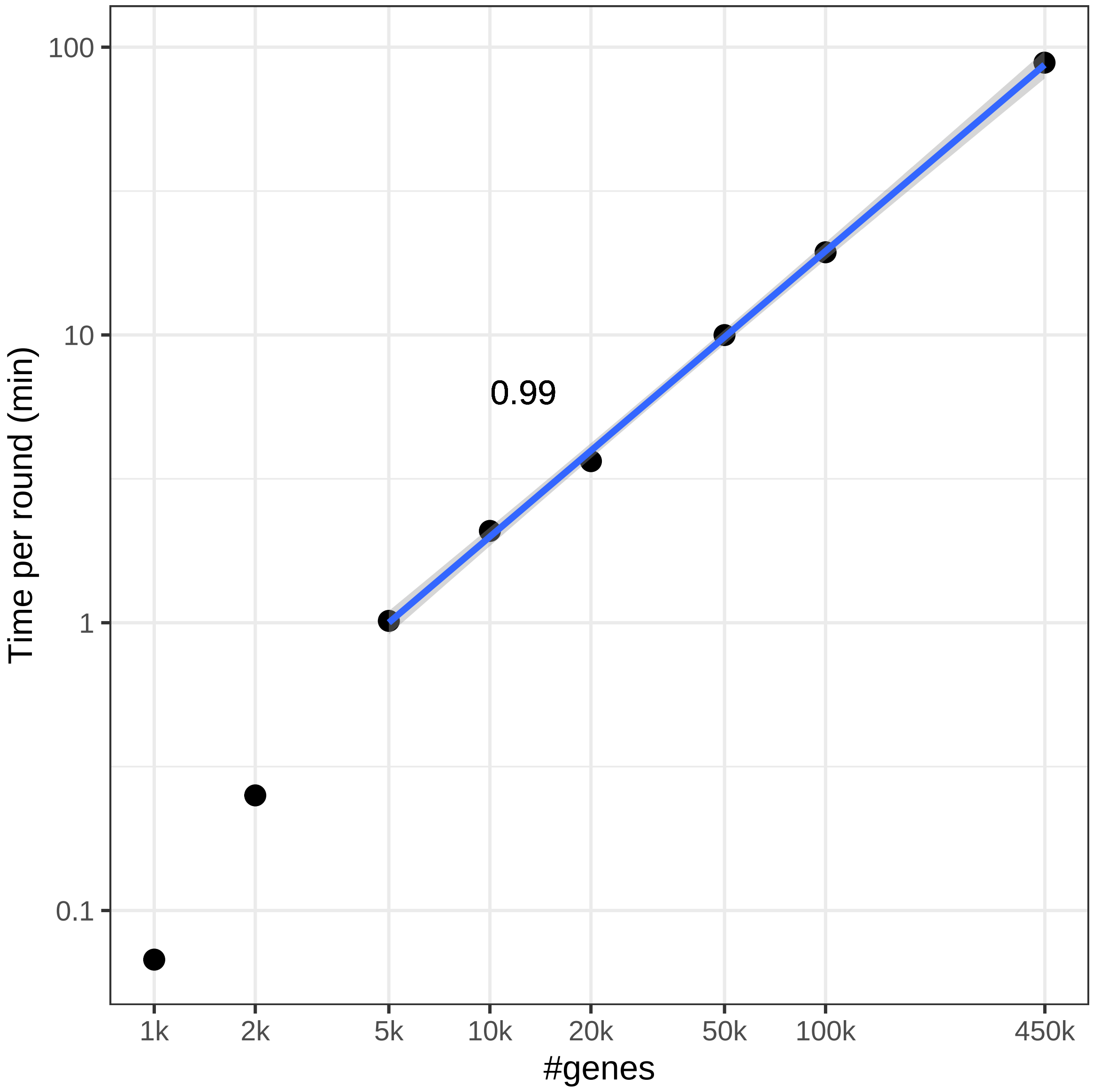


**FIG. S14.** Normalized time per round of placement by dividing running time by the total number of rounds of placements for ASTRAL on the Canis dataset for various $k$ using the new pipeline.

(a) (b)

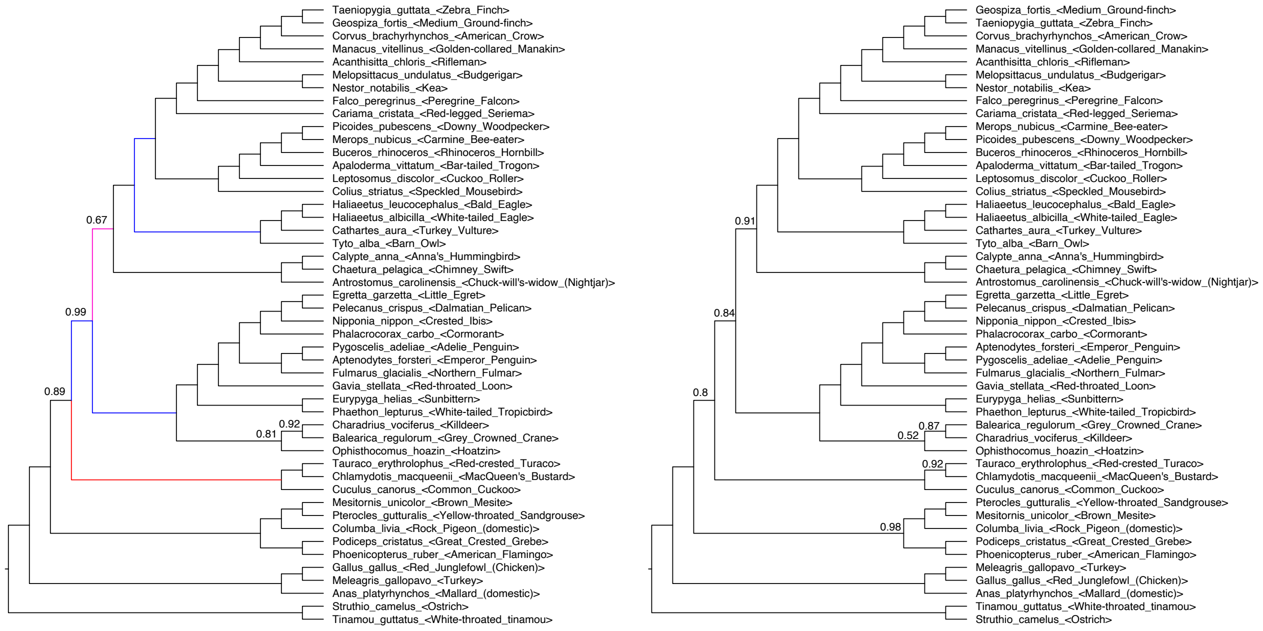


**FIG. S15.** Inferred species trees from (a) wASTRAL-hybrid with normalized bootstrap support values as weights and (b) ASTRAL-III on gene trees with low (<3% bootstrap) support branches contracted on avian dataset. Branches support of 100% are omitted. Branches that disagree with concatenation (blue), MP-EST binned (red) or both (purple) are identified on the wASTRAL-h tree.

(a) (b)

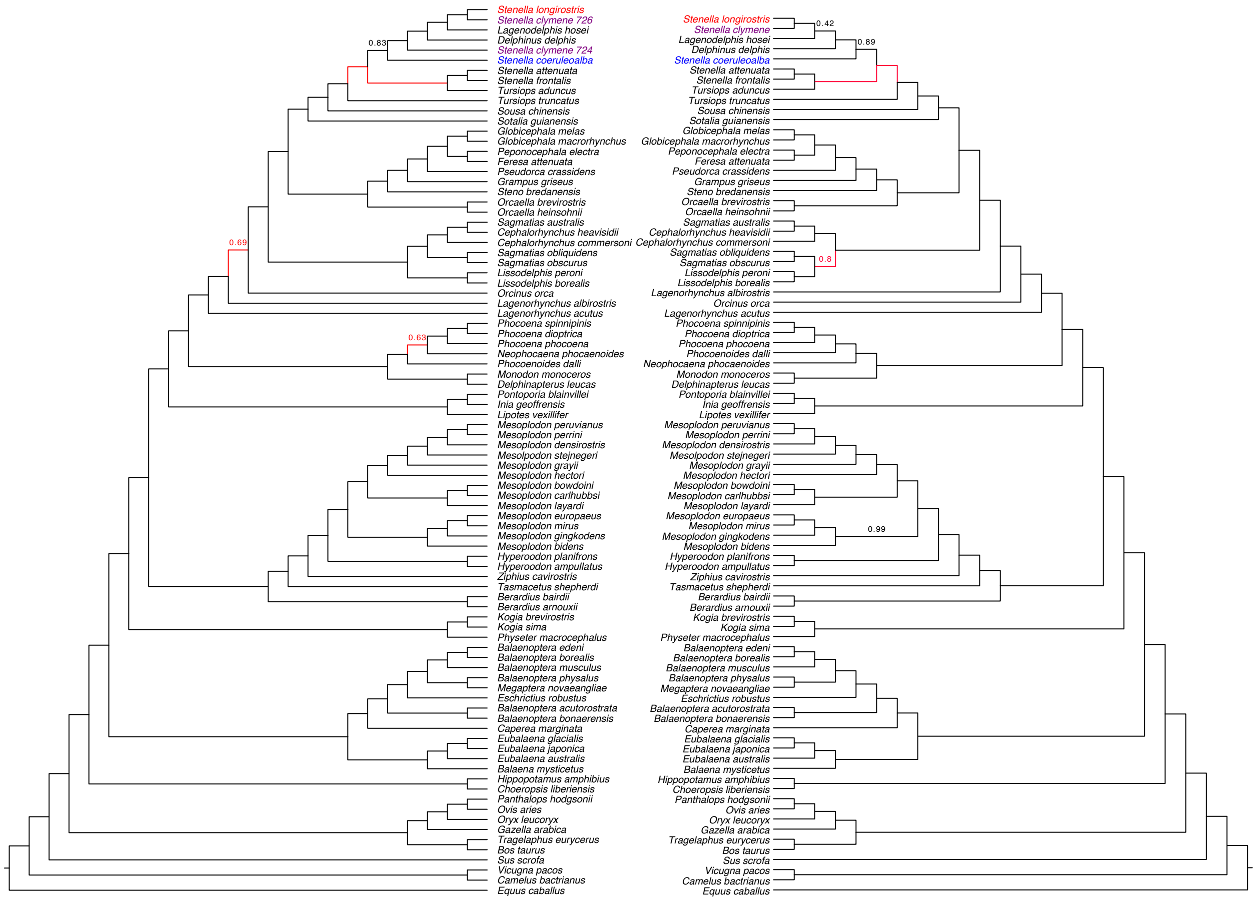


**FIG. S16.** Inferred species trees from (a) wASTRAL-hybrid with normalized Bayesian support values as weights (with clades of taxa from the same species contracted) and (b) ASTRAL-multi on cetacean dataset. Branches support of 100% are omitted. Branches conflicting with RAxML concatenation are marked red.

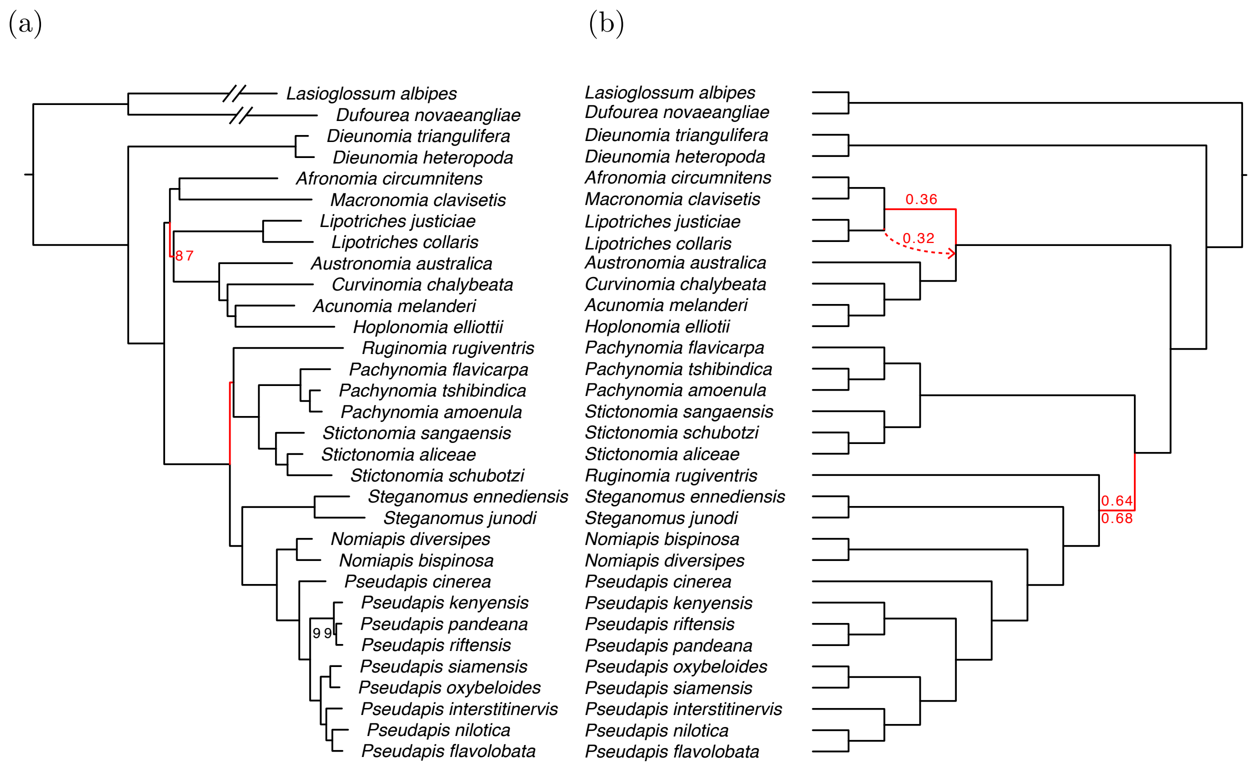


**FIG. S17.** (a) RAxML on concatenated genes; (b) wASTRAL-hybrid (top and solid red line) and ASTRAL-III (bottom and dashed red line) on Nomiinae dataset.

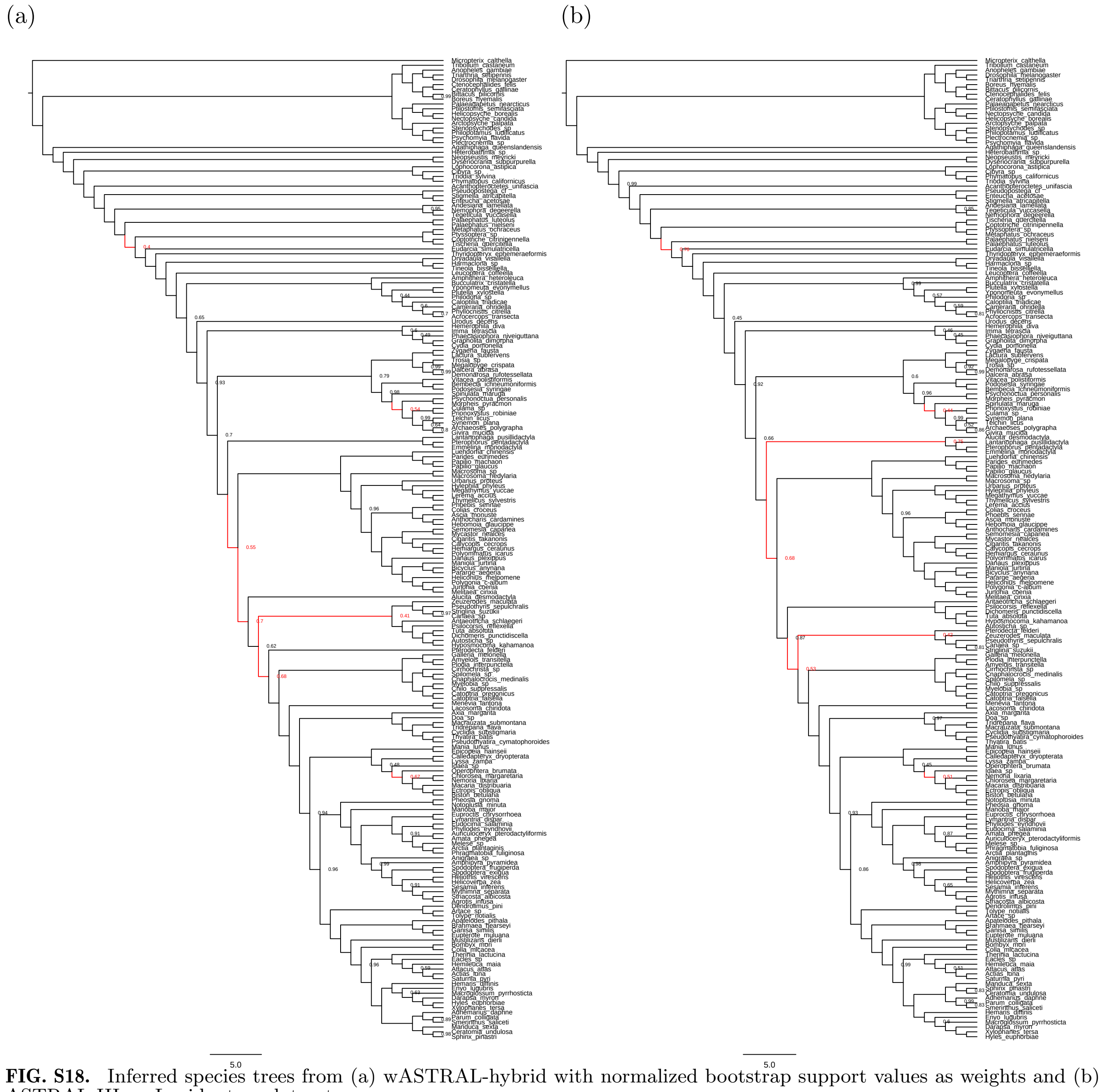

**FIG. S18.** Inferred species trees from (a) wASTRAL-hybrid with normalized bootstrap support values as weights and (b) ASTRAL-III on Lepidoptera dataset.

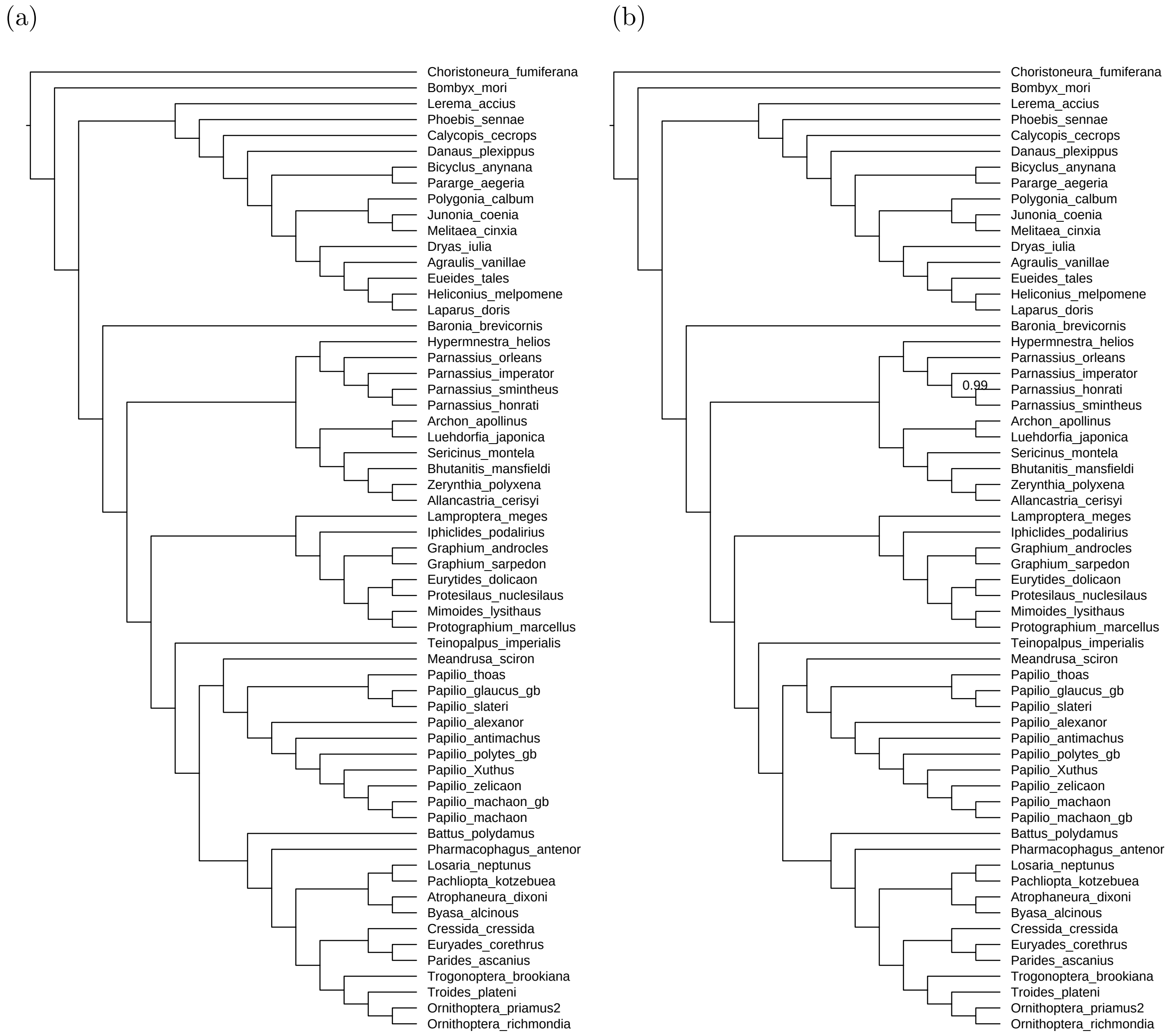


**FIG. S19.** Inferred species trees from (a) wASTRAL-hybrid with normalized approximate Bayesian support values as weights and (b) ASTRAL-III on Papilionidae dataset.

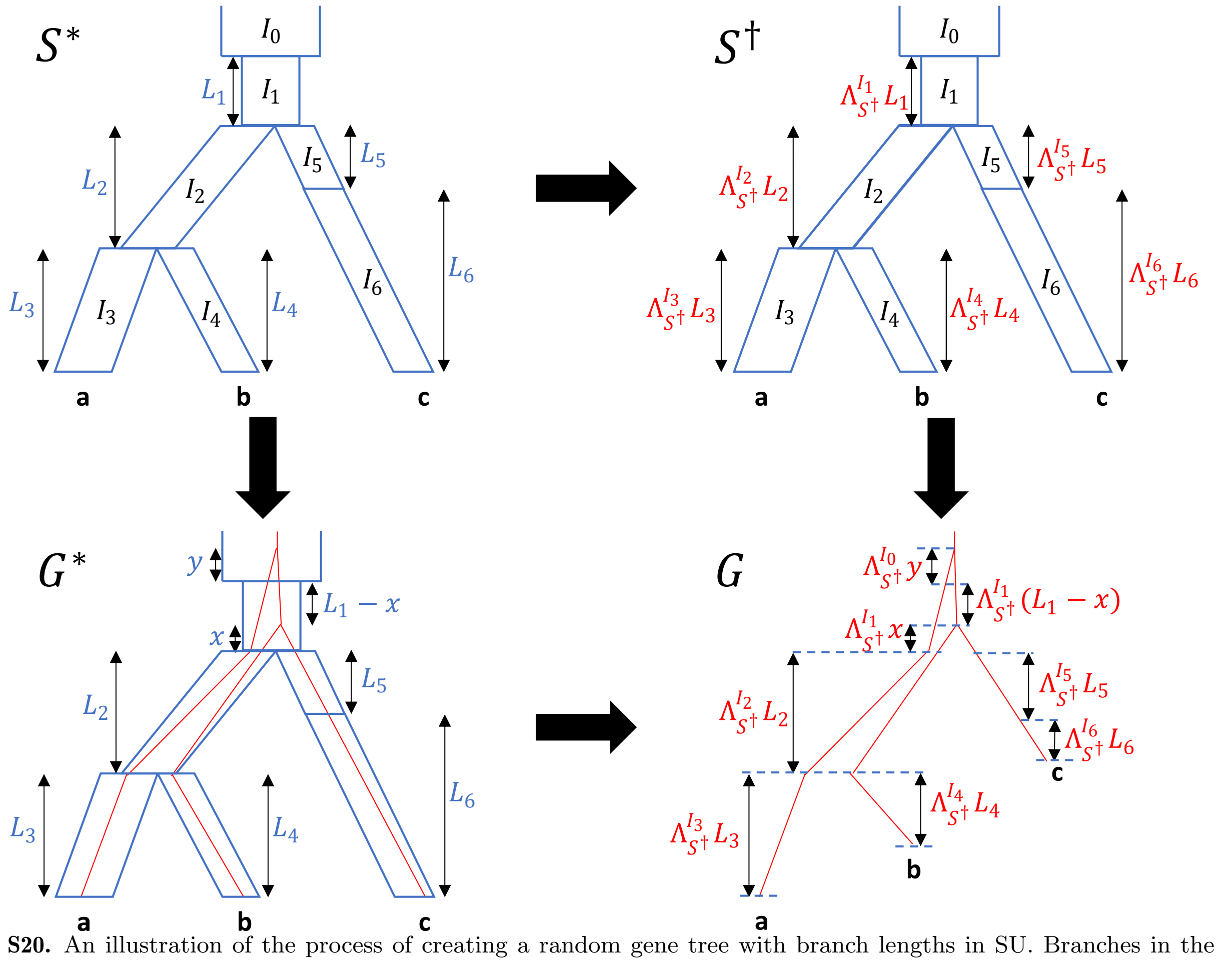


**FIG. S20.** An illustration of the process of creating a random gene tree with branch lengths in SU. Branches in the true species tree $S^*$ are broken into intervals $I_0 \ldots I_6$. The species tree with SU branch lengths $S^\dagger$ is created by multiplying each branch length in $S^*$ with a corresponding multiplier; the multipliers are jointly drawn from some distribution and are drawn independently across gene trees. Gene tree $G^*$ is sampled under MSC process from $S^*$ independent of $S^\dagger$. However, it inherits the same division of its lineages into segments as $S^*$ at the same locations. The gene tree with SU branch lengths $G$ is created by translating branch lengths of $G^*$ into SU by multiplying the CU length of each of segment $I_i$ by $\Lambda^{I_i}_{S^\dagger}$, the multiplier associated with the segment $I_i$ in $S^\dagger$ and hence $G$.

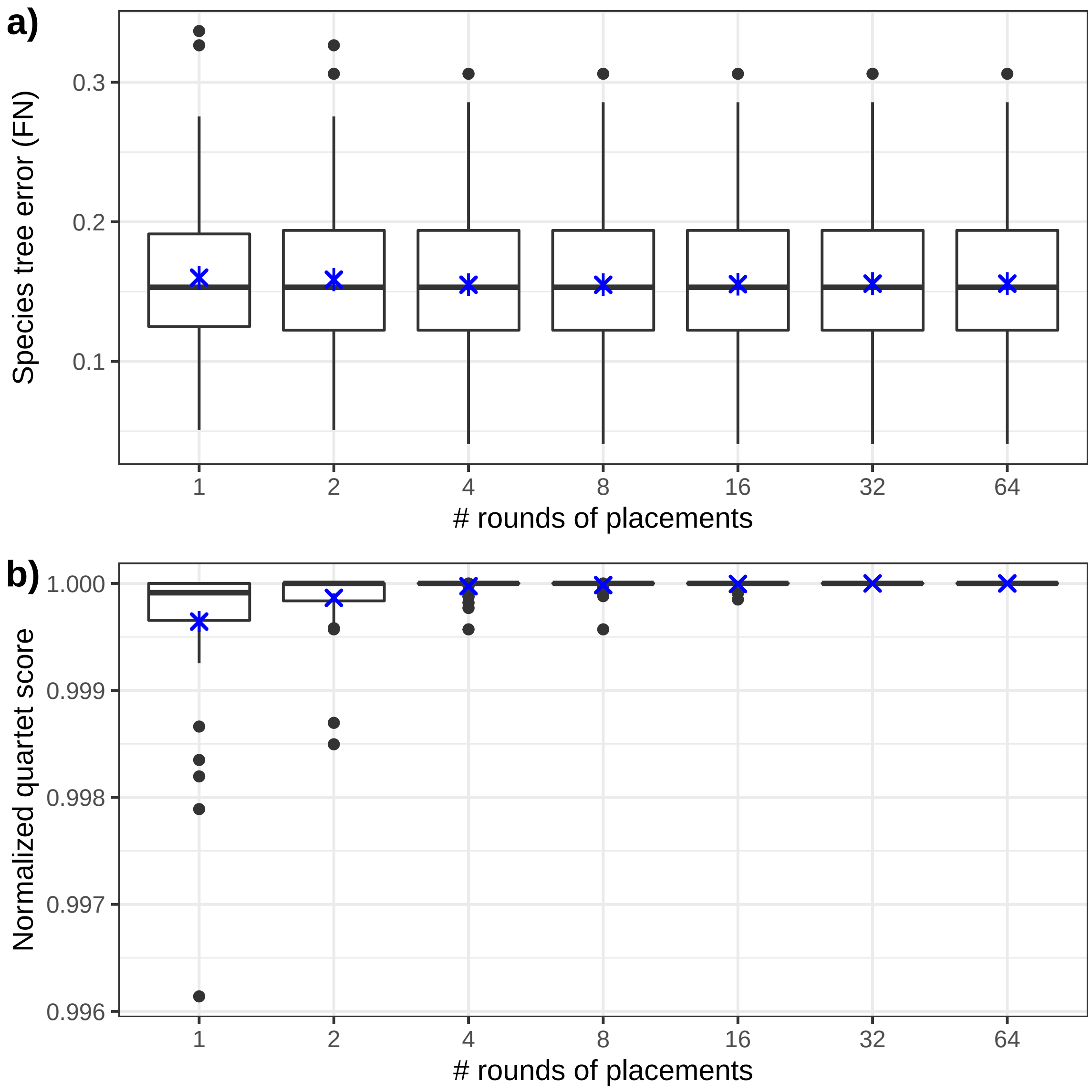


**FIG. S21.** The species tree estimation error (FN) of wASTRAL-h on S100 dataset as we change the number of rounds of placements in the base algorithm ($r$). The most difficult case where gene length=200 and $k$=50 is selected. Mean and standard error (50 replicates) are shown in blue.

**Supplementary Algorithm**

**Algorithm S1** Recursive placement algorithm. `Place` inserts the species $i$ into an existing species tree $S$ and computes tripartition scores $W(A|B|C,\mathcal{G}) := \sum_{G\in\mathcal{G}} W(A\cap\mathcal{L}_G|B\cap\mathcal{L}_G|C\cap\mathcal{L}_G,G)$ for all tripartitions resulting from adding $i$ onto each branch of $S$. A global counter $Q$ and a set of per-node counters $w_A, w_B, w_C, w^+_{..}, w^-_{..}, w_{..|.}, w_{..|..}$ are all initialized to 0. `OptimalTreeDP` is defined in Algorithm S2. Each gene tree is rooted on an arbitrary branch $e$ and the support of $e$ is kept for the branch on one side of the root and zero support is given to the branch on the other side of root. $\mathcal{L}_v$ is the set of leaves under $v$.

**procedure** PLACE($i,S,\mathcal{G}$) ▷ Places species $i$ on tree $S$ according to $\mathcal{G}$
    $W \leftarrow$ empty lookup table ▷ global variables
    COLORLEAFSET($\mathcal{L}_S$,C,$\emptyset,\mathcal{G},W$) ▷ Color all leaves of $S$ as $C$
    COLORLEAFSET($\{i\}$,B,$\emptyset,\mathcal{G},W$) ▷ Color new species $i$ as $B$
    COLORNODE(the root of $S,i,S,\mathcal{G},W$) ▷ Traverse $S$ bottom up
    $O \leftarrow$ OPTIMALTREEDP($\mathcal{L}_S\cup\{i\},\mathcal{L}_S\cup\{i\},W$)
    **return** ($W,O$, edge of $S$ onto which $i$ is added to get $O$)
**procedure** COLORLEAFSET($\mathcal{L}^*,X,T,\mathcal{G},W$) ▷ Condition: Coloring $\mathcal{L}^*$ as $X$ should match $T$
    **for** $G\in\mathcal{G}$ **do**
        **for** $j\in\mathcal{L}^*\cap\mathcal{L}_G$ **do**
            $W[T] \leftarrow$ UPDATECOUNTERS(leaf node corresponding to $j$ in $g$, $X$)
**procedure** COLORNODE($w,i,S,\mathcal{G},W$) ▷ On start: $i$ is $B$, others are $C$; On exit: $w$ is $A$, others kept
    **if** $w$ is a leaf **then**
        COLORLEAFSET($\mathcal{L}_w$,A,$\mathcal{L}_w|\{i\}|\mathcal{L}_S-\mathcal{L}_w,\mathcal{G},W$)
    **else**
        $(u,v) :=$ ( the larger child of $w$, the smaller child of $w$ )
        COLORNODE($v,i,S,\mathcal{G},W$) ▷ Recurse on $v$, the smaller child
        COLORLEAFSET($\mathcal{L}_v$,C,$\emptyset,\mathcal{G},W$) ▷ Undo coloring of $v$ to enable recursing on $u$.
        COLORNODE($u,i,S,\mathcal{G},W$) ▷ Recurse on $u$, the large child
        COLORLEAFSET($\mathcal{L}_v$,B,$\mathcal{L}_u|\{i\}\cup\mathcal{L}_v|\mathcal{L}_S-\mathcal{L}_w,\mathcal{G},W$) ▷ Tripartition of $w$ when adding $i$ above $v$
        COLORLEAFSET($\{i\}$,A,$\{i\}\cup\mathcal{L}_u|\mathcal{L}_v|\mathcal{L}_S-\mathcal{L}_w,\mathcal{G},W$) ▷ Tripartition of $w$ when adding $i$ above $u$
        COLORLEAFSET($\{i\}$,C,$\mathcal{L}_u|\mathcal{L}_v|\{i\}\cup\mathcal{L}_S-\mathcal{L}_w,\mathcal{G},W$) ▷ Tripartition of $w$ when adding $i$ above $w$
        COLORLEAFSET($\{i\}$,B,$\emptyset,\mathcal{G},W$)
        COLORLEAFSET($\mathcal{L}_v$,A,$\mathcal{L}_w|\{i\}|\mathcal{L}_S-\mathcal{L}_w,\mathcal{G},W$) ▷ Tripartition of the new parent of $i$ and $w$
**procedure** RECURSIVEUPDATE($w$)
    $(u,v,e) :=$ ( the left child of $w$, the right child of $w$, the parent branch of $w$ )
    **for** $(X,Y,Z)\in\{(A,B,C),(B,C,A),(C,A,B)\}$ **do**
        $Q \leftarrow Q - w_{XX|YZ}$
        $w_{XX|YZ} \leftarrow v_X u_{YZ|X} + u_X v_{YZ|X} + u_{XX|Z} v_Y + v_{XX|Z} u_Y + u_{XX|Y} v_Z + v_{XX|Y} u_Z$
            $+ (u^+_{YZ} v^+_{XX} - u^-_{YZ} v^-_{XX}) + (u^+_{XX} v^+_{YZ} - u^-_{XX} v^-_{YZ})$
        $Q \leftarrow Q + w_{XX|YZ}$
        **if** $w$ is not the root **then**
            $(w_X, w_Y, w_Z) \leftarrow \big((u_X+v_X)e^{-l(e)}, (u_Y+v_Y)e^{-l(e)}, (u_Z+v_Z)e^{-l(e)}\big)$
            $w^+_{XX} \leftarrow u^+_{XX} + v^+_{XX} + u_X v_X$
            $w^-_{XX} \leftarrow (u^-_{XX} + v^-_{XX} + u_X v_X)\big(1-s(e)\big)$
            $w^+_{YZ} \leftarrow u^+_{YZ} + v^+_{YZ} + u_Y v_Z + u_Z v_Y$
            $w^-_{YZ} \leftarrow (u^-_{YZ} + v^-_{YZ} + u_Y v_Z + u_Z v_Y)\big(1-s(e)\big)$
            $w_{YZ|X} \leftarrow \big(u_{YZ|X} + v_{YZ|X} + (u^+_{YZ} - u^-_{YZ})v_X + u_X(v^+_{YZ} - v^-_{YZ})\big)e^{-l(e)}$
            $w_{XX|Y} \leftarrow \big(u_{XX|Y} + v_{XX|Y} + (u^+_{XX} - u^-_{XX})v_Y + u_Y(v^+_{XX} - v^-_{XX})\big)e^{-l(e)}$
            $w_{XX|Z} \leftarrow \big(u_{XX|Z} + v_{XX|Z} + (u^+_{XX} - u^-_{XX})v_Z + u_Z(v^+_{XX} - v^-_{XX})\big)e^{-l(e)}$
            RECURSIVEUPDATE(the parent of $w$)
**procedure** UPDATECOUNTERS($w,X$) ▷ $w$ is a leaf, $X$ is a color
    $e :=$ the parent branch of $w$
    $(w_A, w_B, w_C) \leftarrow (0,0,0)$
    $w_X \leftarrow e^{-l(e)}$
    RECURSIVEUPDATE(the parent of $w$)
    **return** Q

**Algorithm S2** The Algorithm S2 of $O(n^2 kH \log n)$ running time. At start, the function is called as with $\mathcal{L}_S, \mathcal{G}, r$ as input.

```
procedure NaivePlacement(T, G, r)
    W* ← empty lookup table from tripartitions to their weights
    for i ∈ {1,...,r} do
        shuffle T
        S_i ← tree with leaves T_1, T_2, and T_3
        for j ∈ {4,...,|T|} do
            W_i, S_i, e ← Place(T_j, S_i, G)
        Add all elements of W_i to W*
    return OptimalTreeDP(T, T, W*)
procedure OptimalTreeDP(P, L, W)
    if DPTree(P) available then
        return DPTree(P)
    if |P| = 1 then
        DPScore(P) ← 0
        DPTree(P) ← Singleton rooted tree with leafset P
    else
        X ← −∞
        for A ∈ {A : W[A|P−A|L−P] has been computed} do
            S_1 ← OptimalTreeDP(A, L, W)
            S_2 ← OptimalTreeDP(P−A, L, W)
            if DPScore(A) + DPScore(P−A) + W[A|P−A|L−P] > X then
                X ← DPScore(A) + DPScore(P−A) + W[A|P−A|L−P]
                DPTree(P) ← merge subtrees S_1 and S_2 at root
        DPScore(P) ← X
    return DPTree(P)
```

**Algorithm S3** The DAC algorithm of $O(n^{1.5+\epsilon}k)$ running time given some assumptions. OptimalTreeDP and NaivePlacement are defined in Algorithm S2, and Place is defined in Algorithm S1. At start, the function is called as with $\mathcal{L}_S, \mathcal{G}, r$ as input.

**procedure** TwoStepPlacement($T, \mathcal{G}, r$)
    $W^* \leftarrow$ empty lookup table from tripartitions to their weights
    **for** $i \in \{1, ..., r\}$ **do**
        $T_i \leftarrow$ a subsample of $T$ by removing each element independently with probability $1 - 1/\sqrt{|T|}$
        $S_i :=$ NaivePlacement($T_i, \mathcal{G}, \sqrt{|T|}$)
        **for** $e \in E_{S_i}$ **do**
            $C_e \leftarrow$ empty list
        **for** $j \in T - T_i$ **do**
            $W, S_o, e \leftarrow$ Place($j, S_i, \mathcal{G}$)
            add $T_j$ to $C_e$
        $C_\emptyset \leftarrow$ empty list
        $S'_i \leftarrow S_i$
        **for** $e \in$ branches of $S_i$ **do**
            $S_e \leftarrow S_i$
            **for** $j \in C_e$ **do**
                $W, S_o, e' \leftarrow$ Place($j, S_e, \mathcal{G}$)
                **if** $e' \in S_i - \{e\}$ **then**
                    add $j$ to $C_\emptyset$
                **else**
                    $S_e \leftarrow S_o$
            $S'_i \leftarrow$ The merger of compatible trees $S_e$ and $S'_i$
        **for** $j \in C_\emptyset$ **do**
            $W_i, S'_i, e \leftarrow$ Place($j, S'_i, \mathcal{G}$)
        **if** $C_\emptyset = \emptyset$ **then**
            $W_i, S'_i, e \leftarrow$ Place($\emptyset, S'_i, \mathcal{G}$)
        Add all elements of $W_i$ to $W^*$
    **return** OptimalTreeDP($T, T, W^*$)

**Proofs**

Weighting by support: Proof of Proposition 1 and Theorem 1

For ease of reference, we reproduce Table 2 from the main paper here:

| $\mathbb{E}[(\cdot)(\cdot)\vert\alpha_{G,Q}]$ | $\delta_G(ab\vert cd)$ | $\delta_G(ac\vert bd)$ |
|---|---|---|
| $\delta_{G^*}(ab\vert cd)$ | $\geq\frac{1}{3}(1+2\theta_Q)\big(\alpha_{G,Q}+\frac{1}{3}(1-\alpha_{G,Q})(1-\beta_Q)\big)$ | $\leq\frac{1}{3}(1+2\theta_Q)\big(\frac{1}{3}(1-\alpha_{G,Q})(1+\beta_Q)\big)$ |
| $\delta_{G^*}(ac\vert bd)$ | $\geq\frac{1}{3}(1-\theta_Q)\big(\frac{1}{3}(1-\alpha_{G,Q})(1-\beta_Q)\big)$ | $\leq\frac{1}{3}(1-\theta_Q)\big(\alpha_{G,Q}+\frac{1}{3}(1-\alpha_{G,Q})(1+\beta_Q)\big)$ |
| $\delta_{G^*}(ad\vert bc)$ | $\geq\frac{1}{3}(1-\theta_Q)\big(\frac{1}{3}(1-\alpha_{G,Q})(1-\beta_Q)\big)$ | $\leq\frac{1}{3}(1-\theta_Q)\big(\frac{1}{3}(1-\alpha_{G,Q})(1+\beta_Q)\big)$ |
| $\mathbb{E}[(\cdot)(\cdot)\vert\alpha_{G,Q}]$ | $w_G(ab\vert cd)$ | $w_G(ac\vert bd)$ |
| $\delta_{G^*}(ab\vert cd)$ | $\geq\frac{1}{3}(1+2\theta_Q)\big(\alpha_{G,Q}+\frac{1}{3}(1-\alpha_{G,Q})(1-\beta_Q)\big)^2$ | $\leq\frac{1}{3}(1+2\theta_Q)\big(\frac{1}{3}(1-\alpha_{G,Q})(1+\beta_Q)\big)^2$ |
| $\delta_{G^*}(ac\vert bd)$ | $\geq\frac{1}{3}(1-\theta_Q)\big(\frac{1}{3}(1-\alpha_{G,Q})(1-\beta_Q)\big)^2$ | $\leq\frac{1}{3}(1-\theta_Q)\big(\alpha_{G,Q}+\frac{1}{3}(1-\alpha_{G,Q})(1+\beta_Q)\big)^2$ |
| $\delta_{G^*}(ad\vert bc)$ | $\geq\frac{1}{3}(1-\theta_Q)\big(\frac{1}{3}(1-\alpha_{G,Q})(1-\beta_Q)\big)^2$ | $\leq\frac{1}{3}(1-\theta_Q)\big(\frac{1}{3}(1-\alpha_{G,Q})(1+\beta_Q)\big)^2$ |

Recall that the expected value and variance of $\alpha_{G,Q}$ across genes is denoted by $\bar{\alpha_Q}$ and $\sigma_\alpha^2$.

Proposition 1. *For each estimated gene tree $G$, $\mathbb{E}\big[\delta_G(ab|cd)-\delta_G(ac|bd)\big]\geq\theta_Q\bar{\alpha_Q}-\frac{2}{3}(1-\bar{\alpha_Q})\beta_Q$ and $\mathbb{E}\big[w_G(ab|cd)-w_G(ac|bd)\big]\geq\frac{1}{9}\theta_Q(3+2\beta_Q)(\bar{\alpha_Q}^2+\sigma_\alpha^2)+\frac{2}{9}(3-\beta_Q)\theta_Q\bar{\alpha_Q}-\frac{4}{9}(1-\bar{\alpha_Q})\beta_Q$.*

*Proof.* To prove the Proposition, we start with the following lemma.

Lemma 1. *For each estimated gene tree $G$ with a given $\alpha_{G,Q}$,*

$$\mathbb{E}\big[\delta_G(ab|cd)-\delta_G(ac|bd)\big|\alpha_{G,Q}\big]\geq\theta_Q\alpha_{G,Q}-\frac{2}{3}(1-\alpha_{G,Q})\beta_Q$$

*and*

$$\mathbb{E}\big[w_G(ab|cd)-w_G(ac|bd)\big|\alpha_{G,Q}\big]\geq\frac{1}{9}(3\alpha_{G,Q}-2\beta_Q+2\alpha_{G,Q}\beta_Q+6)\theta_Q\alpha_{G,Q}-\frac{4}{9}(1-\alpha_{G,Q})\beta_Q\,.$$

*Proof.* From Table 2, we can compute

$$\begin{aligned}
&\mathbb{E}\big[\delta_G(ab|cd)-\delta_G(ac|bd)\big|\alpha_{G,Q}\big]\\
=&\mathbb{E}\Big[\big(\delta_G(ab|cd)-\delta_G(ac|bd)\big)\big(\delta_{G^*}(ab|cd)+\delta_{G^*}(ac|bd)+\delta_{G^*}(ad|bc)\big)\Big|\alpha_{G,Q}\Big]\\
\geq&\frac{1}{3}\big((1+2\theta_Q)\alpha_{G,Q}+\frac{1}{3}(1-\alpha_{G,Q})(1-\beta_Q)\big)-\frac{1}{3}\big((1-\theta_Q)\alpha_{G,Q}+\frac{1}{3}(1-\alpha_{G,Q})(1+\beta_Q)\big)\\
=&\theta_Q\alpha_{G,Q}-\frac{2}{3}(1-\alpha_{G,Q})\beta_Q\,;
\end{aligned}$$

similarly,

$$\begin{aligned}
&\mathbb{E}\big[w_G(ab|cd)-w_G(ac|bd)\big|\alpha_{G,Q}\big]\\
=&\mathbb{E}\Big[\big(w_G(ab|cd)-w_G(ac|bd)\big)\big(\delta_{G^*}(ab|cd)+\delta_{G^*}(ac|bd)+\delta_{G^*}(ad|bc)\big)\Big|\alpha_{G,Q}\Big]\\
\geq&\frac{1}{3}(1+2\theta_Q)\alpha_{G,Q}\big(\alpha_{G,Q}+\frac{2}{3}(1-\alpha_{G,Q})(1-\beta_Q)\big)+\big(\frac{1}{3}(1-\alpha_{G,Q})(1-\beta_Q)\big)^2\\
&-\frac{1}{3}(1-\theta_Q)\alpha_{G,Q}\big(\alpha_{G,Q}+\frac{2}{3}(1-\alpha_{G,Q})(1+\beta_Q)\big)-\big(\frac{1}{3}(1-\alpha_{G,Q})(1+\beta_Q)\big)^2\\
\geq&\frac{1}{9}(3\alpha_{G,Q}-2\beta_Q+2\alpha_{G,Q}\beta_Q+6)\theta_Q\alpha_{G,Q}-\frac{4}{9}(1-\alpha_{G,Q})\beta_Q\,.
\end{aligned}$$

□

From this lemma, we can prove the proposition. First, assume $\alpha_{G,Q}$ is drawn from a discrete distribution. Then,

$$\begin{aligned}\mathbb{E}\big[\delta_G(ab|cd)-\delta_G(ac|bd)\big] &= \sum_{\alpha_{G,Q}}\mathbb{E}\big[\delta_G(ab|cd)-\delta_G(ac|bd)\big|\alpha_{G,Q}\big]\mathrm{P}(\alpha_{G,Q})\\ &\geq \sum_{\alpha_{G,Q}}\big(\theta_Q\alpha_{G,Q}-\frac{2}{3}(1-\alpha_{G,Q})\beta_Q\big)\mathrm{P}(\alpha_{G,Q}) = \theta_Q\bar{\alpha_Q}-\frac{2}{3}(1-\bar{\alpha_Q})\beta_Q\end{aligned}$$

and

$$\begin{aligned}&\mathbb{E}\big[w_G(ab|cd)-w_G(ac|bd)\big] = \sum_{\alpha_{G,Q}}\mathbb{E}\big[w_G(ab|cd)-w_G(ac|bd)\big|\alpha_{G,Q}\big]\mathrm{P}(\alpha_{G,Q})\\ &\geq \sum_{\alpha_{G,Q}}\big(\frac{1}{9}(3\alpha_{G,Q}-2\beta_Q+2\alpha_{G,Q}\beta_Q+6)\theta_Q\alpha_{G,Q}-\frac{4}{9}(1-\alpha_{G,Q})\beta_Q\big)\mathrm{P}(\alpha_{G,Q})\\ &=\frac{1}{9}\theta_Q(3+2\beta_Q)\mathbb{E}[\alpha_{G,Q}^2]+\frac{2}{9}(3-\beta_Q)\theta_Q\bar{\alpha_Q}-\frac{4}{9}(1-\bar{\alpha_Q})\beta_Q\\ &=\frac{1}{9}\theta_Q(3+2\beta_Q)(\bar{\alpha_Q}^2+\sigma_\alpha^2)+\frac{2}{9}(3-\beta_Q)\theta_Q\bar{\alpha_Q}-\frac{4}{9}(1-\bar{\alpha_Q})\beta_Q\,.\end{aligned}$$

It is straightforward to change these calculations to use integral instead of sum and $\mathrm{P}(\alpha_{G,Q})$ to the PDF in the case that the distribution of $\alpha_{G,Q}$ is continuous. □

---

THEOREM 1. *Given estimated gene trees furnished with support generated under MSC+Error+Support model, there exist conditions where (3) guarantee a statistically consistent estimator of $S^*$ but (2) does not, and the reverse is not true.*

*Proof.* Recall that (1) states

$$W(S,\mathcal{G}) := \sum_{G\in\mathcal{G}}\sum_{Q\in\mathcal{Q}(S)} w_G(S\upharpoonright Q)\,.$$

It means that in order to produce a statistically consistent estimator using (1), the following equation must be satisfied for the true species tree topology $S^*$ and any species tree topology $S$:

$$\mathbb{E}\big[W(S^*,\mathcal{G})-W(S,\mathcal{G})\big] = |\mathcal{G}|\sum_{Q\in\mathcal{Q}(S)}\mathbb{E}\big[w_G(S^*\upharpoonright Q)-w_G(S\upharpoonright Q)\big]\geq 0 \tag{9}$$

Notice that proving for any quartet $Q=\{a,b,c,d\}$ we have $\mathbb{E}\big[w_G(ab|cd)-w_G(ac|bd)\big]\geq 0$ and $\mathbb{E}\big[w_G(ab|cd)-w_G(ad|bc)\big]\geq 0$ where $S^*\upharpoonright Q=ab|cd$ is sufficient to prove (9); on the other hand, proving for any quartet $Q=\{a,b,c,d\}$ where the internal branch of $S^*\upharpoonright Q$ corresponds to only one branch in $S^*$, we have $\mathbb{E}\big[w_G(ab|cd)-w_G(ac|bd)\big]\geq 0$ and $\mathbb{E}\big[w_G(ab|cd)-w_G(ad|bc)\big]\geq 0$ where $S^*\upharpoonright Q=ab|cd$ is necessary to prove (9).

Thus, from Proposition 1, we have guaranteed statistical consistency for weighted ASTRAL for support under

$$D=\bigcap_{Q\in\mathcal{Q}(S)}\big\{(\theta_Q,\bar{\alpha_Q},\sigma_\alpha,\beta_Q)\in[0,1]^4 : \frac{1}{9}\theta_Q(3+2\beta_Q)(\bar{\alpha_Q}^2+\sigma_\alpha^2)+\frac{2}{9}(3-\beta_Q)\theta_Q\bar{\alpha_Q}-\frac{4}{9}(1-\bar{\alpha_Q})\beta_Q\geq 0\big\}\,.$$

Similarly, we have guaranteed statistical consistency for unweighted ASTRAL under

$$D'=\bigcap_{Q\in\mathcal{Q}(S)}\big\{(\theta_Q,\bar{\alpha_Q},\sigma_\alpha,\beta_Q)\in[0,1]^4 : \theta_Q\bar{\alpha_Q}-\frac{2}{3}(1-\bar{\alpha_Q})\beta_Q\geq 0\big\}\,.$$

To prove Theorem 1, we only need to prove that $D'$ is a proper subset of $D$.

We can prove $D' \subseteq D$, as for any $Q$, if $(\theta_Q, \bar{\alpha_Q}, \sigma_\alpha, \beta_Q) \in [0,1]^4$ and $\theta_Q \bar{\alpha_Q} - \frac{2}{3}(1-\bar{\alpha_Q})\beta_Q \geq 0$, then

$$\frac{1}{9}\theta_Q(3+2\beta_Q)(\bar{\alpha_Q}^2+\sigma_\alpha^2)+\frac{2}{9}(3-\beta_Q)\theta_Q\bar{\alpha_Q}-\frac{4}{9}(1-\bar{\alpha_Q})\beta_Q$$
$$=\frac{1}{9}\theta_Q(3+2\beta_Q)\sigma_\alpha^2+\frac{1}{3}\theta_Q(1-\theta_Q)\bar{\alpha_Q}^2+(\frac{1}{3}\theta_Q\bar{\alpha_Q}+\frac{2}{3})(\theta_Q\bar{\alpha_Q}-\frac{2}{3}(1-\bar{\alpha_Q})\beta_Q)\geq 0\,.$$

We can also prove $D' \neq D$, as if for some $Q$, $\theta_Q = 0.25, \bar{\alpha_Q} = 0.5, \beta_Q = 0.4$,

$$\theta_Q\bar{\alpha_Q}-\frac{2}{3}(1-\bar{\alpha_Q})\beta_Q=-\frac{1}{120}<0$$

and

$$\frac{1}{9}\theta_Q(3+2\beta_Q)(\bar{\alpha_Q}^2+\sigma_\alpha^2)+\frac{2}{9}(3-\beta_Q)\theta_Q\bar{\alpha_Q}-\frac{4}{9}(1-\bar{\alpha_Q})\beta_Q=\frac{7}{720}+\frac{19}{180}\sigma_\alpha^2>0\,.$$

Thus $D'$ is a proper subset of $D$ and we conclude the proof.

□

#### Weighting by length: Proof of Propositions 2 and 3 and Theorem 2

Before providing the proofs, we remind the reader of one property of the coalescent model. According to the coalescent model, at any point along a branch of the species tree with $i$ gene tree lineages, the time (i.e., distance) $x$ to the next coalescent event, reducing the number of lineages to $i-1$, is exponentially distributed with the rate $\binom{i}{2}$, resulting in probability density function (PDF):

$$\frac{i(i-1)}{2}e^{-\frac{i(i-1)}{2}x}, \tag{10}$$

and the two lineages that coalesce are independent of $x$.

Proposition 2. *For a true quartet species tree $S^*$ with topology $ab|cd$ and input gene trees $\mathcal{G}$ generated under the naive model with any multiplier $\lambda$, let $f$ be the distance between anchors of $S^*$. As $f\to 0$, given $k=\Theta(f^{-2})$ gene trees, we have $Var[X_G]=\Theta_f(1)$ and*

$$\frac{\mathbb{E}[X_G]}{\sqrt{Var[X_G]}}=\frac{1+4\lambda}{1+2\lambda}\sqrt{\frac{3}{2}}f+O(f^2).$$

*Proof.* We analyze balanced and unbalanced trees separately.

Case 1: Unbalanced trees (i.e., the root of $S^*$ has a terminal branch as a child). W.o.l.g., we assume the root branch is located on branch leading to $d$.

Let $p$,$q$, and $r$ be the MRCA nodes of $(a,b)$, $(a,c)$, and $(a,d)$ on rooted species tree $S^*$, respectively. Let $p'$ and $r'$ be the points of coalescence of leaves $a$,$b$ and leaves $c$,$d$ on the rooted gene tree $G$, respectively. Let $x$, $y_0$, and $z$ be the CU difference in heights of points $(p,p')$, $(q,r)$, and $(r,r')$, respectively. Note that $f$ is the length of $(p,q)$. Let $L:=l_{S^*}(a,p)+l_{S^*}(b,p)+l_{S^*}(c,r)+l_{S^*}(d,r)$. Notice that $l_G(a,p)+l_G(b,p)+l_G(c,r)+l_G(d,r)=\lambda L$ and $l_G(a,b)+l_G(c,d)=\lambda(2x+2z+L)$.

Let $f_X(x)$ be the probability density that $x$ is the CU difference in heights of $(p,p')$ and $p'$ is the lowest point of coalescence. Notice that by (10):

$$f_X(x)=\begin{cases} e^{-x} & 0\le x\le f\\ \frac{1}{\binom{2}{3}}\left(e^{-f}\binom{2}{3}e^{-\binom{2}{3}(x-f)}\right)=e^{-3x+2f} & f\le x\le f+y_0\\ \frac{1}{\binom{2}{4}}\left(e^{-f}e^{-\binom{2}{3}y_0}\binom{2}{4}e^{-\binom{2}{4}(x-f-y_0)}\right)=e^{-6x+5f+3y_0} & f+y_0\le x\end{cases}$$

Let $f_{Z|X}(z;x)$ be the probability density that $z$ is the CU difference in heights of $(r,r')$, conditioned on that $x$ is the CU difference in heights of $(p,p')$ and $p'$ is the lowest point of coalescence. Notice that:

$$f_{Z|X}(z;x)=\begin{cases} e^{-z} & 0\le x\le f+y_0 \text{ and } 0\le z\\ e^{-\left(z-(x-f-y_0)\right)}=e^{-z+x-f-y_0} & 0\le x-f-y_0\le z\end{cases}$$

We specify three coalescence scenarios by indicator functions $\delta_1,\delta_2,\delta_3$: *i)* $\delta_1$ indicates $0\le x<f$; *ii)* $\delta_2$ indicates $f\le x<f+y_0$; *iii)* $\delta_3$ indicates $f+y_0\le x$.

Note that

$$\mathbb{E}\left[w_G(ab|cd)\right]=\mathbb{E}\left[(\delta_1+\delta_2+\delta_3)w_G(ab|cd)\right]$$
$$\mathbb{E}\left[w_G^2(ab|cd)\right]=\mathbb{E}\left[(\delta_1+\delta_2+\delta_3)w_G^2(ab|cd)\right].$$

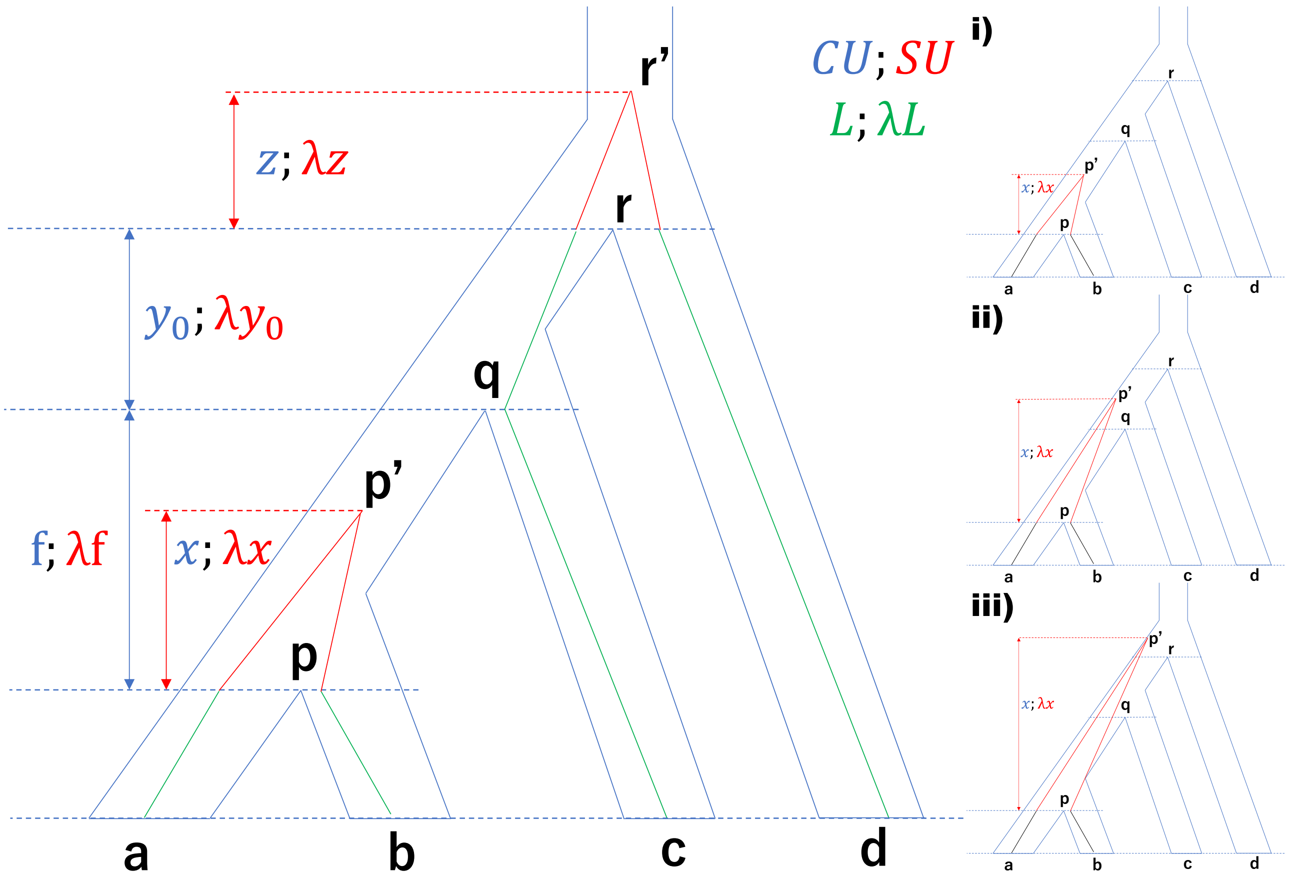


**FIG. S22.** Illustration of the unbalanced case. Lengths in CU/SU units are denoted in blue/red. Branches in green have a total length $L/\lambda L$ in CU/SU units. The right-hand side shows the position of $p'$ in relation to $q$ and $r$ in various cases.

Similarly, since only scenarios 2 and 3 have deep coalescence events that may lead to gene tree disagreement with the species tree, and by the symmetry of all three topologies under scenarios 2 and 3,

$$\mathbb{E}\big[w_G(ac|bd)\big] = \mathbb{E}\big[(\delta_2+\delta_3)w_G(ab|cd)\big]$$
$$\mathbb{E}\big[w_G^2(ac|bd)\big] = \mathbb{E}\big[(\delta_2+\delta_3)w_G^2(ab|cd)\big].$$

Thus,

$$\mathbb{E}[X_G] = \mathbb{E}\big[w_G(ab|cd)\big] - \mathbb{E}\big[w_G(ac|bd)\big] = \mathbb{E}\big[\delta_1 w_G(ab|cd)\big], \tag{11}$$

and since $w_G(ab|cd)w_G(ac|bd) = 0$,

$$\begin{aligned}\mathrm{Var}[X_G] &= \mathbb{E}[X_G^2] - \mathbb{E}^2[X_G] = \mathbb{E}\big[w_G^2(ab|cd) + w_G^2(ac|bd)\big] - \mathbb{E}^2[X_G] \\ &= \mathbb{E}\big[(\delta_1 + 2\delta_2 + 2\delta_3)w_G^2(ab|cd)\big] - \mathbb{E}^2[X_G].\end{aligned} \tag{12}$$

We next compute both elements of (11) as well as some elements of (12) (others will not be necessary).

- $\delta_1$: When $G$ has topology $ab|cd$, $p'$ must be the lowest point of coalescence. Thus,

$$\begin{aligned}&\mathbb{E}\big[\delta_1 w_G(ab|cd)\big] \\ &= \int_0^f \int_0^{+\infty} e^{-\lambda(2x+2z+L)} f_X(x) f_{Z|X}(z;x)\,dz\,dx \\ &= \int_0^f \int_0^{+\infty} e^{-\lambda(2x+2z+L)} e^{-x} e^{-z}\,dz\,dx \\ &= \frac{e^{-\lambda L}(1-e^{-(1+2\lambda)f})}{(1+2\lambda)^2};\end{aligned}$$

$$\mathbb{E}\left[\delta_1 w_G^2(ab|cd)\right] \le \mathbb{E}\left[\delta_1 w_G(ab|cd)\right] = O(f).$$

- $\delta_2$: When $G$ has topology $ab|cd$, $p'$ must be the lowest point of coalescence. Thus,

$$\begin{aligned}
&\mathbb{E}\left[\delta_2 w_G^2(ab|cd)\right]\\
&= \int_f^{f+y_0}\int_0^{+\infty} e^{-\lambda(4x+4z+2L)} f_X(x) f_{Z|Y}(z;y)\,dz\,dx\\
&= \int_f^{f+y_0}\int_0^{+\infty} e^{-\lambda(4x+4z+2L)} e^{-3x+2f} e^{-z}\,dz\,dx\\
&= \frac{1-e^{-(3+4\lambda)y_0}}{(1+4\lambda)(3+4\lambda)} e^{-(1+4\lambda)f-2\lambda L}.
\end{aligned}$$

- $\delta_3$: When $G$ has the topology $ab|cd$, either $p'$ or $q'$ must be the lowest point of coalescence, and by symmetry, the two cases must have the same PDFs. Thus,

$$\begin{aligned}
&\mathbb{E}\left[\delta_3 w_G^2(ab|cd)\right]\\
&= \int_{f+y_0}^{+\infty}\int_{x-f-y_0}^{+\infty} e^{-\lambda(4x+4z+2L)} 2 f_X(x) f_{Z|X}(z;x)\,dz\,dx\\
&= \int_{f+y_0}^{+\infty}\int_{x-f-y_0}^{+\infty} e^{-\lambda(4x+4z+2L)} 2e^{-6x+5f+3y_0} e^{-z+x-f-y_0}\,dz\,dx\\
&= \int_{f+y_0}^{+\infty} e^{-4\lambda(x+x-f-y_0)-2\lambda L} 2e^{-6x+5f+3y_0} \frac{1}{1+4\lambda}\,dx\\
&= \frac{1}{(3+4\lambda)(1+4\lambda)} e^{-(1+4\lambda)f-(3+4\lambda)y_0-2\lambda L}.
\end{aligned}$$

Replacing in (11), we get

$$\mathbb{E}[X_G] = \mathbb{E}\left[\delta_1 w_G(ab|cd)\right] = \frac{e^{-\lambda L}(1-e^{-(1+2\lambda)f})}{(1+2\lambda)^2} = \frac{e^{-\lambda L}}{1+2\lambda} f + O(f^2);$$

and replacing in (12), we get

$$\begin{aligned}
\mathrm{Var}[X_G] &= \mathbb{E}\left[(\delta_1+2\delta_2+2\delta_3) w_G^2(ab|cd)\right] - \mathbb{E}^2[X_G] = \mathbb{E}\left[2(\delta_2+\delta_3) w_G^2(ab|cd)\right] + O(f)\\
&= \frac{2e^{-(1+4\lambda)f-2\lambda L}}{(3+4\lambda)(1+4\lambda)} + O(f) = \frac{2e^{-2\lambda L}}{(3+4\lambda)(1+4\lambda)} + O(f),
\end{aligned}$$

from which our assumption of $\mathrm{Var}[X_G] = \Omega(1)$ follows.

Case 2: Balanced tree.

Let $p,q$, and $r$ be the MRCA nodes of $(a,b)$, $(c,d)$, and $(a,d)$ on rooted species tree $S^*$, respectively. Let $p'$ and $q'$ be the points of coalescence of leaves $a,b$ and leaves $c,d$ on the rooted gene tree $G$, respectively. Let $x$, $x_0$, $y$, and $y_0$ be the CU difference in heights of points $(p,p')$, $(p,r)$, $(q,q')$, and $(q,r)$, respectively. Note that $f = x+y$ is CU length of path $(p,q)$. Let $L := l_{S^*}(a,p) + l_{S^*}(b,p) + l_{S^*}(c,q) + l_{S^*}(d,q)$. Notice that $l_G(a,p) + l_G(b,p) + l_G(c,q) + l_G(d,q) = \lambda L$ and $l_G(a,b) + l_G(c,d) = \lambda(2x+2y+L)$.

We specify three coalescence scenarios by indicator functions $\delta_1,\delta_2,\delta_3$: *i)* $\delta_1$ indicates $0 \le x < x_0$; *ii)* $\delta_2$ indicates $x_0 \le x, 0 \le y < y_0$; *iii)* $\delta_3$ indicates $x_0 \le x, y_0 \le y$.

Note that

$$\mathbb{E}\left[w_G(ab|cd)\right] = \mathbb{E}\left[(\delta_1+\delta_2+\delta_3) w_G(ab|cd)\right]$$
$$\mathbb{E}\left[w_G^2(ab|cd)\right] = \mathbb{E}\left[(\delta_1+\delta_2+\delta_3) w_G^2(ab|cd)\right].$$

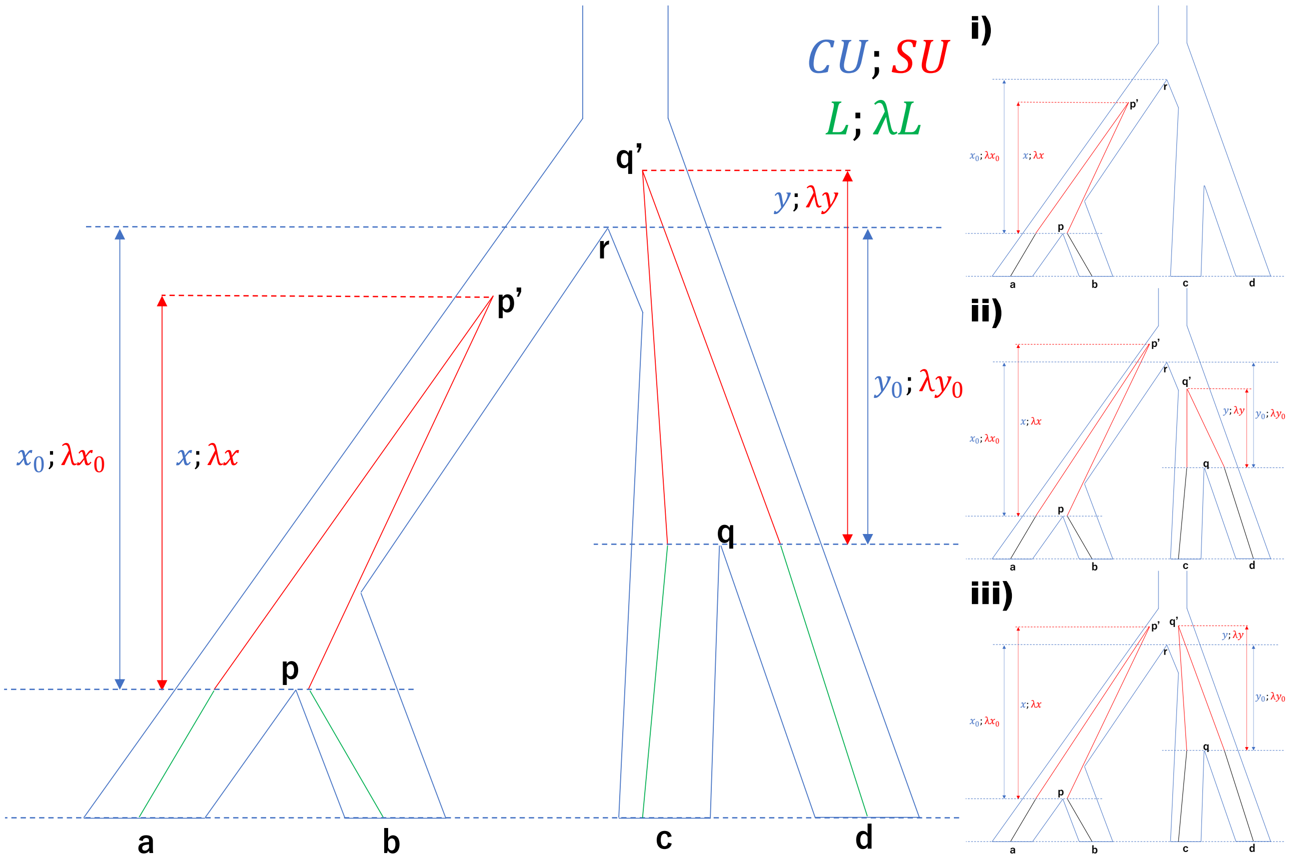


**FIG. S23.** Illustration of the unbalanced case. Lengths in CU/SU units are denoted in blue/red. Branches in green have a total length $L/\lambda L$ in CU/SU units. The right-hand side shows the position of $p'$ and $q'$ in relation to $r$ in various cases.

Similarly, since only scenarios 3 have deep coalescence events that may lead to gene tree disagreement with the species tree, and by the symmetry of all three topologies under scenarios 3,

$$\mathbb{E}\left[w_G(ac|bd)\right] = \mathbb{E}\left[\delta_3 w_G(ab|cd)\right]$$
$$\mathbb{E}\left[w_G^2(ac|bd)\right] = \mathbb{E}\left[\delta_3 w_G^2(ab|cd)\right].$$

Thus,

$$\mathbb{E}[X_G] = \mathbb{E}\left[w_G(ab|cd)\right] - \mathbb{E}\left[w_G(ac|bd)\right] = \mathbb{E}\left[(\delta_1+\delta_2) w_G(ab|cd)\right]; \tag{13}$$

and since $w_G(ab|cd)w_G(ac|bd) = 0$,

$$\begin{aligned}\mathrm{Var}[X_G] &= \mathbb{E}[X_G^2] - \mathbb{E}^2[X_G] = \mathbb{E}\left[w_G^2(ab|cd) + w_G^2(ac|bd)\right] - \mathbb{E}^2[X_G] \\ &= \mathbb{E}\left[(\delta_1+\delta_2+2\delta_3) w_G^2(ab|cd)\right] - \mathbb{E}^2[X_G].\end{aligned} \tag{14}$$

- $\delta_1$: Here,

$$\begin{aligned}\mathbb{E}\left[\delta_1 w_G(ab|cd)\right] &= \int_0^{x_0}\int_0^{+\infty} e^{-\lambda(2x+2y+L)} e^{-x} e^{-y}\, dy\, dx \\ &= \frac{e^{-\lambda L}(1-e^{-(1+2\lambda)x_0})}{(1+2\lambda)^2} = \frac{e^{-\lambda L} x_0}{1+2\lambda} + O(x_0^2) = \frac{e^{-\lambda L} x_0}{1+2\lambda} + O(f^2);\end{aligned}$$

  and

$$\mathbb{E}\left[\delta_1 w_G^2(ab|cd)\right] \le \mathbb{E}\left[\delta_1 w_G(ab|cd)\right] = O(f).$$

- $\delta_2$: Here,

$$\mathbb{E}[\delta_2 w_G(ab|cd)] = \int_{x_0}^{+\infty}\int_0^{y_0} e^{-\lambda(2x+2y+L)} e^{-x} e^{-y} dy dx$$
$$= \frac{e^{-\lambda L}(1-e^{-(1+2\lambda)y_0})e^{-(1+2\lambda)x_0}}{(1+2\lambda)^2} = \frac{e^{-\lambda L} y_0}{1+2\lambda} + O(f^2);$$

and

$$\mathbb{E}[\delta_2 w_G^2(ab|cd)] \le \mathbb{E}[\delta_2 w_G(ab|cd)] = O(f).$$

- $\delta_3$: Similar to the unbalanced case, when $G$ has the topology $ab|cd$, either $p'$ or $q'$ must be the lowest point of coalescence, and by symmetry, the two cases must have the same PDFs. Thus,

$$\mathbb{E}[\delta_3 w_G^2(ab|cd)] = \int_{x_0}^{+\infty}\int_{x-x_0+y_0}^{+\infty} e^{-\lambda(4x+4y+2L)} 2e^{-x_0} e^{-y_0} e^{-6x+6x_0} e^{-y+x-x_0+y_0} dy dx$$
$$= \int_{x_0}^{+\infty} e^{-4\lambda(x+x-x_0+y_0)-2\lambda L} 2e^{-x_0} e^{-y_0} e^{-6x+6x_0} \frac{1}{1+4\lambda} dx$$
$$= \frac{1}{(3+4\lambda)(1+4\lambda)} e^{-(1+4\lambda)(x_0+y_0)-2\lambda L} = \frac{1}{(3+4\lambda)(1+4\lambda)} e^{-(1+4\lambda)f-2\lambda L}.$$

Replacing in (13), we get

$$\mathbb{E}[X_G] = \mathbb{E}[(\delta_1+\delta_2) w_G(ab|cd)] = \frac{e^{-\lambda L}(x_0+y_0)}{1+2\lambda} + O(f^2) = \frac{e^{-\lambda L} f}{1+2\lambda} + O(f^2);$$

and replacing in (14), we get

$$\mathrm{Var}[X_G] = \mathbb{E}[(\delta_1+\delta_2+2\delta_3) w_G^2(ab|cd)] - \mathbb{E}^2[X_G]$$
$$= \mathbb{E}[2\delta_3 w_G^2(ab|cd)] + O(f)$$
$$= \frac{2e^{-(1+4\lambda)f-2\lambda L}}{(3+4\lambda)(1+4\lambda)} + O(f) = \frac{2e^{-2\lambda L}}{(3+4\lambda)(1+4\lambda)} + O(f),$$

from which our assumption of $\mathrm{Var}[X_G] = \Theta_f(1)$ follows.

Thus, in both balanced and unbalanced cases,

$$\frac{\mathbb{E}[X_G]}{\sqrt{\mathrm{Var}[X_G]}} = \frac{\frac{e^{-\lambda L}}{1+2\lambda} f + O(f^2)}{\sqrt{\frac{2e^{-2\lambda L}}{(1+4\lambda)(3+4\lambda)}} + O(f)} = \sqrt{1+\frac{4\lambda+4\lambda^2}{3(1+2\lambda)^2}}\sqrt{\frac{3}{2}} f + O(f^2)$$

□

---

PROPOSITION 3. *For a true quartet species tree $S^*$ with topology $ab|cd$ and input gene trees $\mathcal{G}$ generated under the variable rate model, let $f$ be the distance between anchors of $S^*$ and $L$ be the total length of all other branches. Assume that for every branch segment $I$, the variance of its multiplier is bounded above: $Var(\Lambda^I_{S^\dagger}) \le \varepsilon^2$ where $\varepsilon^2 = \frac{e^{-\lambda L}}{(16+32\lambda)+(6+32\lambda+32\lambda^2)L}\left(\frac{20(\lambda+\lambda^2)}{9(1+2\lambda)^2}\right)^3$. As $f \to 0$, given $k = \Theta(f^{-2})$ gene trees, we have $Var[X_G] = \Theta_f(1)$ and*

$$\frac{\mathbb{E}[X_G]}{\sqrt{Var[X_G]}} \ge \sqrt{\frac{3}{2}}\left(1-\frac{4\lambda^2}{(1+4\lambda)^2}\right)^{-\frac{1}{2}} f + O(f^2).$$

*Proof.* We follow the same logic in proof of Proposition 2.

Case 1: Unbalanced trees. Let $P(x)$ be functions to random variables denoting SU difference in heights of points $(p,p')$ where $p'$ is $x$ CU distance above $p$; let $R(z)$ be functions to random variables denoting SU difference in heights of points $(r,r')$ where $r'$ is $z$ CU distance above $r$. Note that $P(f+y_0)+R(z)=P(f+y_0+z)$ where $P(f+y_0)$ denote the SU length of $(p,r)$. Let random variable $\Lambda:=\left(l_{S^\dagger}(a,p)+l_{S^\dagger}(b,p)+l_{S^\dagger}(c,r)+l_{S^\dagger}(d,r)\right)$ be the total SU terminal branch lengths and the constant value $L$ be the CU distance corresponding to $\Lambda$.

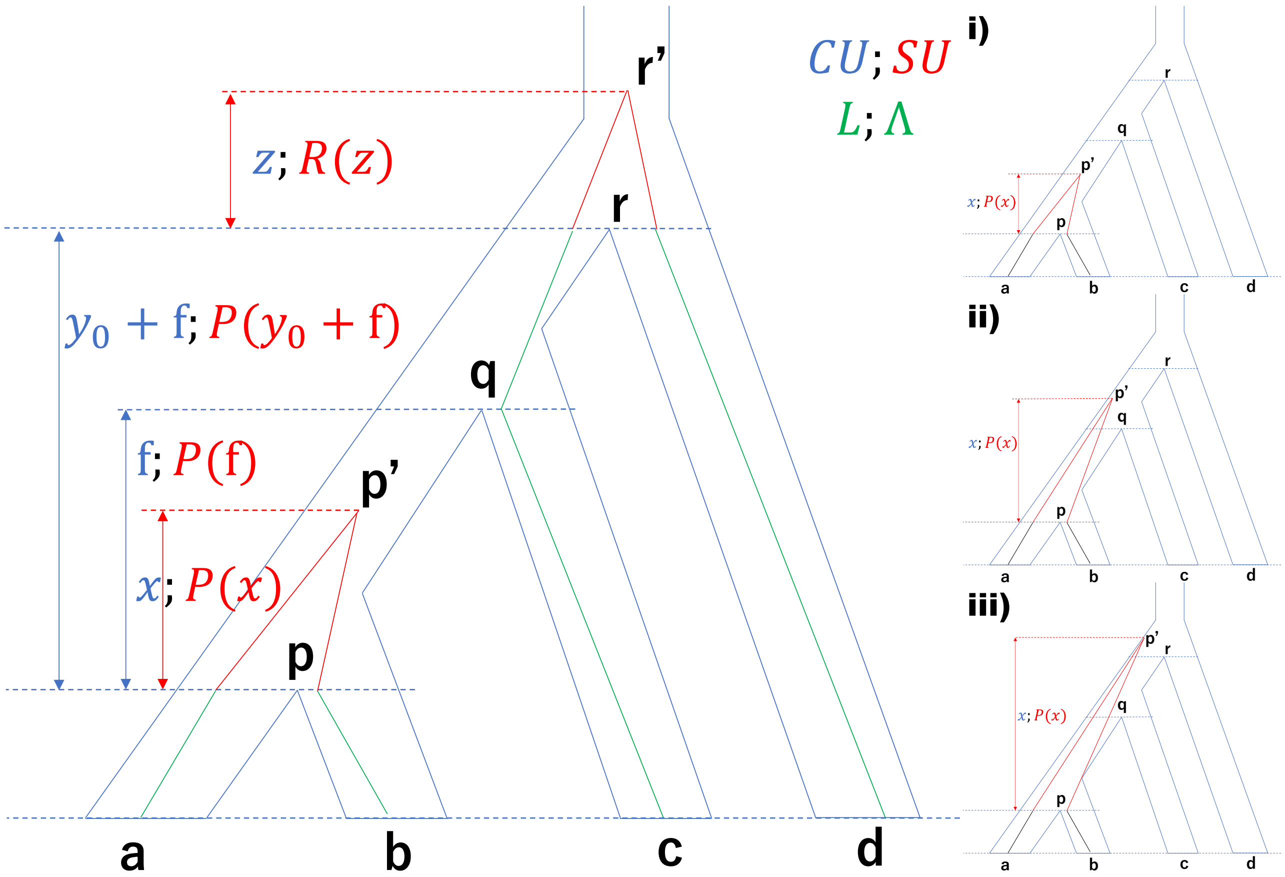


**FIG. S24.** Illustration of the unbalanced case. Lengths in CU/SU units are denoted in blue/red. Branches in green have a total length $L/\Lambda$ in CU/SU units. The right-hand side shows the position of $p'$ in relation to $q$ and $r$ in various cases.

- $\delta_1$: When $G$ has topology $ab|cd$, $p'$ must be the lowest point of coalescence. Thus,

$$
\begin{aligned}
&\mathbb{E}\left[\delta_1 w_G(ab|cd)\right]\\
=&\mathbb{E}\left[\int_0^f\int_0^{+\infty} e^{-2P(x)-2R(z)-\Lambda} f_X(x) f_{Z|X}(z;x)\,dz\,dx\right]\\
=&\mathbb{E}\left[\int_0^f\int_0^{+\infty} e^{-2P(x)-2R(z)-\Lambda} e^{-x}e^{-z}\,dz\,dx\right]\\
=&\mathbb{E}\left[\int_0^f\int_0^{+\infty} e^{-2P(x)-2R(z)-\Lambda-x-z}\,dz\,dx\right];
\end{aligned}
$$

and

$$\mathbb{E}\left[\delta_1 w_G^2(ab|cd)\right]\leq \mathbb{E}\left[\delta_1 w_G(ab|cd)\right]=O(f).$$

- $\delta_2$: When $G$ has topology $ab|cd$, $p'$ must be the lowest point of coalescence. Thus,

$$\begin{aligned}&\mathbb{E}\big[\delta_2 w_G^2(ab|cd)\big]\\ =&\mathbb{E}\Big[\int_f^{f+y_0}\int_0^{+\infty} e^{-4P(x)-4R(z)-2\Lambda} f_X(x) f_{Z|Y}(z;y)\,dzdx\Big]\\ =&\mathbb{E}\Big[\int_f^{f+y_0}\int_0^{+\infty} e^{-4P(x)-4R(z)-2\Lambda} e^{-3x+2f} e^{-z}\,dzdx\Big]\\ =&\int_f^{f+y_0}\int_0^{+\infty}\mathbb{E}\big[e^{-4P(x)-4R(z)-2\Lambda}\big] e^{-3x-z+2f}\,dzdx.\end{aligned}$$

- $\delta_3$: When $G$ has the topology $ab|cd$, either $p'$ or $q'$ must be the lowest point of coalescence, and by symmetry, the two cases must have the same PDFs. Thus,

$$\begin{aligned}&\mathbb{E}\big[\delta_3 w_G^2(ab|cd)\big]\\ =&\mathbb{E}\Big[\int_{f+y_0}^{+\infty}\int_{x-f-y_0}^{+\infty} e^{-4P(x)-4R(z)-2\Lambda} 2 f_X(x) f_{Z|X}(z;x)\,dzdx\Big]\\ =&\mathbb{E}\Big[\int_{f+y_0}^{+\infty}\int_{x-f-y_0}^{+\infty} e^{-4P(x)-4R(z)-2\Lambda} 2e^{-6x+5f+3y_0} e^{-z+x-f-y_0}\,dzdx\Big]\\ =&\int_{f+y_0}^{+\infty}\int_{x-f-y_0}^{+\infty}\mathbb{E}\big[e^{-4P(x)-4R(z)-2\Lambda}\big] 2e^{-5x-z+4f+2y_0}\,dzdx.\end{aligned}$$

Replacing in (11), by Jensen's inequality, we get

$$\begin{aligned}\mathbb{E}[X_G]=\mathbb{E}\big[\delta_1 w_G(ab|cd)\big]=&\mathbb{E}\Big[\int_0^f\int_0^{+\infty} e^{-2P(x)-2R(z)-\Lambda-x-z}\,dzdx\Big]\\ \geq&\int_0^f\int_0^{+\infty} e^{\mathbb{E}[-2P(x)-2R(z)-\Lambda-x-z]}\,dzdx\\ =&\int_0^f\int_0^{+\infty} e^{-2\lambda x-2\lambda z-\lambda L-x-z}\,dzdx\\ =&\frac{e^{-\lambda L}(1-e^{-(1+2\lambda)f})}{(1+2\lambda)^2}=\frac{e^{-\lambda L}}{1+2\lambda}f+O(f^2).\end{aligned}$$

And replacing in (12), we get

$$\begin{aligned}\mathrm{Var}[X_G]=&\mathbb{E}\big[(\delta_1+2\delta_2+2\delta_3)w_G^2(ab|cd)\big]-\mathbb{E}^2[X_G]=\mathbb{E}\big[2(\delta_2+\delta_3)w_G^2(ab|cd)\big]+O(f)\\ =&\int_f^{f+y_0}\int_0^{+\infty}\mathbb{E}\big[e^{-4P(x)-4R(z)-2\Lambda}\big]2e^{-3x-z+2f}\,dzdx\\ &+\int_{f+y_0}^{+\infty}\int_{x-f-y_0}^{+\infty}\mathbb{E}\big[e^{-4P(x)-4R(z)-2\Lambda}\big]4e^{-5x-z+2f+2y_0}\,dzdx+O(f),\end{aligned}$$

from which our assumption of $\mathrm{Var}[X_{G^*}]=\Theta_f(1)$ follows.

Let $F_P(u;x)$, $F_R(v;z)$, and $F_\Lambda(w)$ be the CDF of $P(x)$, $R(z)$, and $\Lambda$ respectively; let $F_{PR\Lambda}(u,v,w;x,z)$ and $F_{PR\Lambda}(u,v,w;x,z)$ be the joint CDF and the joint PDF. Let $F_P^{-1}(t;x)$, $F_R^{-1}(t;z)$, and $F_\Lambda^{-1}(t)$ be the inverse function of CDF of $P(x)$, $R(z)$, and $\Lambda$ respectively.

Then,

$$\begin{aligned}
&\mathbb{E}\Big[e^{-2\big(2P(x)+2R(z)+\Lambda\big)}\Big]\\
&=\int_0^{+\infty}\int_0^{+\infty}\int_0^{+\infty}e^{-2(2u+2v+w)}F_{PR\Lambda}(u,v,w;x,z)dwdvdu\\
&=\int_0^{+\infty}\int_0^{+\infty}\int_0^{+\infty}e^{-2(2u+2v+w)}\frac{\partial^3F_{PR\Lambda}}{\partial u\partial v\partial w}dwdvdu\\
&=\int_0^{+\infty}\int_0^{+\infty}\Big(e^{-2(2u+2v+w)}\frac{\partial^2F_{PR\Lambda}}{\partial u\partial v}\Big|_{w=0}^{+\infty}\\
&\quad-\int_0^{+\infty}(-2)e^{-2(2u+2v+w)}\frac{\partial^2F_{PR\Lambda}}{\partial u\partial v}dw\Big)dvdu\\
&=\int_0^{+\infty}\int_0^{+\infty}\int_0^{+\infty}2e^{-2(2u+2v+w)}\frac{\partial^2F_{PR\Lambda}}{\partial u\partial v}dvdudw\\
&=\int_0^{+\infty}\int_0^{+\infty}\Big(2e^{-2(2u+2v+w)}\frac{\partial F_{PR\Lambda}}{\partial u}\Big|_{v=0}^{+\infty}-\int_0^{+\infty}(-8)e^{-2(2u+2v+w)}\frac{\partial F_{PR\Lambda}}{\partial u}dv\Big)dudw\\
&=\int_0^{+\infty}\int_0^{+\infty}\int_0^{+\infty}8e^{-2(2u+2v+w)}\frac{\partial F_{PR\Lambda}}{\partial u}dudwdv\\
&=\int_0^{+\infty}\int_0^{+\infty}\Big(8e^{-2(2u+2v+w)}F_{PR\Lambda}(u,v,w;x,z)\Big|_{u=0}^{+\infty}\\
&\quad-\int_0^{+\infty}(-32)e^{-2(2u+2v+w)}F_{PR\Lambda}(u,v,w;x,z)du\Big)dwdv\\
&=\int_0^{+\infty}\int_0^{+\infty}\int_0^{+\infty}32e^{-2(2u+2v+w)}F_{PR\Lambda}(u,v,w;x,z)dwdvdu\\
&\le\int_0^{+\infty}\int_0^{+\infty}\int_0^{+\infty}32e^{-2(2u+2v+w)}\min\{F_P(u;x),F_R(v;z),F_\Lambda(w)\}dwdvdu\\
&=\int_0^{+\infty}\int_0^{+\infty}\int_0^{+\infty}32e^{-2(2u+2v+w)}\Big(\int_0^1 1_{t\le F_P(u;x)}1_{t\le F_R(v;z)}1_{t\le F_\Lambda(w)}dt\Big)dwdvdu\\
&=\int_0^1\int_0^{+\infty}\int_0^{+\infty}\int_0^{+\infty}32e^{-2(2u+2v+w)}1_{u\ge F_P^{-1}(t;x)}1_{v\ge F_R^{-1}(t;z)}1_{w\ge F_\Lambda^{-1}(t)}dwdvdudt\\
&=\int_0^1\int_{F_P^{-1}(t;x)}^{+\infty}\int_{F_R^{-1}(t;z)}^{+\infty}\int_{F_\Lambda^{-1}(t)}^{+\infty}32e^{-2(2u+2v+w)}dwdvdudt\\
&=\int_0^1 e^{-2\big(2F_P^{-1}(t;x)+2F_R^{-1}(t;z)+F_\Lambda^{-1}(t)\big)}dt.
\end{aligned}$$

Thus, for any $0<t_0<1$,

$$\begin{aligned}
&\mathbb{E}\Big[e^{-2\big(2P(x)+2R(z)+\Lambda\big)}\Big]\\
&\le\int_0^1 e^{-2\big(2F_P^{-1}(t;x)+2F_R^{-1}(t;z)+F_\Lambda^{-1}(t)\big)}dt\\
&\le\int_0^{t_0}\overbrace{e^{-2\big(2F_P^{-1}(0;x)+2F_R^{-1}(0;z)+F_\Lambda^{-1}(0)\big)}}^{1}dt+\int_{t_0}^1 e^{-2\big(2F_P^{-1}(t_0;x)+2F_R^{-1}(t_0;z)+F_\Lambda^{-1}(t_0)\big)}dt\\
&\le t_0+e^{-2\big(2F_P^{-1}(t_0;x)+2F_R^{-1}(t_0;z)+F_\Lambda^{-1}(t_0)\big)}.
\end{aligned}$$

By Chebyshev's inequality (using $t_0^{-\frac{1}{2}}$ as the constant), $F_P^{-1}(t_0;x)\ge(\lambda-\frac{\varepsilon}{\sqrt{t_0}})x$, $F_R^{-1}(t_0;z)\ge(\lambda-\frac{\varepsilon}{\sqrt{t_0}})y$, and $F_\Lambda^{-1}(t_0)\ge(\lambda-\frac{\varepsilon}{\sqrt{t_0}})L$. Thus,

$$\mathbb{E}\Big[e^{-2\big(2P(x)+2R(z)+\Lambda\big)}\Big]\le t_0+e^{(-\lambda+\frac{\varepsilon}{\sqrt{t_0}})(4x+4z+2L)}.$$

Thus,

$$
\begin{aligned}
\mathrm{Var}[X_{G^*}] \le & \int_{f}^{f+y_0}\int_0^{+\infty}\Big(t_0+e^{(-\lambda+\frac{\varepsilon}{\sqrt{t_0}})(4x+4z+2L)}\Big)2e^{-3x-z+2f}dzdx \\
&+\int_{f+y_0}^{+\infty}\int_{x-f-y_0}^{+\infty}\Big(t_0+e^{(-\lambda+\frac{\varepsilon}{\sqrt{t_0}})(4x+4z+2L)}\Big)4e^{-5x-z+2f+2y_0}dzdx+O(f) \\
=&\int_{f}^{f+y_0}\Big(2t_0e^{-3x+2f}+\frac{2}{1+4\lambda-\frac{4\varepsilon}{\sqrt{t_0}}}e^{(-\lambda+\frac{\varepsilon}{\sqrt{t_0}})(4x+2L)-3x+2f}\Big)dx \\
&+\int_{f+y_0}^{+\infty}\Big(4t_0e^{-6x+3f+3y_0}+\frac{4}{1+4\lambda-\frac{4\varepsilon}{\sqrt{t_0}}}e^{(-\lambda+\frac{\varepsilon}{\sqrt{t_0}})(8x-4f-4y_0+2L)-6x+3f+3y_0}\Big)dx+O(f) \\
=&\frac{2}{3}t_0(e^{-f}-e^{-f-3y_0})+\frac{2}{(1+4\lambda-\frac{4\varepsilon}{\sqrt{t_0}})(3+4\lambda-\frac{4\varepsilon}{\sqrt{t_0}})}\big(e^{(-\lambda+\frac{\varepsilon}{\sqrt{t_0}})(4f+2L)-f}-e^{(-\lambda+\frac{\varepsilon}{\sqrt{t_0}})(4f+4y_0+2L)-f-3y_0}\big) \\
&+\frac{4}{6}t_0e^{-3f-3y_0}+\frac{4}{(1+4\lambda-\frac{4\varepsilon}{\sqrt{t_0}})(6+8\lambda-\frac{8\varepsilon}{\sqrt{t_0}})}e^{(-\lambda+\frac{\varepsilon}{\sqrt{t_0}})(4f+4y_0+2L)-3f-3y_0}+O(f) \\
=&\frac{2}{3}t_0+\frac{2e^{-2L(\lambda-\frac{\varepsilon}{\sqrt{t_0}})}}{(1+4\lambda-\frac{4\varepsilon}{\sqrt{t_0}})(3+4\lambda-\frac{4\varepsilon}{\sqrt{t_0}})}+O(f).
\end{aligned}
$$

Case 2: Balanced tree. Let $P(x)$ be functions to random variables denoting SU difference in heights of points $(p,p')$ where $p'$ is $x$ CU distance above $p$; let $Q(y)$ be functions to random variables denoting SU difference in heights of points $(q,q')$ where $q'$ is $y$ CU distance above $q$. Note that $P(x_0+z)-P(x_0)=Q(y_0+z)-Q(y_0)$ where $P(x_0)$ and $Q(y_0)$ denote the SU length of $(p,r)$ and $(q,r)$, respectively. Let random variable $\Lambda:=\big(l_{S^\dagger}(a,p)+l_{S^\dagger}(b,p)+l_{S^\dagger}(c,q)+l_{S^\dagger}(d,q)\big)$ be the total SU terminal branch lengths and the constant value $L$ be the CU distance corresponding to $\Lambda$.

- $\delta_1$: Here,

  $$\mathbb{E}\big[\delta_1 w_G(ab|cd)\big]=\mathbb{E}\Big[\int_0^{x_0}\int_0^{+\infty}e^{-2P(x)-2Q(y)-\Lambda}e^{-x}e^{-y}dydx\Big];$$

  and

  $$\mathbb{E}\big[\delta_1 w_G^2(ab|cd)\big]\le\mathbb{E}\big[\delta_1 w_G(ab|cd)\big]=O(f).$$

- $\delta_2$: Here,

  $$\mathbb{E}\big[\delta_2 w_G(ab|cd)\big]=\mathbb{E}\Big[\int_{x_0}^{+\infty}\int_0^{y_0}e^{-2P(x)-2Q(y)-\Lambda}e^{-x}e^{-y}dydx\Big];$$

  and

  $$\mathbb{E}\big[\delta_2 w_G^2(ab|cd)\big]\le\mathbb{E}\big[\delta_2 w_G(ab|cd)\big]=O(f).$$

- $\delta_3$: Similar to the unbalanced case, when $G$ has the topology $ab|cd$, either $p'$ or $q'$ must be the lowest point of coalescence, and by symmetry, the two cases must have the same PDFs. Thus,

  $$
  \begin{aligned}
  \mathbb{E}\big[\delta_3 w_G^2(ab|cd)\big]=&\mathbb{E}\Big[\int_{x_0}^{+\infty}\int_{x-x_0+y_0}^{+\infty}e^{-4P(x)-4Q(y)-2\Lambda}2e^{-x_0}e^{-y_0}e^{-6x+6x_0}e^{-y+x-x_0+y_0}dydx\Big] \\
  =&\int_{x_0}^{+\infty}\int_{x-x_0+y_0}^{+\infty}\mathbb{E}\Big[e^{-4P(x)-4Q(y)-2\Lambda}\Big]2e^{-5x-y+4x_0}dydx.
  \end{aligned}
  $$

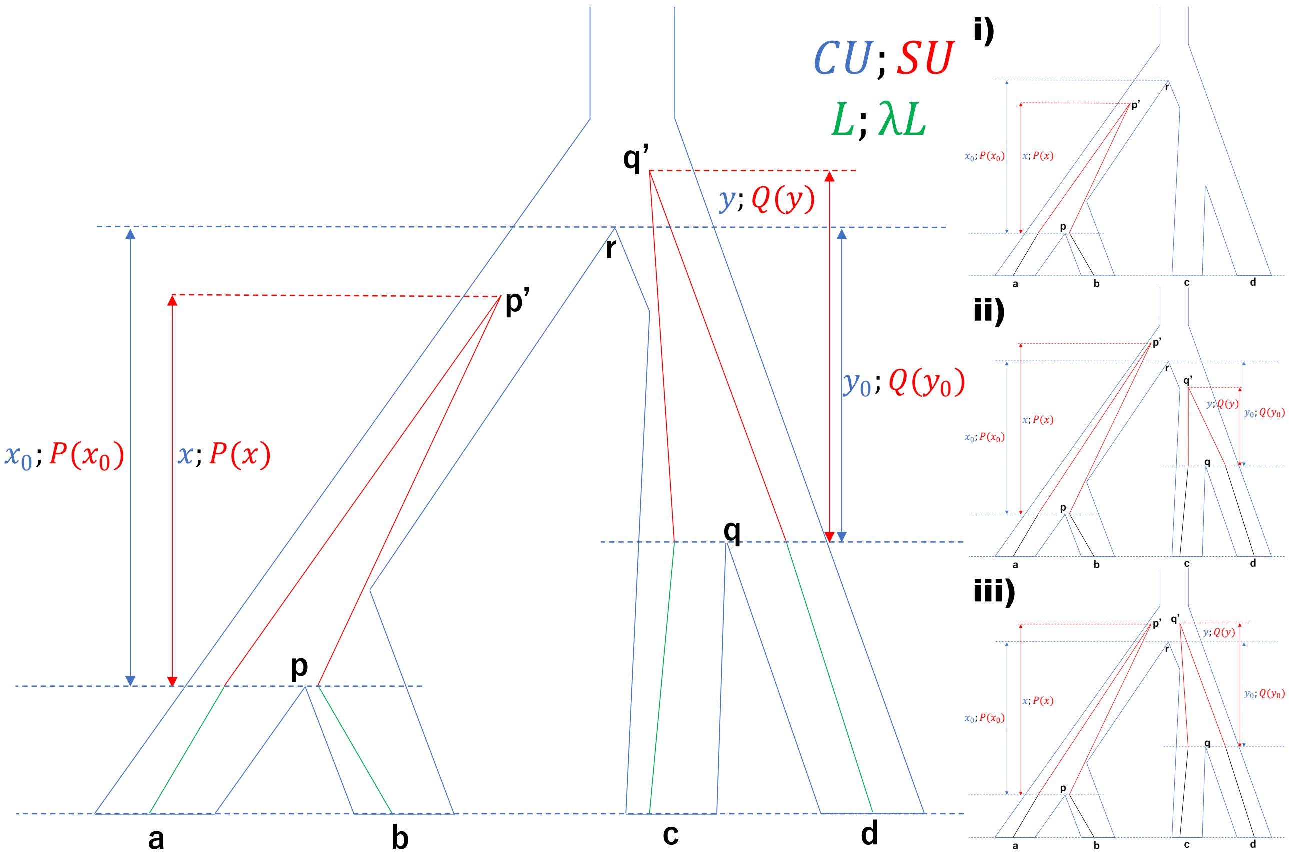


**FIG. S25.** Illustration of the unbalanced case. Lengths in CU/SU units are denoted in blue/red. Branches in green have a total length $L/\Lambda$ in CU/SU units. The right-hand side shows the position of $p'$ and $q'$ in relation to $r$ in various cases.

Replacing in (13), we get

$$\begin{aligned}\mathbb{E}[X_G] &= \mathbb{E}\left[(\delta_1+\delta_2)w_G(ab|cd)\right]\\ &= \mathbb{E}\Big[\int_0^{x_0}\int_0^{+\infty} e^{-2P(x)-2Q(y)-\Lambda}e^{-x}e^{-y}dydx + \int_{x_0}^{+\infty}\int_0^{y_0} e^{-2P(x)-2Q(y)-\Lambda}e^{-x}e^{-y}dydx\Big]\\ &\geq \int_0^{x_0}\int_0^{+\infty} e^{-2\lambda x-2\lambda y-\lambda L}e^{-x}e^{-y}dydx + \int_{x_0}^{+\infty}\int_0^{y_0} e^{-2\lambda x-2\lambda y-\lambda L}e^{-x}e^{-y}dydx\\ &= \frac{(x_0+y_0)e^{-\lambda L}}{1+2\lambda} + O(f^2) = \frac{fe^{-\lambda L}}{1+2\lambda} + O(f^2);\end{aligned}$$

and replacing in (14), for any $0<t_0<1$,

$$\begin{aligned}
\mathrm{Var}[X_G] &= \mathbb{E}\left[(\delta_1+\delta_2+2\delta_3)w_G^2(ab|cd)\right]-\mathbb{E}^2[X_G]\\
&= \mathbb{E}\left[2\delta_3 w_G^2(ab|cd)\right]+O(f)\\
&= \int_{x_0}^{+\infty}\int_{x-x_0+y_0}^{+\infty}\mathbb{E}\left[e^{-4P(x)-4Q(y)-2\Lambda}\right]4e^{-5x-y+4x_0}dydx\\
&\le \int_{x_0}^{+\infty}\int_{x-x_0+y_0}^{+\infty}\left(t_0+e^{(-\lambda+\frac{\varepsilon}{\sqrt{t_0}})(4x+4y+2L)}\right)4e^{-5x-y+4x_0}dydx+O(f)\\
&= \int_{x_0}^{+\infty}\left(4e^{-6x-y_0+5x_0}t_0+\frac{4}{1+4\lambda-\frac{4\varepsilon}{\sqrt{t_0}}}e^{-6x-y_0+5x_0+(-\lambda+\frac{\varepsilon}{\sqrt{t_0}})(8x-4x_0+4y_0+2L)}\right)dx+O(f)\\
&= \frac{4}{6}e^{-x_0-y_0}t_0+\frac{4}{(1+4\lambda-\frac{4\varepsilon}{\sqrt{t_0}})(6+8\lambda-\frac{8\varepsilon}{\sqrt{t_0}})}e^{-x_0-y_0+(-\lambda+\frac{\varepsilon}{\sqrt{t_0}})(4x_0+4y_0+2L)}+O(f)\\
&= \frac{2}{3}t_0+\frac{2e^{-2L(\lambda-\frac{\varepsilon}{\sqrt{t_0}})}}{(1+4\lambda-\frac{4\varepsilon}{\sqrt{t_0}})(3+4\lambda-\frac{4\varepsilon}{\sqrt{t_0}})}+O(f),
\end{aligned}$$

from which our assumption of $\mathrm{Var}[X_G]=\Theta_f(1)$ follows.

Thus, for both balanced and unbalanced trees, the variance is bounded the by same expression, and thus in both cases,

$$\begin{aligned}
\mathrm{Var}[X_{G^*}] &\le \frac{2}{3}t_0+2\frac{\frac{e^{-2\lambda L}}{(1+4\lambda)(3+4\lambda)}}{(1-\frac{4\varepsilon}{(1+4\lambda)\sqrt{t_0}})(1-\frac{4\varepsilon}{(3+4\lambda)\sqrt{t_0}})e^{-\frac{2\varepsilon L}{\sqrt{t_0}}}}+O(f)\\
&\le \frac{2}{3}t_0+2\frac{\frac{e^{-2\lambda L}}{(1+4\lambda)(3+4\lambda)}}{(1-\frac{4\varepsilon}{(1+4\lambda)\sqrt{t_0}})(1-\frac{4\varepsilon}{(3+4\lambda)\sqrt{t_0}})(1-\frac{2\varepsilon L}{\sqrt{t_0}})}+O(f)\\
&\le \frac{2}{3}t_0+2\frac{\frac{e^{-2\lambda L}}{(1+4\lambda)(3+4\lambda)}}{(1-\frac{4\varepsilon}{(1+4\lambda)\sqrt{t_0}}-\frac{4\varepsilon}{(3+4\lambda)\sqrt{t_0}}-\frac{2\varepsilon L}{\sqrt{t_0}})}+O(f)\\
&= \frac{2}{3}t_0+\frac{2e^{-2\lambda L}}{(3+16\lambda+16\lambda^2)-\frac{\varepsilon}{\sqrt{t_0}}((16+32\lambda)+(6+32\lambda+32\lambda^2)L)}+O(f).
\end{aligned}$$

Now, let $C:=(16+32\lambda)+(6+32\lambda+32\lambda^2)L, t_0=\left(\frac{C^{\frac{1}{3}}\varepsilon^{\frac{1}{3}}}{(3+16\lambda+16\lambda^2)e^{\frac{2}{3}\lambda L}}\right)^2$, we get

$$\begin{aligned}
\mathrm{Var}[X_{G^*}] &\le \frac{2e^{-2\lambda L}}{3(3+16\lambda+16\lambda^2)^2}\left((\varepsilon e^{\lambda L}C)^{\frac{2}{3}}+\frac{9+48\lambda+48\lambda^2}{1-(\varepsilon e^{\lambda L}C)^{\frac{2}{3}}}\right)+O(f)\\
&= \frac{2e^{-2\lambda L}}{3(3+16\lambda+16\lambda^2)}\left(\frac{(\varepsilon e^{\lambda L}C)^{\frac{2}{3}}}{3+16\lambda+16\lambda^2}+3+\frac{3(\varepsilon e^{\lambda L}C)^{\frac{2}{3}}}{1-(\varepsilon e^{\lambda L}C)^{\frac{2}{3}}}\right)+O(f).
\end{aligned}$$

Now, recalling that $\varepsilon=\frac{e^{-\lambda L}}{C}\left(\frac{20(\lambda+\lambda^2)}{9(1+2\lambda)^2}\right)^{\frac{3}{2}}$,

$$
\begin{aligned}
\mathrm{Var}[X_{G^*}] \le & \frac{2}{3(3+16\lambda+16\lambda^2)(1+2\lambda)^2} \\
& \Big(\frac{\frac{20}{9}(\lambda+\lambda^2)}{3+16\lambda+16\lambda^2}+3+\frac{3(\frac{20}{9})(\lambda+\lambda^2)}{1-\frac{\frac{20}{9}(\lambda+\lambda^2)}{1+4\lambda+4\lambda^2}}\Big)+O(f) \\
\le & \frac{2}{3(3+16\lambda+16\lambda^2)(1+2\lambda)^2}\Big(\frac{20}{27}\lambda+3+\frac{\frac{20}{3}(\lambda+\lambda^2)}{1-\frac{5}{9}}\Big)+O(f) \\
= & \frac{2}{3(3+16\lambda+16\lambda^2)(1+2\lambda)^2}\Big(\frac{20}{27}\lambda+3+15(\lambda+\lambda^2)\Big)+O(f) \\
< & \frac{2}{3(1+2\lambda)^2}\Big(\frac{3+16\lambda+15\lambda^2}{3+16\lambda+16\lambda^2}\Big)+O(f).
\end{aligned}
$$

□

---

THEOREM 2. *Under the conditions of Proposition 2 or Proposition 3,*

$$P\Big(\sum_{G\in\mathcal{G}} w_G(ab|cd) \le \sum_{G\in\mathcal{G}} w_G(ac|bd)\Big) \le P\Big(\sum_{G\in\mathcal{G}} \delta_G(ab|cd) \le \sum_{G\in\mathcal{G}} \delta_G(ac|bd)\Big).$$

*Proof*. We start with proving this theorem under the conditions of Proposition 2. Recall $X_G := w_G(ab|cd) - w_G(ac|bd)$ and $Y_G := \delta_G(ab|cd) - \delta_G(ac|bd)$, and let $\bar{X_\mathcal{G}} = \frac{1}{k}\sum_{G\in\mathcal{G}} X_G$ and $\bar{Y_\mathcal{G}} = \frac{1}{k}\sum_{G\in\mathcal{G}} Y_G$. Recall also that under Proposition 2, proved below, under conditions of Theorem 2, we have $\mathrm{Var}[X_G] = \Omega(1)$ and

$$\frac{\mathbb{E}[X_G]}{\sqrt{\mathrm{Var}[X_G]}} = -\sqrt{\frac{3+16\lambda+16\lambda^2}{3+16\lambda+15\lambda^2}}\sqrt{\frac{3}{2}}f + O(f^2). \tag{15}$$

Similarly, we can compute the ratio of mean and variance for $Y$ (corresponding to unweighted ASTRAL):

$$\mathbb{E}[Y_G] := \mathbb{E}\big[\delta_G(ab|cd) - \delta_G(ac|bd)\big] = 1 - e^{-f} = f + O(f^2)$$

$$\mathrm{Var}[Y_G] := \mathrm{Var}\big[\delta_G(ab|cd) - \delta_G(ac|bd)\big] = \frac{5}{3}e^{-f} - e^{-2f} = \frac{2}{3} + O(f)$$

and thus,

$$\frac{\mathbb{E}[Y_G]}{\sqrt{\mathrm{Var}[Y_G]}} = \sqrt{\frac{3}{2}}f + O(f^2). \tag{16}$$

Given Proposition 2, we can use Berry–Esseen theorem to derive

$$\mathrm{P}\big(\bar{X_\mathcal{G}} \le 0\big) = \mathrm{P}\big(\frac{\sqrt{k}}{\sqrt{\mathrm{Var}[X_G]}}(\bar{X_\mathcal{G}} - \mathbb{E}[X_G]) \le -\frac{\sqrt{k}}{\sqrt{\mathrm{Var}[X_G]}}\mathbb{E}[X_G]\big) =$$
$$\Phi(-\sqrt{k}\frac{\mathbb{E}[X_G]}{\sqrt{\mathrm{Var}[X_G]}}) + O(\frac{1}{\sqrt{k}}),$$

where $\Phi$ denotes CDF of the standard Normal distribution. Since $k = \Theta(f^{-2})$,

$$\mathrm{P}\big(\bar{X_\mathcal{G}} \le 0\big) = \Phi(-\sqrt{k}\frac{\mathbb{E}[X_G]}{\sqrt{\mathrm{Var}[X_G]}}) + O(f) \tag{17}$$

and

$$\mathrm{P}\big(\bar{Y_\mathcal{G}} \le 0\big) = \Phi(-\sqrt{k}\frac{\mathbb{E}[Y_G]}{\sqrt{\mathrm{Var}[Y_G]}}) + O(f), \tag{18}$$

Combining equations (17) and (18) with (15) and (16), we get

$$P(\sum_{G\in\mathcal{G}} w_G(ab|cd) \le \sum_{G\in\mathcal{G}} w_G(ac|bd)) = \Phi(-\sqrt{\frac{3+16\lambda+16\lambda^2}{3+16\lambda+15\lambda^2}}\sqrt{\frac{3}{2}}f\sqrt{k}) + O(f)$$

and

$$P(\sum_{G\in\mathcal{G}} \delta_G(ab|cd) \le \sum_{G\in\mathcal{G}} \delta_G(ac|bd)) = \Phi(-\sqrt{\frac{3}{2}}f\sqrt{k}) + O(f)\,.$$

As $f \to 0$, the interval $(-\sqrt{1+\frac{4\lambda+4\lambda^2}{3(1+2\lambda)^2}}\sqrt{\frac{3}{2}}f\sqrt{k}, -\sqrt{\frac{3}{2}}f\sqrt{k})$ does not shrink because $\Theta(f\sqrt{k}) = \Theta(1)$. Thus, we have

$$\Phi(-\sqrt{\frac{3}{2}}f\sqrt{k}) - \Phi(-\sqrt{1+\frac{4\lambda+4\lambda^2}{3(1+2\lambda)^2}}\sqrt{\frac{3}{2}}f\sqrt{k}) = \Theta(1)$$

ensuring that

$$P(\sum_{G\in\mathcal{G}} w_G(ab|cd) \le \sum_{G\in\mathcal{G}} w_G(ac|bd)) \le P(\sum_{G\in\mathcal{G}} \delta_G(ab|cd) \le \sum_{G\in\mathcal{G}} \delta_G(ac|bd))\,.$$

The proof under Proposition 3 is similar. Recall that under Proposition 3, $\mathrm{Var}[X_{G^*}] = \Theta_f(1)$ and

$$\frac{\mathbb{E}[X_{G^*}]}{\sqrt{\mathrm{Var}[X_{G^*}]}} \ge \sqrt{\frac{3}{2}}\Big(1-\frac{4\lambda^2}{(1+4\lambda)^2}\Big)^{-\frac{1}{2}} f + O(f^2)\,. \tag{19}$$

Given this result, the rest of the proof is similar to the proof under the conditions of Proposition 2, culminating in

$$P(\sum_{G^*\in\mathcal{G}} w_{G^*}(ab|cd) \le \sum_{G^*\in\mathcal{G}} w_{G^*}(ac|bd)) \le \Phi\left(-\Big(1-\frac{4\lambda^2}{(1+4\lambda)^2}\Big)^{-\frac{1}{2}}\sqrt{\frac{3}{2}}f\sqrt{k}\right) + O(f)\,.$$

□

#### Placement-based Algorithm

In this section, for a node $v$ in tree $G$, we let $\mathcal{L}_v$ denote the set of leaves under $v$.

*Proof of Theorem 3*

THEOREM 3. *Let $S$ be a species tree, $i$ be a species not in $\mathcal{L}_S$, $\mathcal{S}$ be the set of possible species tree topologies by placing $i$ onto $S$, and $S'$ be the output of Algorithm S1. Then, $W(S',\mathcal{G}) = \max_{\hat{S}\in\mathcal{S}} W(\hat{S},\mathcal{G})$.*

*Proof.* We start with two propositions, proved below.

PROPOSITION 5. *After each call to* `ColorLeafSet`$(\mathcal{L}^*,X,T,\mathcal{G},W)$ *with a $T \neq \emptyset$, $W[T] = \sum_{G\in\mathcal{G}} W(T,G)$.*

PROPOSITION 6. *Before calling* `OptimalTreeDP` *in line 6 of Algorithm S1, lookup table $W$ contains all tripartitions corresponding to internal nodes of all tree topologies in $\mathcal{S}$.*

By Proposition 6, all tripartitions corresponding to internal nodes of all tree topologies in $\mathcal{S}$ pre-computed. Then, `OptimalTreeDP` uses a dynamic programming algorithm similar to the one formulated by Mirarab and Warnow 2015 to compute $\text{argmax}_{\hat{S}\in\mathcal{S}} W(\hat{S},\mathcal{G})$. □

---

PROPOSITION 5. *After each call to* `ColorLeafSet`$(\mathcal{L}^*,X,T,\mathcal{G},W)$ *with a $T \neq \emptyset$, $W[T] = \sum_{G\in\mathcal{G}} W(T,G)$.*

*Proof.* For a gene tree node $w$ and a color $X$, let $\mathcal{L}_w^X$ denote the set of leaves in $\mathcal{L}_w$ colored by $X$. For an internal node $w$, let $u,v$ be the children of $w$, $p$ be the parent of $w$ (if $w$ is not the root), and $e$ denote the branch $(w,p)$. For a leaf $i$ and internal node $w$, let $\mathcal{P}_{i,w}$ denote path between $i$ and $w$ and $s(\mathcal{P}) = 1 - \prod_{\hat{e}\in\mathcal{P}}(1-s(\hat{e}))$. For leaves $i,j$, let $m(i,j)$ denote MRCA of $i$ and $j$. Referring back to Table S1, we first establish the connection between recursive formulas of the algorithm and counter definitions.

- When $u_X = \sum_{i\in\mathcal{L}_u^X} e^{-l(\mathcal{P}_{i,w})}$, $v_X = \sum_{i\in\mathcal{L}_v^X} e^{-l(\mathcal{P}_{i,w})}$,
$$w_X := \Big((u_X + v_X)e^{-l(e)}\Big) = \sum_{i\in\mathcal{L}_w^X} e^{-l(\mathcal{P}_{i,w})} e^{-l(e)} = \sum_{i\in\mathcal{L}_w^X} e^{-l(\mathcal{P}_{i,p})}.$$
- When $u_{XX}^+ = \sum_{\{i,j\}\subseteq\mathcal{L}_u^X} e^{-l(\mathcal{P}_{i,j})}$, $v_{XX}^+ = \sum_{\{i,j\}\subseteq\mathcal{L}_v^X} e^{-l(\mathcal{P}_{i,j})}$,
$$\begin{aligned} w_{XX}^+ := u_{XX}^+ + v_{XX}^+ + u_X v_X &= \sum_{\{i,j\}\subseteq\mathcal{L}_u^X} e^{-l(\mathcal{P}_{i,j})} + \sum_{\{i,j\}\subseteq\mathcal{L}_v^X} e^{-l(\mathcal{P}_{i,j})} + \sum_{i\in\mathcal{L}_u^X} e^{-l(\mathcal{P}_{i,w})} \sum_{j\in\mathcal{L}_v^X} e^{-l(\mathcal{P}_{j,w})} \\ &= \sum_{\{i,j\}\subseteq\mathcal{L}_u^X} e^{-l(\mathcal{P}_{i,j})} + \sum_{\{i,j\}\subseteq\mathcal{L}_v^X} e^{-l(\mathcal{P}_{i,j})} + \sum_{i\in\mathcal{L}_u^X}\sum_{j\in\mathcal{L}_v^X} e^{-l(\mathcal{P}_{i,j})} = \sum_{\{i,j\}\subseteq\mathcal{L}_w^X} e^{-l(\mathcal{P}_{i,j})}. \end{aligned}$$
- For $X \neq Y$, when $u_{XY}^+ = \sum_{(i,j)\in\mathcal{L}_u^X\times\mathcal{L}_u^Y} e^{-l(\mathcal{P}_{i,j})}$, $v_{XY}^+ = \sum_{(i,j)\in\mathcal{L}_v^X\times\mathcal{L}_v^Y} e^{-l(\mathcal{P}_{i,j})}$,
$$\begin{aligned} w_{XY}^+ &:= u_{XY}^+ + v_{XY}^+ + u_X v_Y + u_Y v_X \\ &= \sum_{(i,j)\in\mathcal{L}_u^X\times\mathcal{L}_u^Y} e^{-l(\mathcal{P}_{i,j})} + \sum_{(i,j)\in\mathcal{L}_v^X\times\mathcal{L}_v^Y} e^{-l(\mathcal{P}_{i,j})} + \sum_{(i,j)\in\mathcal{L}_u^X\times\mathcal{L}_v^Y} e^{-l(\mathcal{P}_{i,j})} + \sum_{(i,j)\in\mathcal{L}_v^X\times\mathcal{L}_u^Y} e^{-l(\mathcal{P}_{i,j})} \\ &= \sum_{\{i,j\}\subseteq\mathcal{L}_w^X\times\mathcal{L}_w^X} e^{-l(\mathcal{P}_{i,j})}. \end{aligned}$$

- When $u^-_{XX} = \sum_{\{i,j\}\subseteq \mathcal{L}^X_u} e^{-l(\mathcal{P}_{i,j})} \prod_{\hat{e}\in\mathcal{P}_{m(i,j),w}} \big(1-s(\hat{e})\big)$, $v^-_{XX} = \sum_{\{i,j\}\subseteq \mathcal{L}^X_v} e^{-l(\mathcal{P}_{i,j})} \prod_{\hat{e}\in\mathcal{P}_{m(i,j),w}} \big(1-s(\hat{e})\big)$,

$$\begin{aligned} w^-_{XX} &:= (u^-_{XX} + v^-_{XX} + u_X v_X)\big(1-s(e)\big) \\ &= \sum_{\{i,j\}\subseteq \mathcal{L}^X_u} e^{-l(\mathcal{P}_{i,j})} \prod_{\hat{e}\in\mathcal{P}_{m(i,j),p}} \big(1-s(\hat{e})\big) + \sum_{\{i,j\}\subseteq \mathcal{L}^X_v} e^{-l(\mathcal{P}_{i,j})} \prod_{\hat{e}\in\mathcal{P}_{m(i,j),p}} \big(1-s(\hat{e})\big) \\ &\quad + \sum_{(i,j)\in\mathcal{L}^X_u\times\mathcal{L}^X_v} e^{-l(\mathcal{P}_{i,j})}\big(1-s(e)\big) = \sum_{\{i,j\}\subseteq \mathcal{L}^X_w} e^{-l(\mathcal{P}_{i,j})} \prod_{\hat{e}\in\mathcal{P}_{m(i,j),p}} \big(1-s(\hat{e})\big). \end{aligned}$$

- When $u^-_{XY} = \sum_{(i,j)\in\mathcal{L}^X_u\times\mathcal{L}^Y_u} e^{-l(\mathcal{P}_{i,j})}\big(1-s(\mathcal{P}_{m(i,j),w})\big)$, $v^-_{XY} = \sum_{(i,j)\in\mathcal{L}^X_u\times\mathcal{L}^Y_u} e^{-l(\mathcal{P}_{i,j})}\big(1-s(\mathcal{P}_{m(i,j),w})\big)$, and $X\neq Y$, similarly,

$$w^-_{XY} := (u^-_{XY} + v^-_{XY} + u_X v_Y + u_Y v_X)\big(1-s(e)\big) = \sum_{(i,j)\in\mathcal{L}^X_w\times\mathcal{L}^Y_w} e^{-l(\mathcal{P}_{i,j})}\big(1-s(\mathcal{P}_{m(i,j),p})\big).$$

- For $X\neq Y$, when $u_{XX|Y} = \sum_{\{i,j\}\subseteq\mathcal{L}^X_u} \sum_{k\in\{k'\in\mathcal{L}^Y_v : \mathcal{L}_{m(i,j)} \subsetneqq \mathcal{L}_{m(i,k')}\}} e^{-l(\mathcal{P}_{i,j})-l(\mathcal{P}_{k,w})} s(\mathcal{P}_{m(i,j),m(i,k)})$, $v_{XX|Y} = \sum_{\{i,j\}\subseteq\mathcal{L}^X_v} \sum_{k\in\{k'\in\mathcal{L}^Y_v : \mathcal{L}_{m(i,j)} \subsetneqq \mathcal{L}_{m(i,k')}\}} e^{-l(\mathcal{P}_{i,j})-l(\mathcal{P}_{k,w})} s(\mathcal{P}_{m(i,j),m(i,k)})$,

$$w_{XX|Y} := \big(u_{XX|Y} + v_{XX|Y} + (u^+_{XX} - u^-_{XX})v_Y + u_Y(v^+_{XX} - v^-_{XX})\big)e^{-l(e)}.$$

Notice that $(u^+_{XX} - u^-_{XX})v_Y = \sum_{\{i,j\}\subseteq\mathcal{L}^X_u} \sum_{k\in\mathcal{L}^Y_v} e^{-l(\mathcal{P}_{i,j})-l(\mathcal{P}_{k,w})} s(\mathcal{P}_{m(i,j),w})$ and $u_Y(v^+_{XX} - v^-_{XX}) = \sum_{\{i,j\}\subseteq\mathcal{L}^X_v} \sum_{k\in\mathcal{L}^Y_u} e^{-l(\mathcal{P}_{i,j})-l(\mathcal{P}_{k,w})} s(\mathcal{P}_{m(i,j),w})$. Thus,

$$\begin{aligned} w_{XX|Y} &= \sum_{\{i,j\}\subseteq\mathcal{L}^X_w} \sum_{k\in\{k'\in\mathcal{L}^Y_w : \mathcal{L}_{m(i,j)} \subsetneqq \mathcal{L}_{m(i,k')}\}} e^{-l(\mathcal{P}_{i,j})-l(\mathcal{P}_{k,w})} s(\mathcal{P}_{m(i,j),m(i,k)}) e^{-l(e)} \\ &= \sum_{\{i,j\}\subseteq\mathcal{L}^X_w} \sum_{k\in\{k'\in\mathcal{L}^Y_w : \mathcal{L}_{m(i,j)} \subsetneqq \mathcal{L}_{m(i,k')}\}} e^{-l(\mathcal{P}_{i,j})-l(\mathcal{P}_{k,p})} s(\mathcal{P}_{m(i,j),m(i,k)}). \end{aligned}$$

- Similarly, when $u_{XY|Z} = \sum_{(i,j)\in\mathcal{L}^X_u\times\mathcal{L}^Y_u} \sum_{k\in\{k'\in\mathcal{L}^Z_u : \mathcal{L}_{m(i,j)} \subsetneqq \mathcal{L}_{m(i,k')}\}} e^{-l(\mathcal{P}_{i,j})-l(\mathcal{P}_{k,w})} s(\mathcal{P}_{m(i,j),m(i,k)})$, $v^-_{XY|Z} = \sum_{(i,j)\in\mathcal{L}^X_v\times\mathcal{L}^Y_v} \sum_{k\in\{k'\in\mathcal{L}^Z_v : \mathcal{L}_{m(i,j)} \subsetneqq \mathcal{L}_{m(i,k')}\}} e^{-l(\mathcal{P}_{i,j})-l(\mathcal{P}_{k,w})} s(\mathcal{P}_{m(i,j),m(i,k)})$, for distinct $X,Y,Z$,

$$w_{XY|Z} = \sum_{(i,j)\in\mathcal{L}^X_w\times\mathcal{L}^Y_w} \sum_{k\in\{k'\in\mathcal{L}^Z_w : \mathcal{L}_{m(i,j)} \subsetneqq \mathcal{L}_{m(i,k')}\}} e^{-l(\mathcal{P}_{i,j})-l(\mathcal{P}_{k,p})} s(\mathcal{P}_{m(i,j),m(i,k)}).$$

- For distinct $X,Y,Z$,

$$\begin{aligned} w_{XX|YZ} &:= v_X u_{YZ|X} + u_X v_{YZ|X} + u_{XX|Z} v_Y + v_{XX|Z} u_Y + u_{XX|Y} v_Z + v_{XX|Y} u_Z \\ &\quad + (u^+_{YZ} v^+_{XX} - u^-_{YZ} v^-_{XX}) + (u^+_{XX} v^+_{YZ} - u^-_{XX} v^-_{YZ}). \end{aligned}$$

Notice that,

$$\begin{aligned} v_X u_{YZ|X} &= \sum_{(h,i,j,k)\in\mathcal{L}^X_v\times\mathcal{L}^Y_u\times\mathcal{L}^Z_u\times\mathcal{L}^X_u} \delta_G(hk|ij) e^{-l(\mathcal{P}_{h,w})} e^{-l(\mathcal{P}_{i,j})-l(\mathcal{P}_{k,w})} s(\mathcal{P}_{m(i,j),m(i,k)}) \\ &= \sum_{(h,i,j,k)\in\mathcal{L}^X_v\times\mathcal{L}^Y_u\times\mathcal{L}^Z_u\times\mathcal{L}^X_u} \delta_G(hk|ij) e^{-l(\mathcal{P}_{i,j})-l(\mathcal{P}_{k,h})} s(\mathcal{P}_{m(i,j),m(i,k)}) \\ &= \sum_{(h,i,j,k)\in\mathcal{L}^X_v\times\mathcal{L}^Y_u\times\mathcal{L}^Z_u\times\mathcal{L}^X_u} w_G(hk|ij). \end{aligned}$$

Similarly,

$$u_X v_{YZ|X} = \sum_{\substack{h\in\mathcal{L}_u^X\\ i\in\mathcal{L}_v^Y\\ j\in\mathcal{L}_v^Z\\ k\in\mathcal{L}_v^X}} w_G(hk|ij), u_{XX|Z} v_Y = \sum_{\substack{\{h,i\}\subseteq\mathcal{L}_u^X\\ j\in\mathcal{L}_u^Z\\ k\in\mathcal{L}_v^Y}} w_G(hi|jk), v_{XX|Z} u_Y = \sum_{\substack{\{h,i\}\subseteq\mathcal{L}_v^X\\ j\in\mathcal{L}_v^Z\\ k\in\mathcal{L}_u^Y}} w_G(hi|jk),$$

$$u_{XX|Y} v_Z = \sum_{\substack{\{h,i\}\subseteq\mathcal{L}_u^X\\ j\in\mathcal{L}_u^Y\\ k\in\mathcal{L}_v^Z}} w_G(hi|jk), v_{XX|Y} u_Z = \sum_{\substack{\{h,i\}\subseteq\mathcal{L}_v^X\\ j\in\mathcal{L}_v^Y\\ k\in\mathcal{L}_u^Z}} w_G(hi|jk).$$

Also,

$$\begin{aligned} u_{YZ}^+ v_{XX}^+ - u_{YZ}^- v_{XX}^- &= \sum_{(h,i)\in\mathcal{L}_u^Y\times\mathcal{L}_u^Z} \sum_{\{j,k\}\subseteq\mathcal{L}_v^X} e^{-l(\mathcal{P}_{h,i})-l(\mathcal{P}_{j,k})} \\ &\quad - \sum_{(h,i)\in\mathcal{L}_u^Y\times\mathcal{L}_u^Z} \sum_{\{j,k\}\subseteq\mathcal{L}_v^X} e^{-l(\mathcal{P}_{h,i})-l(\mathcal{P}_{j,k})} \prod_{\hat{e}\in\mathcal{P}_{m(h,i),w}} (1-s(\hat{e})) \prod_{\hat{e}\in\mathcal{P}_{m(j,k),w}} (1-s(\hat{e})) \\ &= \sum_{(h,i)\in\mathcal{L}_u^Y\times\mathcal{L}_u^Z} \sum_{\{j,k\}\subseteq\mathcal{L}_v^X} e^{-l(\mathcal{P}_{h,i})-l(\mathcal{P}_{j,k})} \left(1 - \prod_{\hat{e}\in\mathcal{P}_{m(h,i),m(j,k)}} (1-s(\hat{e}))\right) \\ &= \sum_{(h,i)\in\mathcal{L}_u^Y\times\mathcal{L}_u^Z} \sum_{\{j,k\}\subseteq\mathcal{L}_v^X} w_G(hi|jk). \end{aligned}$$

Similarly,

$$u_{XX}^+ v_{YZ}^+ - u_{XX}^- v_{YZ}^- = \sum_{\{h,i\}\subseteq\mathcal{L}_u^X} \sum_{(j,k)\in\mathcal{L}_v^Y\times\mathcal{L}_v^Z} w_G(hi|jk).$$

Notice that above cases count exactly once all quartets $hi|jk$ for all leaf nodes $h,i$ colored X, $j$ colored Y, $k$ colored Z such that MRCA of $h,i,j,k$ is $w$; namely,

$$w_{XX|YZ} = \sum_{\{h,i\}\subseteq\mathcal{L}_w^X} \sum_{j\in\mathcal{L}_w^Y} \sum_{k\in\{k':k'\in\mathcal{L}_w^Z,\text{MRCA}(h,i,j,k')=w\}} w_G(hi|jk).$$

- We define $I(G)$ to be the set of internal nodes of gene tree $G$ and $\mathcal{L}_G^X$ be the set of leaves of gene tree $G$ with color $X$. It is trivial to verify that at the

$$Q = \sum_{G\in\mathcal{G}} \sum_{w\in I(G)} w_{AA|BC} + \sum_{G\in\mathcal{G}} \sum_{w\in I(G)} w_{BB|CA} + \sum_{G\in\mathcal{G}} \sum_{w\in I(G)} w_{CC|AB}.$$

At the end of procedure `UpdateCounters`, $\sum_{w\in I(G)} w_{XX|YZ} = \sum_{\{h,i\}\subseteq\mathcal{L}_G^X} \sum_{(j,k)\in\mathcal{L}_G^Y\times\mathcal{L}_G^Z} w_G(hi|jk)$. Thus, $Q$ returned by `UpdateCounters` satisfies:

$$Q = \sum_{G\in\mathcal{G}} \left( \sum_{\substack{\{h,i\}\subseteq\mathcal{L}_G^A\\ (j,k)\in\mathcal{L}_G^B\times\mathcal{L}_G^C}} w_G(hi|jk) + \sum_{\substack{\{h,i\}\subseteq\mathcal{L}_G^B\\ (j,k)\in\mathcal{L}_G^C\times\mathcal{L}_G^A}} w_G(hi|jk) + \sum_{\substack{\{h,i\}\subseteq\mathcal{L}_G^C\\ (j,k)\in\mathcal{L}_G^A\times\mathcal{L}_G^B}} w_G(hi|jk) \right).$$

For tripartition $T = A|B|C$, note that by assumption, before the call, all the gene tree leaves are colored such that recoloring $\mathcal{L}^*$ by $X$ would produce a coloring that matches $T$. Thus, at the end of the call to `ColorLeafSet`, for each gene tree $G$, we have $A\cap\mathcal{L}_G = \mathcal{L}_G^A$, $B\cap\mathcal{L}_G = \mathcal{L}_G^B$, and $C\cap\mathcal{L}_G = \mathcal{L}_G^C$. Then, the value returned by `UpdateCounters` satisfies:

$$Q = \sum_{G\in\mathcal{G}} W(A|B|C,G). \tag{20}$$

It can be easily verified that after each call to `ColorLeafSet`$(\mathcal{L}^*, X, T, \mathcal{G}, W)$, the species tree tripartition $T$ matches the coloring of all gene trees as required by conditions of (20), concluding $W[T] = Q = \sum_{G\in\mathcal{G}} W(T,G)$. □

---

PROPOSITION 6. *Before calling* `OptimalTreeDP` *in line 6 of Algorithm S1, lookup table $W$ contains all tripartitions corresponding to internal nodes of all tree topologies in $\mathcal{S}$.*

*Proof.* Each $\hat{S} \in \mathcal{S}$ places $i$ above a different node $w$ of $S$ creating a new node corresponding to tripartition $\mathcal{L}_w|\{i\}|\mathcal{L}_S - \mathcal{L}_w$ covered in line 24. Besides new nodes, each existing internal node $w$ of $S$ will correspond to a different tripartition after placing $i$ onto $S$ depending on the relative location of $w$ and $i$. Let $u,v$ denote the larger and the smaller child of $w$. Node $w$ corresponds to $\mathcal{L}_u|\{i\}\cup\mathcal{L}_v|\mathcal{L}_S - \mathcal{L}_w$ if $i$ is under $u$, corresponds to $\{i\}\cup\mathcal{L}_u|\mathcal{L}_v|\mathcal{L}_S - \mathcal{L}_w$ if $i$ is under $v$, and corresponds to $\mathcal{L}_u|\mathcal{L}_v|\{i\}\cup\mathcal{L}_S - \mathcal{L}_w$ if $i$ is above $w$. All three cases for each node $w$ is covered in lines 20–22. □

---

*Proof of Theorem 4*

THEOREM 4. *If there exists a species tree topology $S^*$ satisfying that for each quartet subtree $ab|cd$,*

$$\sum_{G\in\mathcal{G}} w(ab|cd) > \max\left(\sum_{G\in\mathcal{G}} w(ac|bd), \sum_{G\in\mathcal{G}} w(ad|bc)\right), \tag{6}$$

*then the output of Algorithm S2 will be $S^*$.*

*Proof.* We start with a Corollary 1 of Theorem 3

COROLLARY 1. *Assuming (6), if $S$ is compatible with the true tree $S^*$, then $S'$ is compatible with $S^*$.*

By induction, $W_i$ in line 8 of Algorithm S2 should contain all tripartitions of $S^*$, as at that time $S_i = S^*$ by Corollary 1. Consequentially, the output of Algorithm S2 must also be $S^*$. □

---

*Proof of Proposition 4*

PROPOSITION 4. *The time complexity of Algorithm S2 is $O(kHn^2\log n)$.*

*Proof.* We begin by a proposition and a corollary.

PROPOSITION 7. *Procedure* `ColorNode` *on any species tree node $w$ takes $O(kH|\mathcal{L}_w|\log|\mathcal{L}_w|)$ time.*

PROOF (SKETCH) OF PROPOSITION 7. We can prove this proposition by induction. For an internal node $w$ with larger child $u$ and smaller child $v$, if for some constant $C \geq \frac{6}{\log 2}$, `ColorNode` on $u$ calls `UpdateCounters` at most $Ck|\mathcal{L}_u|(\log|\mathcal{L}_u|+1)$ times and `ColorNode` on $u$ calls `UpdateCounters` at most $Ck|\mathcal{L}_v|(\log|\mathcal{L}_v|+1)$ times, then `ColorNode` on $w$ calls `UpdateCounters` at most

$$\begin{aligned}
&Ck|\mathcal{L}_u|(\log|\mathcal{L}_u|+1) + Ck|\mathcal{L}_v|(\log|\mathcal{L}_v|+1) + 3k(|\mathcal{L}_v|+1)\\
\leq\, &Ck|\mathcal{L}_u|(\log|\mathcal{L}_w|+1) + Ck|\mathcal{L}_v|(\log\frac{|\mathcal{L}_w|}{2}+1) + 6k|\mathcal{L}_v|\\
\leq\, &Ck|\mathcal{L}_u|(\log|\mathcal{L}_w|+1) + Ck|\mathcal{L}_v|(\log|\mathcal{L}_w|+1) - Ck|\mathcal{L}_v|\log 2 + 6k|\mathcal{L}_v|\\
\leq\, &Ck|\mathcal{L}_w|(\log|\mathcal{L}_w|+1) + (6 - C\log 2)k|\mathcal{L}_v|\\
\leq\, &Ck|\mathcal{L}_w|(\log|\mathcal{L}_w|+1)\,\text{times.}
\end{aligned}$$

It is easy to verify that each `UpdateCounters` takes $O(H_G)$ time where $H_G$ is the height of the gene tree, and thus `ColorNode` on node $w$ takes $O(kH|\mathcal{L}_w|\log|\mathcal{L}_w|)$ time. □

COROLLARY 2 (Corollary of Proposition 7). *For any tree topology $S$ with $n$ species, the* `Place` *procedure on $S$ takes $O(kHn\log n)$ time.*

`NaivePlacement` of taxon set $T$ makes $r(|T|-3)$ calls to `Place`, each of which takes $O(kH|T|\log|T|)$ time. Thus, `NaivePlacement` takes $O(rkH|T|^2\log|T|)$ time and when $T=\mathcal{L}_S$ and $r=O(1)$, $O(rkH|T|^2\log|T|)=O(n^2kH\log n)$. □

---

*Proofs of Theorems 6 and Theorem 5*

THEOREM 6. *Under the conditions of Theorem 4, the DAC Algorithm S3 will output $S^*$.*

*Proof.* By Theorem 4, $S_i$ in line 5 of Algorithm S3 are compatible with $S^*$. With Corollary 1, by induction, each $S_e$ in line 21 of Algorithm S3 is compatible with $S^*$. Consequentially, $W_i$ in line 26 contain all tripartitions of $S^*$, as at that time $S'_i=S^*$, and the output of Algorithm S3 must also be $S^*$. □

THEOREM 5. *When the inequality condition in Theorem 4 is satisfied, then the time complexity of the DAC algorithm is $O(n^{1.5+\epsilon}kH)$ with arbitrarily high probability.*

PROOF (SKETCH). From the inequality (6), we can trivially deduct that $S^*$ is the species tree topology that maximizes the weighted quartet score and each $S_i$ in line 5 of Algorithm S3 is compatible to $S^*$. Also, each $C_e$ in line 15 of Algorithm S3 equals the set of species under the edges coming off of the internal nodes on the path of $S^*$ corresponding to $e$.

We now introduce a proposition

PROPOSITION 8. *With high probability,* $\max_{e\in E_{S_i}}|C_e|\le 2\sqrt{n}\log n+O(\sqrt{n})$.

*Proof.* For each pair of nodes $u,v$ of $S^*$, let $C_{u,v}:=\{x:x\in\mathcal{L}_S,u \text{ is not on } \mathcal{P}_{x,v} \text{ and } v \text{ is not on } \mathcal{P}_{x,u}\}$. It is easy to verify that for every $e$ of $S_i$, $C_e=C_{u,v}$ for some nodes $u,v$ of $S^*$. For every $u$ and $v$ that are sufficiently apart so that $C_{u,v}$ has $2\sqrt{n}\log n+\omega(\sqrt{n})$ elements and a random $T_i$ in line 4 of Algorithm S3,

$$P(C_{u,v}\cap T_i=\emptyset)=\left(1-\frac{1}{\sqrt{n}}\right)^{|C_{u,v}|}\le e^{-\frac{1}{\sqrt{n}}|C_{u,v}|}=\frac{1}{n^2}e^{-\omega(1)}=o\left(\frac{1}{n^2}\right).$$

By union bound, the probability that there exists a pair of nodes $u,v$ of $S^*$ such that $|C_{u,v}|\ge 2\sqrt{n}\log n+\omega(\sqrt{n})$ and $C_{u,v}\cap T_i=\emptyset$ is $o(1)$. Since, by definition, $C_e\cap T_i=\emptyset$ for every $C_e$, with high probability, there exists no $C_e$ having $2\sqrt{n}\log n+\omega(\sqrt{n})$ elements. □

Since $|T_i|\sim\text{Binomial}(n,\frac{1}{\sqrt{n}})$, with high probability $|T_i|=O(\sqrt{n})$ and calling `NaivePlacement` on line 5 takes $O(n^{1.5}kH\log n)$ time. It is easy to confirm that $C_\emptyset=\emptyset$ and every call to `Place` takes as input a species tree topology of $O(\sqrt{n}\log n)$ species with high probability. Thus, with high probability, each call to `Place` takes $O(\sqrt{n}kH\log^2 n\log\log n)$ time and all $O(n)$ calls to `Place` takes $O(n^{1.5}kH\log^2 n\log\log n)$ time. Therefore, the time complexity of the DAC algorithm is $O(n^{1.5}kH\log^2 n\log\log n)=O(n^{1.5+\epsilon}kH)$ with high probability. □
